# The active subset of grassland soil microbiomes changes with soil depth, water availability and prominently features predatory bacteria and episymbionts

**DOI:** 10.1101/2024.12.19.629468

**Authors:** Petar I. Penev, Katerina Estera-Molina, George M. Allen, Rohan Sachdeva, Shufei Lei, Ka Ki Law, Jordan Hoff, Steven J. Blazewicz, Jennifer Pett-Ridge, Jillian F. Banfield

## Abstract

Mediterranean grasslands, vital natural and agricultural ecosystems, experience seasonal variation in water content that likely affect microbial activity. We used metagenomics-informed stable isotope probing to investigate how the activities of microorganisms in Angelo Reserve (2160 mm rainfall) and Hopland (956 mm rainfall) soil change over depth and the seasons. At both sites, we find that the relative abundances of organisms in shallow soil changes relatively little but the most abundant organisms vary greatly with soil depth. Notably the highest levels of isotope incorporation, indicative of growth, occurs in deep soils. The active part of the 0-10 cm soil community varies over time, especially in Hopland soils during the fall rewetting. We defined a large, novel clade of Actinomycetota with notable capacity for thiosulfate oxidation whose representatives are prevalent and active in deep soils (>20 cm) across both ecosystems. Active Saccharibacteria unexpectedly encode nucleotide synthesis genes that enabled isotope incorporation while growing in shallow Angelo soils over all time periods. In contrast to predicted episymbiotic lifestyles of Saccharibacteria, other highly active bacteria are predicted predators. Obligately predatory Pseudobdellovibrio are active in intermediate depth Hopland soils whereas bacteria of the order Haliangiales are active in shallow Angelo soils. Supporting predatory lifestyles of Haliangiales, we used *in silico* structure prediction to assemble a large protein complex that we identify as a contractile injection system. Overall, the results indicate the potential for active carbon turnover in deep grassland soil and strong seasonal changes in the active members of microbial communities, despite relatively minor shifts in community composition.

## Introduction

Soil is the planet’s second largest active pool of carbon and it has been under direct anthropogenic influence for thousands of years. One of the terrestrial ecosystems with major importance to humanity are grasslands, since they cover approximately 40% of the earth’s surface and 80% of agriculturally active land over a wide range of geological and climatic conditions [1]. Grasslands overlay about a quarter of the terrestrial area and store about a third of the carbon locked on land [2, 3]. Microorganisms are some of the main determinants of carbon persistence and turnover within grassland soils. They consume plant and microbially derived polymers and incorporate carbon and nitrogen in proteins, lipids, nucleic acids, polysaccharides, pigments, and other metabolites. Microorganisms change their environments via biofilm formation, predation, extracellular enzyme production, and other secreted polymers. Cells interact with minerals and their carbon and nitrogen compounds may be sequestered by adsorption to mineral surfaces [4–6].

Environmental moisture content is a major controller of microbial behavior, due to the critical importance of water in cellular biochemistry [7], as a controller of microbial activity due to fluctuating availability of nutrients [8], and as a medium for intercellular communication [9]. With the changing climate, soil moisture variations will become more extreme [10] and the importance of water availability to microbial dynamics will increase [11–14].

In mediterranean climates that feature a long dry summer period, water availability changes dramatically over the seasons. During the spring, plants provide plentiful resources and microbes flourish. Through the interaction with plant roots, microbial and virus actors move plant-derived carbon into CO_2_, dissolved organic C, biomass and mineral-associated organic matter [15, 16]. Microbes employ a rich variety of traits that allow them to persist during periods of rapid environmental change [17, 18]. During the dry down microbes actively accumulate and synthesize compatible solutes to protect themselves from osmotic stress, secrete water-retaining extracellular polymeric substances, modify their membranes to maintain fluidity [19], use trehalose and control their ion levels to prevent protein damage [20, 21], employ DNA repair and heat shock proteins [22, 23], and finally undergo sporulation [19, 24]. After the first rains of the fall season, increased metabolic and mineralization activity [25–28], can lead to the consumption of soil carbon in minutes [26, 29, 30]. This period of time is also linked to considerable bacterial and fungal mortality [31, 32] due to osmotic stress [33] and possibly predation.

Deep soils comprise the majority of the global soil carbon pool [34] and have large potential to sequester carbon [35]. Microbial types and traits change with soil depth [36–38] and deep soils (>30 cm) are thought to be linked to lower microbial activity compared to shallow soils. Important metabolic traits are associated with deep soil microorganisms, such as ammonia oxidation, “dark autotrophy”, CO_2_ fixation, Wood-Ljungdahl pathway [39, 40]. Carbon dynamics in deep soils respond to environmental changes [41–43] there are important knowledge gaps regarding which microbes are active in these regions, and at what levels.

Quantitative stable isotope probing (qSIP) allows the identification of active community members through the detection of newly-synthesized DNA molecules that are enriched with a stable isotope [15, 44]. SIP methods have shown that only a fraction of soil organisms are active at a given place and time [32, 45–47]. Here, we applied H_2_^18^O qSIP-informed metagenomics to analyze soils sampled at three depths from two Mediterranean climate grassland ecosystems. Unlike amplicon-based SIP, this approach extensively samples the genomes of soil organisms, enabling linkage between active organisms and the capacities that define their lifestyles and potential ecosystem impacts. Additionally, using H_2_^18^O for isotope delivery ensures that enrichment is not dependent on the microorganisms feeding strategy. The study captured the three most important periods for the water year: spring peak plant productivity, summer dry down, and first fall wet up. We estimated bacterial and archaeal taxon-specific activities and connected them to traits of importance, and used soil physicochemical measurements to provide context. Overall, the results clarify how microbial community composition and activity change with depth and time over periods of dramatic variation in water availability in grassland soil ecosystems.

## Results

### Soil sampling, SIP incubation, and assembly

In order to investigate how the soil microbiomes, and particularly the active members of soil microbiomes, change over the “water year”, with soil depth and annual rainfall input, we collected 24 cores of soil from each of the Angelo and Hopland field sites. At each site, at eight time points we collected three replicate cores. Sampling began April 6th 2022, subsequent samples were collected throughout the spring season and summer seasons every ∼2 weeks, until the soils reached their driest point, and finally after the first fall rain event. Of particular interest were changes in microbiome structure and activity before and after the first rain event following the long dry season. At Hopland we were able to sample 24 hours before and 48 hours after rain, in September, 2022. For Angelo, the first rainfall was not predicted with sufficient time to enable sampling beforehand, but samples were collected 17 hours after the first rain.

From each core, we extracted samples from five depth intervals between the surface and 80 cm. For all 48 samples, we measured soil moisture, the total C/N ratio, pH (**Supp. Fig. 1, Supp. Data 1**), and for one time point at three depths - moisture retention curves (see Methods; **Supp. Fig. 2**). Angelo soils had on average 5-10% more moisture, compared to Hopland soils (**Supp. Fig. 1**). Shallow soils experience large variation in moisture content over the seasons with much wetter shallow soil during the rainy season. In contrast, the moisture content of deep soils (50-80 cm), well below the observed rooting depth of ∼30 cm, was relatively constant (**Supp. Fig. 1**).

To obtain a quantitative measure of microbial activity, we incubated all samples with natural abundance water and isotopically heavy water (H_2_^18^O). Additionally, we measured the CO_2_ flux from each sample over a seven day period as an indication of total microbial respiration. General linear model showed a positive relationship between field moisture content and CO_2_ rate for samples at 0-10 cm and no strong signal for deeper soils (**Supp. Fig. 3**). Assuming water must be taken up during growth for synthesis of DNA, we tracked incorporation of ^18^O from water into DNA as an indicator of the activity of any given species. The more active a species is, the denser the DNA molecules. Thus, metagenomic DNA from three depths and five timepoints was fractionated by density and atom fraction excess (AFE) values for sequences encoding marker genes was calculated [44, 48]. We found that activity of organisms varies over time and depth, with highest AFE values of 0.7 (**Fig. 2**).

Enough DNA for fractionation and sequencing was recovered from all but one sample (**Supp. Data 2**). In total we sequenced 89 biological unfractionated samples twice, once without a SIP incubation (T0) and another time with H_2_^16^O incubation (^16^O). We also sequenced 890 fractionated samples (five for each isotope for each biological sample). In total, we generated almost 5 Terabase pairs of raw read data. The data were assembled into 545 Gigabase pairs (Gbp) contiguous sequences, of which 1.5% was in contigs longer than 50 Kilobase pairs (Kbp). Metagenomic binning produced 4493 unique bins with completeness higher than 75% and contamination lower than 25%, determined by single copy gene (SCG) analysis from dRep [49].

### Community composition

To gain a broad understanding of the community composition we classified organisms at approximately the species level based on the sequences of the genes encoding ribosomal protein S3 (rpS3) [50]. After clustering the sequences from both ecosystems at 99% nucleotide identity, we generated 52,680 rpS3 species-level clusters, labeled as species groups (SGs). Of those, 9441 SG sequences were found in bins (18% of all SGs), accounting for 47 ± 9% of the reads that map to these genes.

We used the co-occurrence of rpS3 sequences in SGs in a network analysis to evaluate microbiome compositional shifts over time, replicates, and soil depths (**Fig. 1 A)**. Spatial variables such as ecosystem (Hopland vs. Angelo), depth, and replicate demonstrate strong influence on the community composition. Samples from shallow soils show smaller centrality values, indicating few rpS3 sequences shared with deeper soils. This is indicative of low species similarity between shallow and deep soils.

**Figure 1:**
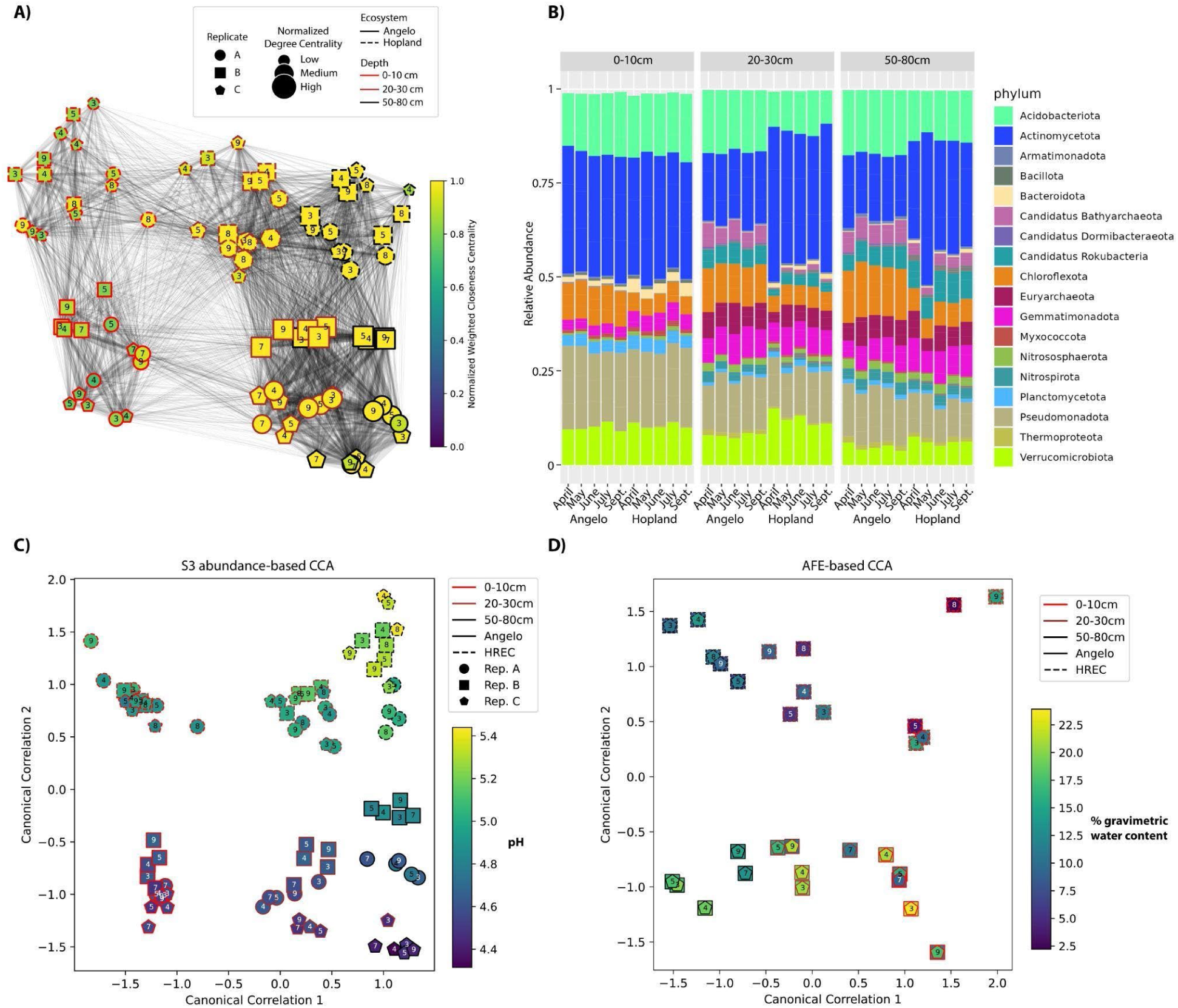
Relative abundance based on rpS3 species groups (SGs), influence of spatio-temporal variables, and atom fraction excess (AFE) Canonical Correspondence Analysis (CCA). A) Network analysis, based on rpS3 SG sequence identity clusters at 99% identity, where each sample is represented as a node and an edge is drawn when there is a cluster with SGs from two samples. The weight of edges is shown with the intensity of the drawn line and corresponds to the number of SGs shared between samples across all clusters. Nodes are colored by normalized weighted closeness centrality and their size corresponds to normalized degree centrality. Shapes of nodes indicate replicates, edge colors distinguish between depths, and dashed or solid edge lines indicate Hopland or Angelo. Node labels indicate sampling time. B) Relative abundance values calculated from mapping all sample reads to rpS3 contigs, assembled from the respective sample. C) CCA calculated using total rpS3 SG abundances and D) using atom fraction excess values including sample metadata (Water content, pH, total C/N, CO_2_ rate, ng DNA extracted per gram soil). CCA was applied on the species groups explaining 80% of the variance, determined with a PCA ordination (**Supp. Fig. 13**). Water content and pH for each sample are plotted respectively on D) and C).

Abundance values were calculated for each SG in each sample, using the mapping depth of time zero (T0) reads to contigs containing the rpS3 gene. Rank abundance curves reveal lower metagenomic diversity in deeper samples compared to shallow ones across all timepoints, with more organisms at relatively high abundance compared to shallow soils (**Supp. Fig. 4**). Conversely, shallow soils contain a few highly abundant organisms, with the remainder of organisms at similarly low abundance levels.

Interestingly, qSIP data indicate that highly abundant organisms in shallow soils are generally inactive (**Supp. Fig. 4**). Highly active species are found throughout the entire abundance curve distribution including the tail end (**Supp. Fig. 5,6**), meaning that even species with low metagenomic abundances have high levels of replication and can influence the soil carbon turnover at all soil depths.

The 100 most abundant organisms in 0-10 cm soil are similar over time, especially in Angelo soils (**Supp. Fig. 7-12**). In contrast, the 100 most abundant organisms in deep soil are quite variable over time. Furthermore, the organisms present at highest abundances in 0-10 cm soil are very different from those most abundant in deeper soils (**Supp. Fig. 7-12**).

The soil phylum level microbial diversity is generally in agreement with previous studies of the Angelo and Hopland sites [47], and shows little change over time (**Fig. 1 B**). Actinomycetota dominate all samples, followed by Acidobacteriota, Pseudomonadota, and Verrucomicrobiota. As previously observed at the Angelo [38, 39]broadly across the US [39], archaeal relative abundances increase with soil depth.

Taxonomic distributions observed through rpS3 abundance-based Canonical Correspondence Analysis (CCA) match the distributions established with network analysis (**Fig. 1 A**) and closely track pH changes (**Fig. 1 C**). Thus, we evaluated patterns of diversity at the two sites as revealed by the CCA analysis. The taxonomic diversity for deep compared to shallow samples is greater across canonical correlation 2 of the CCA due to the lower similarity among replicates in the deep compared to shallow samples (**Fig. 1 C, Supp. Fig. 7-12**). However, when using activity calculated from SIP data in the CCA ordination analysis, active organisms in the shallow samples are more diverse compared to those from deep samples (**Fig. 1 D**). We conclude that deep soils have a broad taxonomic diversity but a constrained set of active species throughout the year at both sites. The reverse is true for shallow soils, where the diversity of active species is greater than in deep soils at all different timepoints, especially in Hopland (**Fig. 1 D**).

While the network, taxonomic, rank, and rpS3 abundance-based CCA analysis were not able to determine any effect of the time component, the SIP activity-based CCA reveals a distinctive microbial behavior over time (**Fig. 1 D**). The organisms in shallow Hopland soil that grew in response to laboratory wet up are similar for samples collected before and after the first Fall rain event and very different to those that grew after laboratory wet up of soils from time points corresponding to spring peak plant productivity and summer dry-down.

### Seasonal dynamics in shallow soils

To understand the seasonal dynamics of shallow soil organisms we clustered SGs based on AFE for each timepoint, creating cohorts of seasonal activity (**Fig. 3**). Cohorts active <u>only</u> during peak plant productivity (May) have few representatives and have the lowest activities of any cohort, both in Angelo and Hopland (**Supp. Fig. 16**). The May-only cohort in Angelo contains representatives exclusively from the Actinomycetota phylum. Most organisms active during peak plant productivity are also active at other timepoints so they are not in this cohort. Bacteroidota show highest levels of activity during April and May (around the time of peak plant growth), and September (**Fig. 2,3**). Very few other organisms show the same temporal activity, so Bacteroidota were placed in a general cohort that includes organisms active at various time points (“Various” cohort, **Fig. 3**).

**Figure 2:**
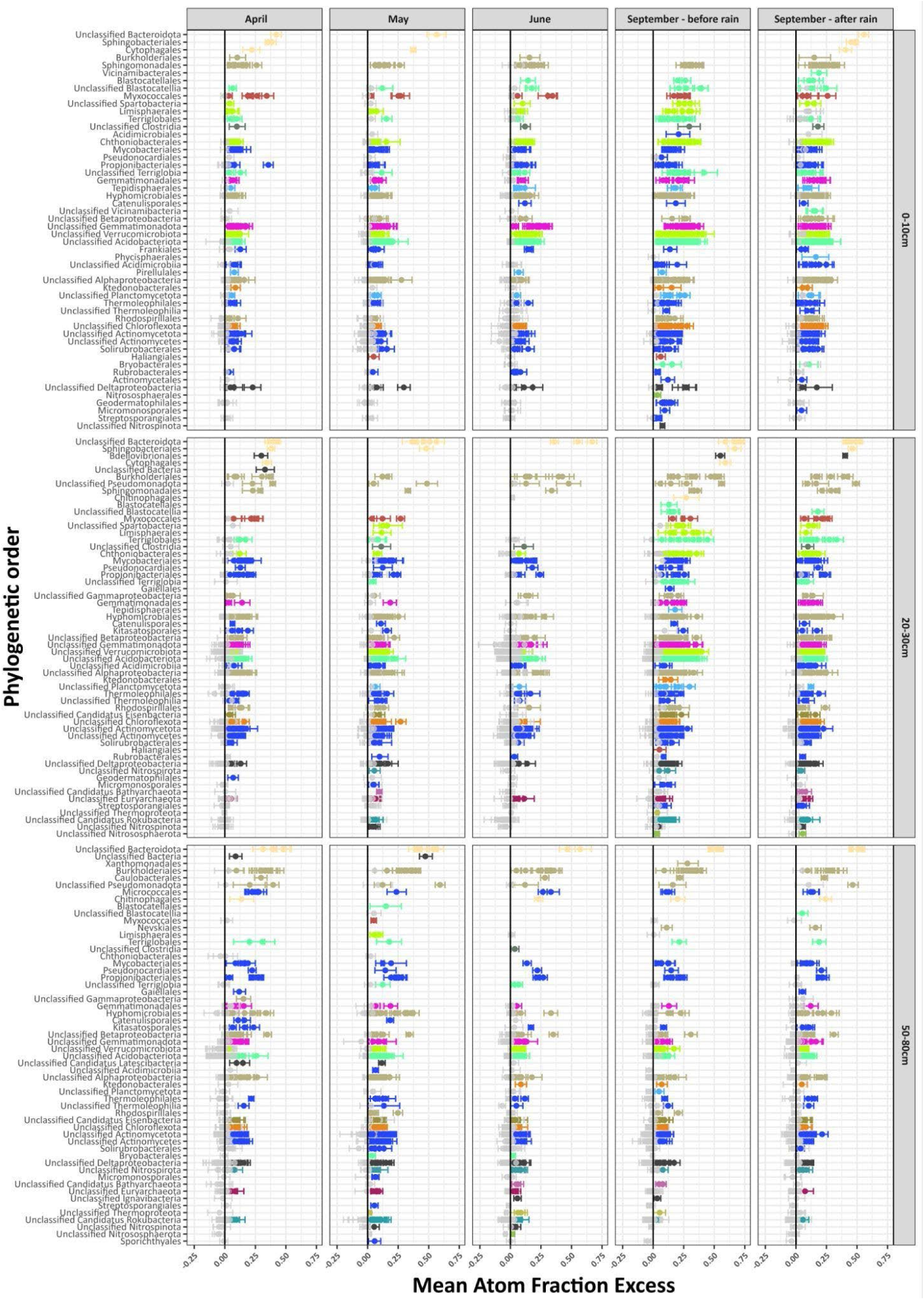
Atom fraction excess results from Hopland samples. Each biological sample is represented as a subplot on the figure with five timepoints as columns and three depths as rows. rpS3 SGs with detected isotope excess are organized by phylogenetic order and are arranged in descending AFE values. Each phylogenetic order can have multiple rpS3 contigs and are colored by phylum as in Figure 1 when the lower confidence interval on the AFE calculation is above zero. rpS3 SGs with confidence interval overlapping zero are colored gray. Error bars are 90% confidence intervals based on a bootstrapping procedure of 1000 resamples of the weighted average densities based on three replicates.

**Figure 3:**
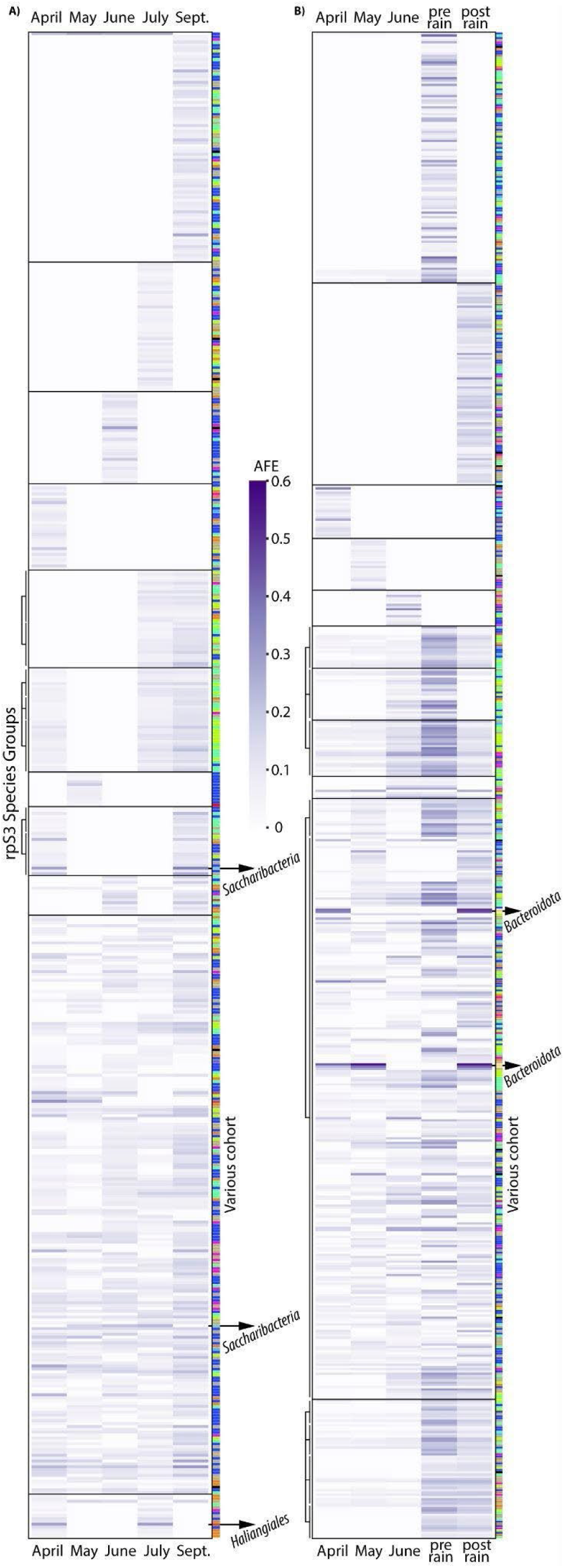
Seasonal cohorts of organisms, based on AFE results from 0-10 cm soils for A) Angelo and B) Hopland. The AFE of each organism (represented by rpS3 species group) for each timepoint is shown with a purple gradient. Cohorts are shown with black boxes and are based on clusters, defined by the similarity of the AFE activity for each organism pair (**Supp. Fig. 14,15**). Cohorts can include single or multiple clusters. When there are multiple clusters, they are indicated by brackets on the left side of the plots. Merging multiple clusters in a cohort was determined manually. The phylum of each organism is indicated with a colored line on the right side of each plot and the colors match those in Fig. 1. The cohort that includes species with various activities is indicated with “Various cohort” on the right side of the plots. The last two timepoints for Hopland (“pre rain” and “post rain”) are in September.

There were two cohorts that were active either before or after the fall rain event (top two clusters in **Fig. 3**). These cohorts are large and diverse, including many representatives from Acidobacteriota, Actinomycetota, Chloroflexota, and Verrucomicrobiota. Interestingly, when examining cohorts of organisms active both before and after the fall rain event in Angelo (**Fig. 3 A**) there is a distinct lack of Actinomycetota representatives and larger proportion of Acidobacteriota and Verrucomicrobiota (**Fig. 3 A**).

A Saccharibacteria, known to have epibiotic lifestyle [51], is active only in April and September and clusters with active representatives spanning the Chloroflexota, Gemmatimonadota, Pseudomonadota, Verrucomicrobiota, and the Actinomycetota phyla (**Fig. 3**). Actinomycetota are of particular interest as possible hosts, given prior studies showing these bacteria are Saccharibacteria hosts [51]. The Actinomycetota include a very diverse set of organisms that show patterns of activity in Angelo and Hopland leading to their placement in the Various cohort and in cohorts that are active at specific timepoints (**Fig. 3**, **Supp. Fig. 16**). Another Saccharibacteria species is active only in May, June, and July, but there are no organisms with similar activity patterns. The lack of correlated active species obfuscates candidate hosts for this putative episymbiont.

We sought functions that are characteristic of, or shared by, cohorts. Processing of starch and chitin are among the mostly highly represented functions in all cohorts at both sites (**Supp. Fig. 17**). Also highly prevalent at both sites are genes for processing a diverse range of complex carbohydrates (e.g., xylans, amorphous cellulose, xyloglucan, beta-mannan, and polyphenolic compounds) (**Supp. Fig. 17**). Also highly represented in all cohorts from both sites are genes for acetate kinase, acetyl CoA synthase, and pyruvate oxidation, key enzymes in central carbon metabolism (**Supp. Fig. 17**). Notably, ≥ 91% of genomes from organisms in all Angelo cohorts, except the one active only in April (74%), have genes for metabolism of starch that may derive from degradation of plant material in late Spring. Starch metabolism is less prevalent in Hopland at some time points, including in the April cohort. Chitin degradation genes are prevalent in most Angelo and Hopland cohorts, with less representation in the May cohort from Angelo (**Supp. Fig. 17**).

Interestingly, the Angelo and Hopland cohorts comprising organisms active only in May show the largest difference in functions compared to other cohorts (**Supp. Fig. 17**). In Angelo, half of the organisms from the May cohort contain nitrite and arsenate reductases. At Hopland, 25% of the genomes of organisms from this cohort have genes involved in interconverting nitrate and nitrite, compared to no more than 13% of genomes in any other cohorts (**Supp. Fig. 17**). Thus, we infer that nitrate/nitrite metabolism is a feature of bacteria only active in May at both sites.

Approximately half of the organisms active only in June in both Angelo and Hopland soils have the capacity to oxidize thiosulfate. This function is only present in 0 - 28 % and 12 - 38 % of genomes of organisms in other cohorts at Angelo and Hopland, respectively.

### Stable isotope incorporation reveals highly active Saccharibacteria

Some of the organisms with highest activity in shallow soil from the Angelo site are two unclassified Saccharibacteria - one active in April and September, the other during the dry down period of May, June, and July (**Fig. 4**). These are Candidate Phyla Radiation (CPR) bacteria [52], which are known to have limited biosynthetic capabilities and have been shown in some cases to live as episymbionts [53]. Saccharibacteria described previously have reduced nucleotide synthesis capabilities [54], however the Saccharibacteria we found active throughout the wet-up and plant productivity periods (April and September) has complete pathways for purine and pyrimidine synthesis (**Supp. Fig. 18**). The Saccharibacteria active in the dry down period (May, June, and July) has many more genes for nucleotide biosynthesis than most Saccharibacteria described from a wide diversity of environments (**Fig. 5**). This finding extends the variety of metabolic capacities known for Saccharibacteria (**Fig. 5**, **Supp. Fig. 19**) and provides evidence for direct isotope incorporation during growth, rather than reliance on recycled nucleotides.

**Figure 4:**
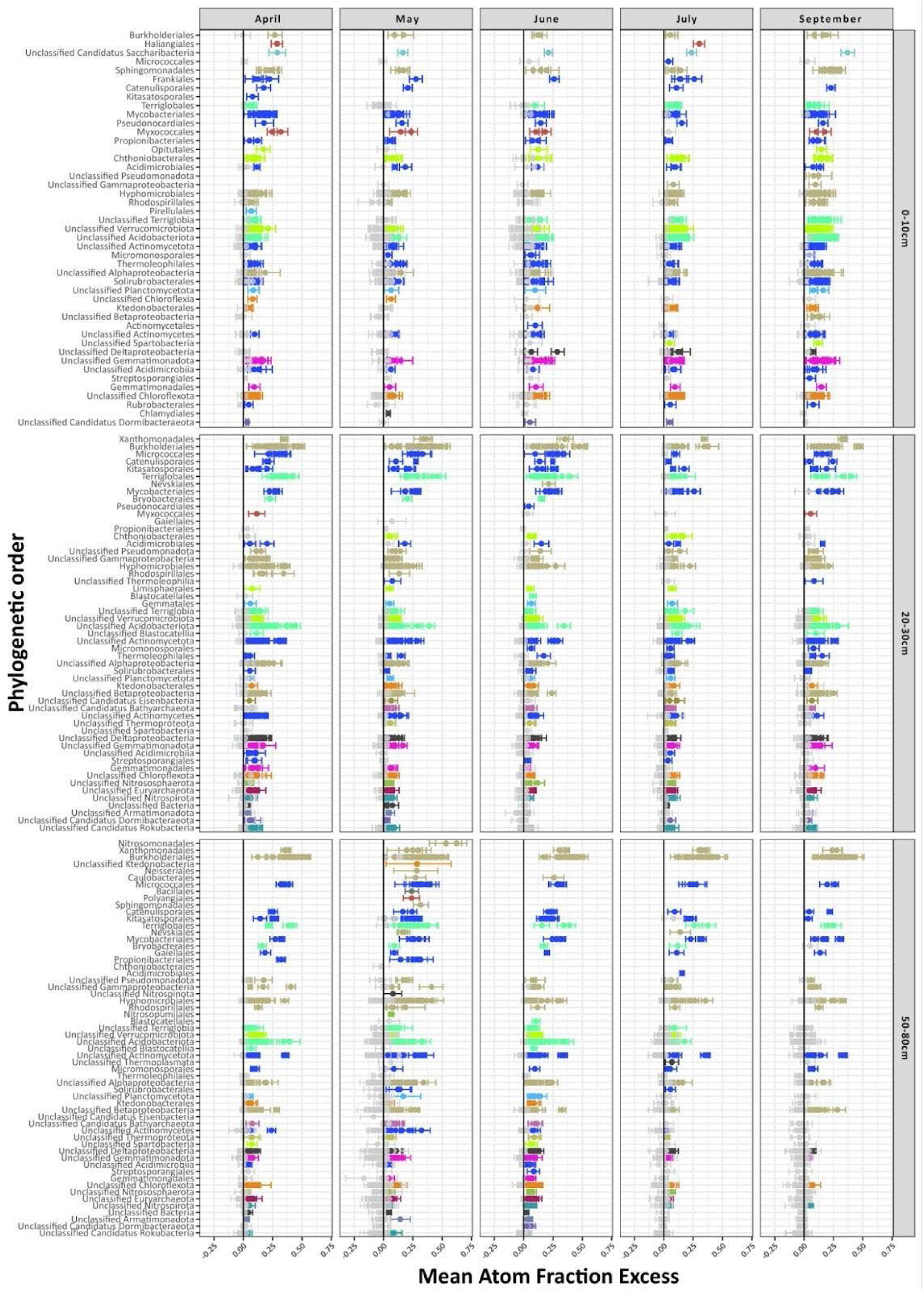
Atom fraction excess results from Angelo samples. Each biological sample is represented as a subplot on the figure with different timepoints as columns and different depths as rows. rpS3 SGs with detected isotope excess are organized by phylogenetic order and are arranged in descending AFE values. Each phylogenetic order can have multiple rpS3 contigs and are colored by phylum as in Figure 1 when the lower confidence interval on the AFE calculation is above zero. rpS3 SGs with confidence interval overlapping zero are colored gray. Error bars are calculated as described in Figure 2.

**Figure 5:**
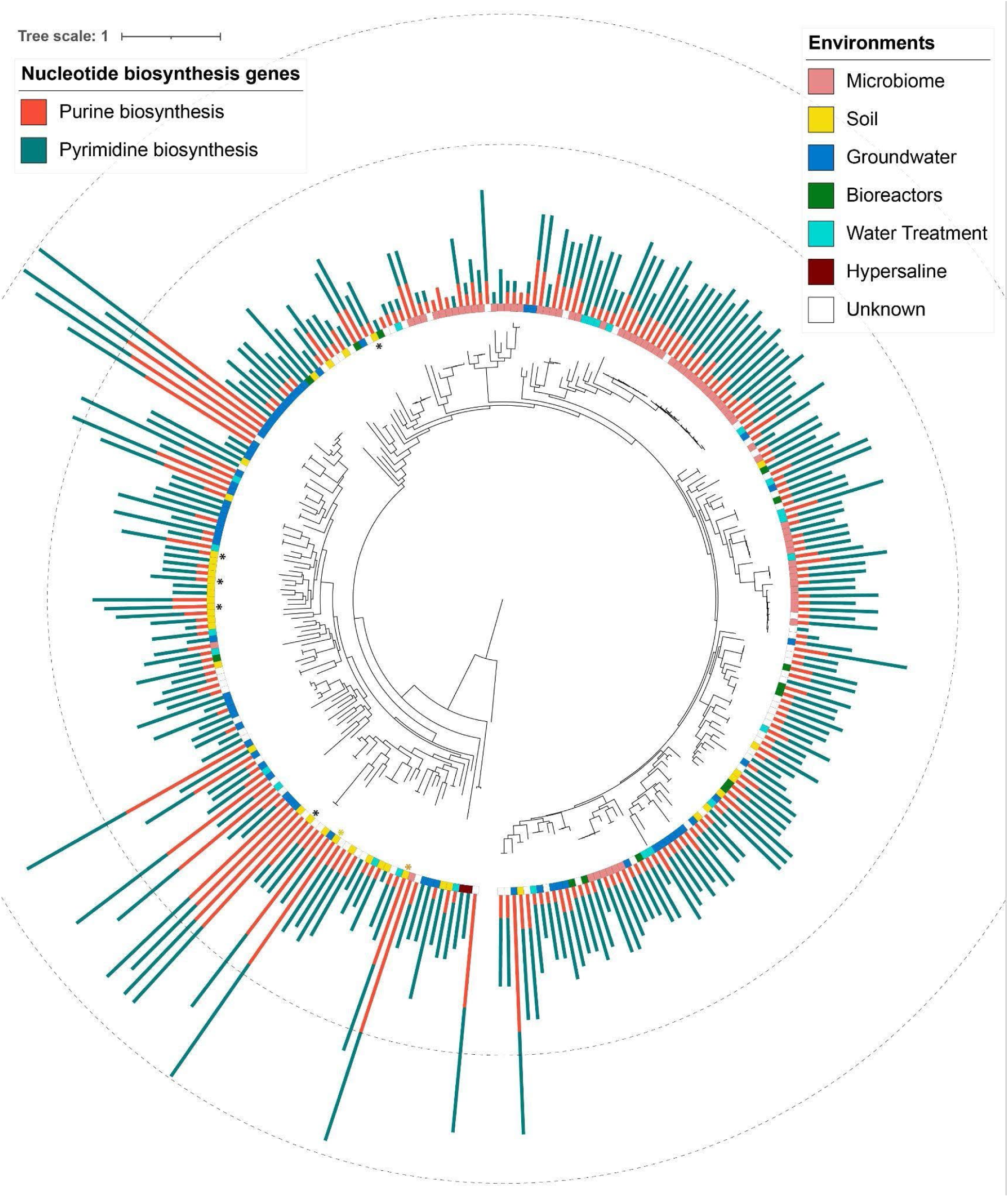
Phylogenetic tree of rpS3 sequences from this project and publicly available Saccharibacteria metagenomes. Environmental source is indicated with a color strip and retrieved from [67] and ggKbase. Non-redundant count of nucleotide biosynthesis genes, present in each metagenome, are shown as bars. The number of genes present in the complete pathways for Purine and Pyrimidine synthesis are shown as two concentric dashed lines. Asterisks indicate metagenomes from this study. Orange asterisk indicates the Saccharibacteria active in May, June, and July, while the yellow asterisk is the one active in April and September. Phylogenetic tree with labeled metagenomes, bootstrap values, and percent completeness of the nucleotide synthesis pathways is available in the supplementary materials (**Supp. Fig. 19**).

The active Saccharibacteria contain genes for F-type ATPase and ubiquinol oxidase, suggesting the capacity of aerobic respiration (previously reported in Saccharibacteria [54, 55]). The Saccharibacteria active throughout the dry-down period also contains genes for an inorganic carbon pump [56]. These are used to concentrate CO_2_ (as bicarbonate) for improved function of RuBisCo [57], yet Saccharibacteria lack RuBisCo. Bicarbonate can control toxin expression in the parasitic *V. cholera* and *B. anthracis* [58, 59], so those genes may control the epibiotic cycle of Saccharibacteria.

### Bacteroidota species from Hopland soils are highly active at all times and depths

Bacteroidota were some of the most active organisms across all timepoints and depths (**Fig. 2**), except the driest timepoints in shallow soils. Classified examples fall in the Sphingobacteriales and Cytophagales orders. Cytophagales are cellulolytic bacteria involved in organic matter turnover [60] and remineralization of organic materials into micronutrients [61]. Multiple other Bacteroidota are unclassified. The functional annotation of the Bacteroidota MAGs show a rich variety of CAZy enzymes in support of roles in carbon turnover and remineralization (Supp. Data 3). All active Bacteroidota in our study have CAZy capabilities for processing arabinose, beta-mannan, chitin, mixed-linkage glucans, starch, xylans, and xyloglucan (**Supp. Fig. 20**).

### Deep soils harbor highly active organisms and a nucleocytoplasmic large DNA virus

Unexpectedly, stable isotope incubations of deep soils (50-80 cm) show relatively high levels of activity, sometimes with the highest AFE values in our dataset (**Fig. 2,3**). Some of the most active SG in both Angelo and Hopland soils during all timepoints are from Actinomycetota (Microccales), Pseudomonadota (Xanthomonadales, Burkholderiales), and Acidobacteriota (Terriglobales). The most active organism in Angelo deep soils during the May sampling (**Fig. 4**) was *Methylobacillus flagellatus.* The MAG for this organism contains a PQQ-dependent dehydrogenase from the methanol/ethanol family; this is consistent with previous results identifying *M. flagellatus* as a methylotrophic aerobic bacteria involved in carbon recycling of methane and methanol [62]. Among the most active organisms within Hopland deep soils are members of the Oxalobacteraceae family, the most active MAG of which contains methanol and carbon-monoxide dehydrogenases.

Intriguingly, we identified a giant eukaryotic virus, a member of the *Nucleocytoviricota* [63], in a sample from Hopland 50-80 cm deep soil sampled in June. The draft genome has a length of 381 Kbp (8 contigs) and extremely low GC content of ∼20%. While its DNA was only found in a single replicate, reads from other timepoints map to the genome (**Supp. Fig. 21**). Most of the DNA comes from ^18^O samples and comparing the amounts from different fractions, indicates this virus was actively replicating (**Supp. Fig. 21**). The most common taxonomic affiliations for the NCLDV proteins are to Pythium and Phytophtora, oomycetes (fungus-like eukaryotes with saprophytic or pathogenic lifestyles) that are often plant pathogens. We identified a 31 kb of mitochondrial genome at ∼⅕ of the coverage of the NCLDV and similar atom enrichment, indicating likely activity (**Supp. Fig. 21**). The fungal genome was classified taxonomically as a member of the *Pythium* genus, indicating it as a possible host for the virus.

### Low AFE organisms with important metabolic potentials for carbon and nitrogen cycling

Less active organisms that have a detectable level of isotope incorporation likely also contribute to carbon and nitrogen turnover. For example, Nitrososphaerota have detectable activity in the 20-30 cm and 50-80 cm soils at almost all timepoints in Angelo and in Hopland at several timepoints (**Fig. 2, 3**). These archaea fix CO_2_ and perform ammonia oxidation [64, 65]. Of these, Nitrosopumilales are active in the deepest Angelo soils, but only at one time point (**Fig. 4**). Other active archaea include multiple members of the Thermopasmata phylum (**Supp. Fig. 22**), which have previously been observed to contain copper membrane monooxygenases with a role in aerobic oxidation of ammonia [66].

Bacteria with low but detectable activities included Planctomycetota (active across all timepoints and depths in Angelo and 0-10, and 20-30 cm soils in Hopland, **Fig. 2,3**), Nitrospirota (in 20-30 and 50-80 cm soils of both ecosystems), and Nitrospinota (active in some Hopland soils, **Fig. 2,3**). All of these bacteria have capabilities for complex carbon degradation, nitrification, denitrification, and anaerobic ammonium oxidation. Overall, the moderately active archaea and bacteria in deeper soils are implicated in autotrophy, oxidation of ammonia, and carbon turnover.

### Microbial activity in drier Hopland vs. wetter Angelo soils

Given the substantial difference in timing and amount of rainfall between Hopland and Angelo (**Supp. Fig. 1**) we investigated differences in the activity of taxa shared between the two ecosystems. Of the 652 SGs detected in both Angelo and in Hopland, 496 show activity at any time (**Supp. Fig. 23**). During the wet-up timepoints in 0-10 cm soils Angelo always stayed above 10% moisture levels (**Supp. Fig. 1**) and we observed greater activity of Hopland compared to Angelo microorganisms (**Supp. Fig. 23**). SGs from the Verrucomicrobia and Acidobacteria display the largest differential in activity, with much greater isotopic incorporation observed at Hopland compared to Angelo (**Supp. Fig. 23**), especially for 0-10 cm and 20-30 cm soil. During the driest sampling points, the SG representatives from Hopland are more active than the related SG at Angelo, correlated with the lower moisture content of shallow Hopland soils.

There are 1442 SGs that are active and unique to Angelo, compared to 1676 from Hopland (**Supp. Fig. 24**). At the taxonomic order level, the distribution of activities of a large group of organisms (middle group on **Supp. Fig. 24**) is similar at both sites. However, within the upper and lower order-level groups in **Supp. Fig. 24** there are different SG active at the two sites. For example, Blastocatellia (Acidobacteriota) and Sphingomonadales (Proteobacteria) are more active in shallow soils at Hopland (upper group on **Supp. Fig. 24**), whereas Haliangiales (Myxococcota) and Saccharibacteria (CPR) are more active at Angelo (lower group on **Supp. Fig. 24**). In deeper soils Propionibacteriales (Actinobacteriota) and Eisenbacteria are more active in Hopland, while Burkholderiales (Proteobacteria), Ktedenobacteria (Chloroflexi), and Micrococcales (Actinobacteriota) are more active in Angelo. Overall, the distribution of AFE values shows a skew of -0.31 when Angelo taxa are assigned negative AFE. This implies that microbial activity in deep Angelo soils is higher than in deep Hopland soils.

### Large undescribed and active Actinomycetota clades

Actinomycetota, which is the most highly represented phylum based on relative abundance values (**Fig. 1B**), also has widely varying activity across timepoints, ecosystems, and depths (**Fig. 3,4**). More than 12 distinct orders of Actinomycetota show generally similar levels of activity between Angelo and Hopland in shallow soil (**Fig. 2, 3**). To gain further understanding of the Actinomycetota ecological makeup we constructed a phylogenetic tree from Actinomycetota sequences that showed activity (**Fig. 6**) and included reference sequences from previous research on the Tree of Life [68] and all Actinomycetota annotated S3 sequences from NCBI, clustered at 99% identity (**Supp. Fig. 25**). The tree includes 892 Actinomycetota sequences from our project and 542 reference sequences.The phylogenetic tree uncovers two large clades that are novel, with the exception of some NCBI sequences classified only at the phylum level (outer ring in **Supp. Fig. 25**) and some that we previously reported from Angelo [38]. One of these forms a class level clade (**Supp. Fig. 25**) and the other nests within the Rubrobacteridae class. The diversity with the Rubrobacteridae clade is greatly expanded by our data (**Fig. 6**). Generally, species from those clades are most active in deeper (> 20 cm) Hopland soil, but there is detectable activity in some shallow soils and at Angelo (**Fig. 6**). Compared to other classes in Actinomycetota, the metabolically relevant functions of the novel clades are enriched in thiosulfate oxidation capabilities by SOX complex (**Supp. Fig. 26-29**, **Supp. Data 4**). Two representatives from the novel clade 1 are found in the June seasonal cohort for Angelo and one of them is found in Hopland as well (**Fig. 3**). There is a higher percentage of MAGs containing thiosulfate oxidation capabilities in the June cohorts (**Supp. Fig. 17**). The most active Actinomycetota are species in the Actinobacteridae class, which are present both in Angelo and Hopland, most often in deeper soils (> 20 cm), but not absent from the shallow layer (**Fig. 6**). The strongest activity of the novel clades in Hopland 0-10 and 20-30 cm soils is centered around timepoints that involve rewetting the soil, right before and after the first fall rain event.

**Figure 6:**
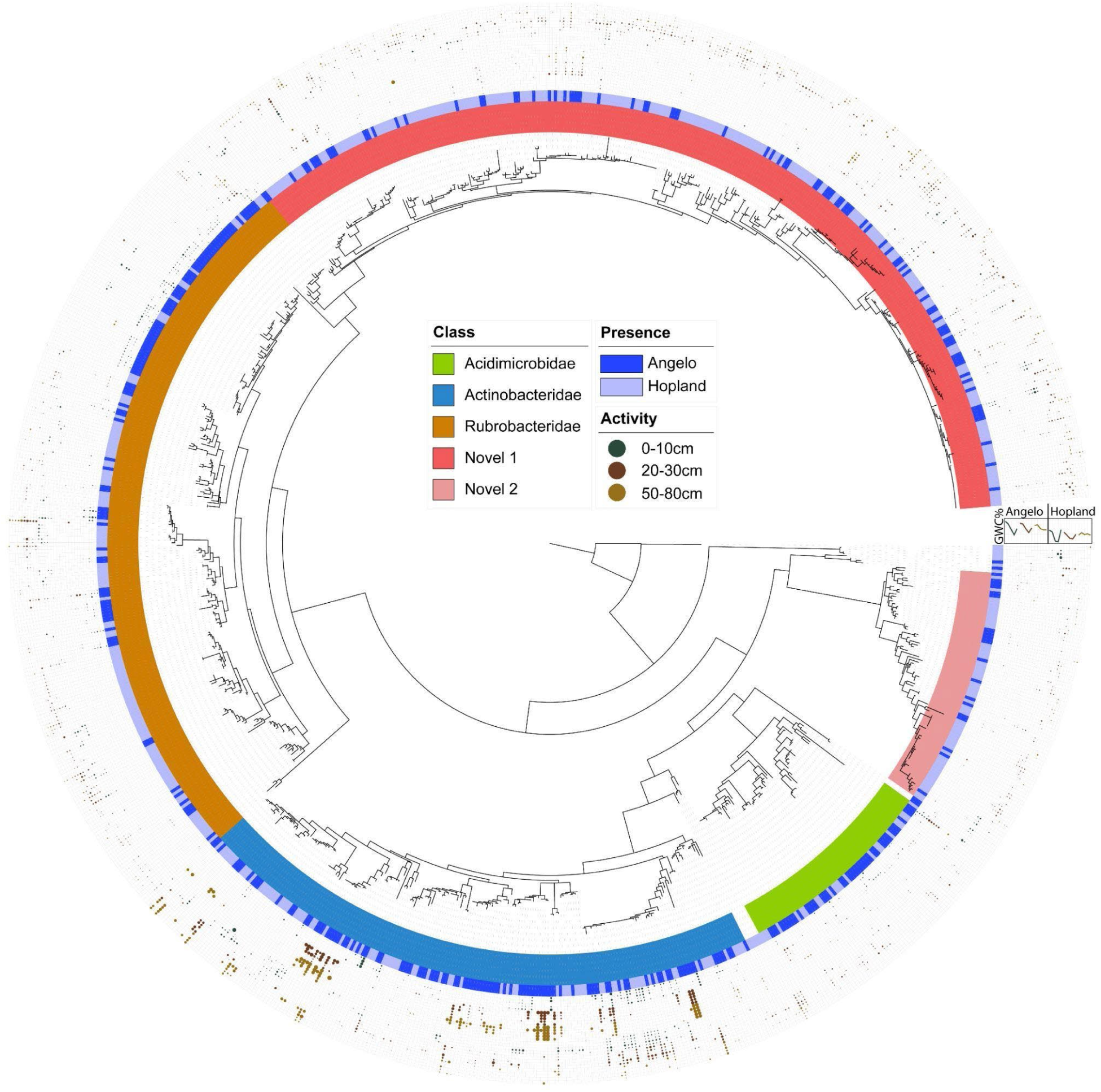
Phylogenetic tree of active Actinomycetotafound during the Water Year. The tree includes 892 Actinomycetota rpS3 sequences, showing AFE values above 0.05. Class level clades are defined based on a phylogenetic tree including Actinomycetota reference sequences (**Supp. Fig. 25**). Stripe label indicates the ecosystem source of the sequence (Angelo or Hopland). Circles organized in concentric rings indicate the activity of each sequence with their size, having larger size indicating greater AFE. The circles are colored by depth and organized by ecosystem, depth, and time. AFE values from Angelo are on the inner rings and Hopland rings are outward. Timepoints repeat for each depth starting with April being inner and ending with September on the outer rings. Gravimetric water content is indicated on the right side, on top of the AFE rings.

### Predatory bacteria are some of the most active organisms

Species from the order Haliangiales in the Myxomycota phylum show high activity in Angelo shallow soils and moderate activity in middle depth soils (**Fig. 4**), yet they comprise < 1% relative abundance of the microbiomes (**Fig. 1 B**). This difference in activity vs. abundance underlines the importance of identifying active taxa with SIP. Sequences from the Haliangiales order show strong activity in April and July, coinciding with activity of Frankiales, Alphaproteobacteria, and Chloroflexota species. Various representatives from Haliangiales order have been previously identified as opportunistic epibiotic predators [69–72].

In Hopland, Myxococotta are moderately active in 0-10 and 20-30 cm samples, but not as active as in Angelo (**Fig. 2**). One of the most active organisms in 20-30 cm Hopland soil is a *Pseudobdellovibrio exovorus*, an obligate predatory member of the Bdellovibrionales order [69, 73]. Higher activity of these putative predators was detected during the wet-up and not during the dry-down.

### Structure-guided annotation reveals mechanisms of predation

To understand the way active predators within our grasslands influence the microbial community, we annotated hypothetical and uncharacterized proteins through the use of structural models. Since structure is more conserved than sequence [74, 75], there is a good likelihood that a shared conserved structural motif can uncover potential functions within the sequences left uncharacterized after sequence similarity searches. Within the MAG for Haliangiales we identified ten syntenic genes, coding all of the most crucial parts of a likely contractile injection system (CIS). After modeling each protein separately and using an available structure (7B5H) as reference [76], we assembled a coherent macromolecular model (**Fig. 7A**), many parts of which are interlocking and have minimal conflicts that are limited to N- or C-terminal regions. Our CIS model has all features necessary for function with very good RMSD values compared to a reference CIS system, including the tube (2.5 RMSD; **Supp. Fig. 30**) and its sheath (4 RMSD; **Supp. Fig. 31**), end tube cap (3.7 RMSD **Supp. Fig. 32**), a spike (3.5 RMSD, **Supp. Fig. 33**) with a PAAR-like domain for its tip (reference lacks PAAR, **Supp. Fig. 33**), and a baseplate cage (**Fig. 7B** 5.3 RMSD comparable to phage T6 baseplate [77]). The baseplate is formed from a heteropolymer of three separate proteins, which fold reasonably well together, based on AlphaFold predicted alignment error (PAE) values (**Supp. Fig. 34**). Assuming three layers of the tube compartment, this complex includes 73 proteins that together form a macromolecular machinery of size 2.8 x 2.6 x 3.8 nm with approximate weight of 3.15 megadaltons. CIS include recycling factors for when they need to be disassembled and such a factor (AAA+ recycling ATPase) is also present within the same genomic region as the CIS genes (**Fig. 7 C**). We searched for similar genomes to the Haliangiales MAG and found that several closely related organisms also contain such a syntenic CIS region with high similarity to our sequences (**Fig. 7 C**). No similar structure was found in the Pseudobdellovibrio MAG, using curated HMMs for CIS [78] and T6SS.

**Figure 7:**
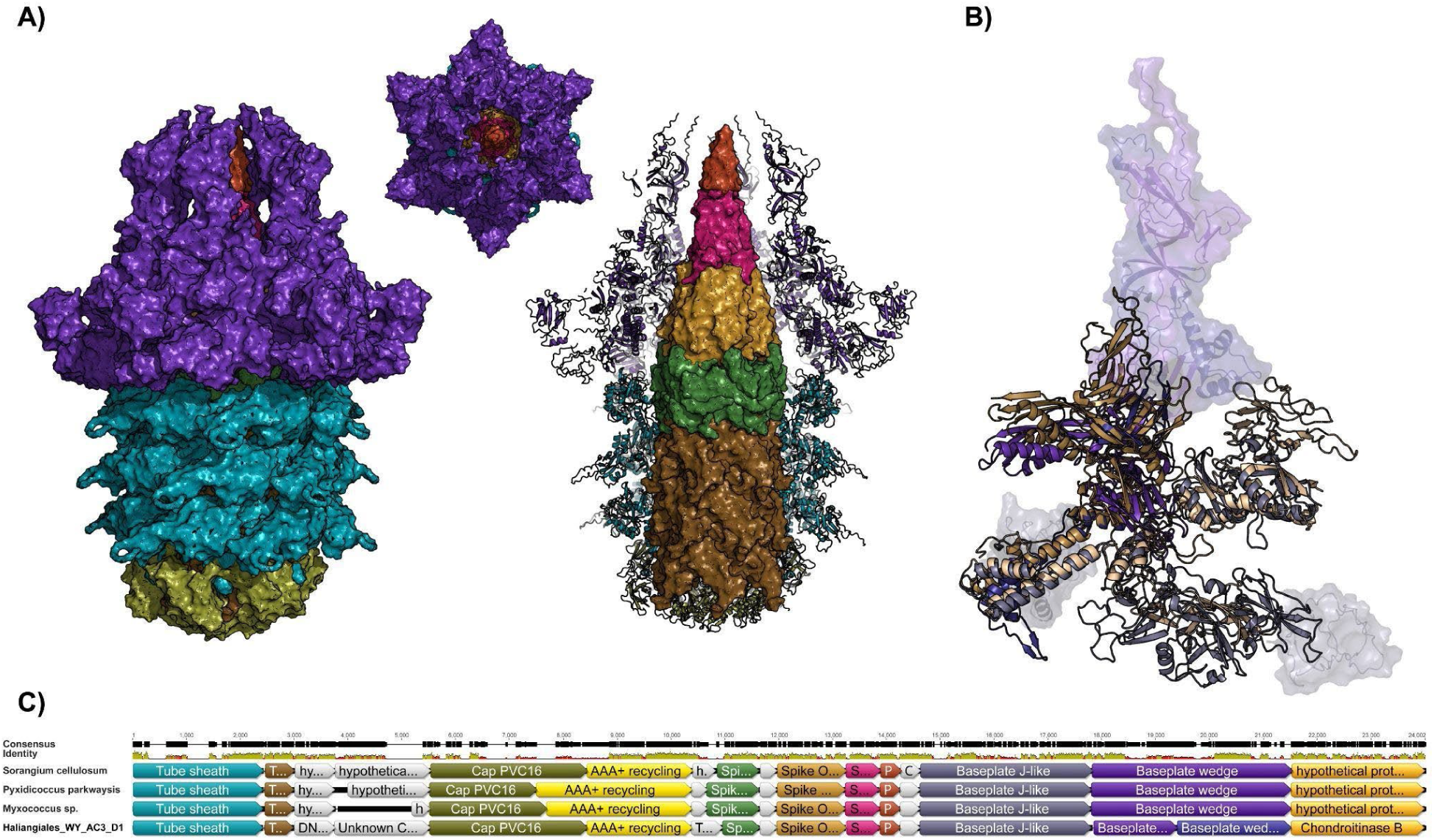
Structural model and synteny of the Haliangiales contractile injection system. A) Modeled structure of the Haliangiales MAG contractile injection system. Shown are side surface view, top view, and inner tube and spike view. Colors indicate different monomers of the complex. B) Superimposition of baseplate heterotrimer to the T6 phage baseplate protein gp6 (PDB: 5IV5). Parts not shared with the phage protein are shown with transparent surface and cartoon. C) Synteny of the CIS genomic region for a few Myxococotta species and our MAG sequence. Colors indicate different monomers of the complex and match panels A) and B).

The baseplate cage in the Haliangiales MAG was modeled using the phage T6 gp6 baseplate protein because there is too much structural divergence to other CIS structures. It is logical to have greatest variability in the structure of the outermost layers of this complex, since it will have roles in host identification and contact. It is possible that host specialization generates unique conformations of the baseplate which are hard to predict.

Myxococcota rarely contain CIS genes [78, 79] and no representatives of the Haliangales order have been observed to carry them. Members of the phylum are known to export their effectors through protein secretion systems of the types II, III, and VI [80–82], and to use a Tad-like apparatus for contact dependent killing [83]. Likely the CIS has been overlooked in Haliangiales because HMMs crafted to match the VgrG-like spike and baseplate proteins do not detect the truncated nature of the proteins, while structural annotation has no issue finding the structural homologies.

### Distribution of T6SS and CIS within MAGs from the Water Year

To elucidate the activity of injection systems throughout the Water Year we used sequence similarity searches across all metagenomic bins. We used curated HMMs for all elements of the CIS [78] and the T6SS (**Supp. Table 1**). For this broad survey we considered only MAGs with at least 4 gene hits in either CIS or T6SS protein groups. No clear relationship between activity and any specific CIS or T6SS gene is apparent (**Supp. Fig. 35**). To connect injection systems presence with the spatiotemporal variables, we filtered out MAGs that do not have hits against all three of the tube, sheath, and spike proteins, assuming that will be a good approximation of an actual CIS or T6SS presence. Sequences from Acidobacteriota, Alphaproteobacteria, and Betaproteobacteria show T6SS activity in Angelo, with Acidobacteriota active in 0-10 cm, Betaproteobacteria in 20-30 cm and together with Alphaproteobacteria in 50-80 cm depths (**Supp. Fig. 36**). Acidobacteriota species containing T6SS show activity also on Hopland 0-10 cm and 20-30 cm. Additionally, Verrucomicrobia, Planctomycetota, Blastocatellia, and Chloroflexota show activity in 0-10 cm, with a clear increase in activity at the time points immediately before and after the first fall wet up (**Supp. Fig. 36**).

Using the curated CIS HMMs, we were unable to detect the spike protein from our Haliangiales MAG, likely because the truncated nature in the spike protein produces too short hits in the sequence search. Exploring the rest of the active species, containing CIS we see that there are few hits in Angelo shallow soils. The only active representatives are from the Actinomycetes class at 50-80 cm depth (**Supp. Fig. 37**). In Hopland shallow soils, Bacteroidota, containing the CIS show high levels of activity during the peak plant productivity and after the fall wet up (**Supp. Fig. 37**). Other active CIS-containing groups in deeper Hopland soils are Actinomycetes and Deltaproteobacteria. The presence of these predatory machinery in deep soils uncovers a surprisingly complex and rich microbial environment that could support higher trophic levels.

### Haliangiales MAG contain likely secretion effectors

To determine the effect of the Haliangiales CIS, we used sequence homology to identify proteins that can influence host cells. Possible effectors present in the MAG are 14 genes containing von Willebrand domains (Supp. Data 5), found to be present in cytotoxic proteins of *Legionella* [84] and *Liberibacter asiaticus* [85]. Our *Haliangiales* metagenome also contains rearrangement hotspot repeat proteins (RHS) (Supp. Data 5), known to act as encapsulation devices for toxins [86].

To expand the set of possible effectors we applied structural homology searches within hypothetical proteins. A likely effector is located immediately downstream of the genes for the injection system - protein, containing the parallel beta-helix domain of a possible alginate lyase, and also found in phage tail spikes [87, 88]. This protein and its location are conserved across other Myxococcota (**Fig. 7C**). We also identified a gene with structural similarity to a TAL effector repeat protein (**Supp. Fig. 38**) found in plant pathogens from the genus *Xanthomonas* [89, 90].

## Discussion

The grassland soils studied here are populated by many and diverse groups of soil microorganisms, some of which form clades that may be essentially specific to grassland soils (**Fig. 6**) and whose activities vary with depth and season (**Fig. 2, 4**). These organisms likely respond to specific periods of water and organic carbon substrate availability. Spatial variables and pH are strong predictors of the community composition while seasonal timepoints are not. Although this has been reported previously in shallow soils [91, 92], it is notable this pattern holds in deep soils, which have rarely been investigated. Our study is the first to provide ^18^O (thus C source-independent) SIP microbial activities at species level for deep grassland soils (>50 cm).

Deep soils store more than 50% of the total soil C stocks [93, 94] and have higher CO_2_ concentrations compared to shallow soils [95–97]. In fact, in a broad study of soil profiles, it was shown that the 30 - 100 cm depth region stores as much carbon as is present in the 0 - 30 cm region [94]. There are very few SIP studies in deep soils (>20 cm) [98, 99], and to our knowledge, no SIP studies have used whole genome sequencing in soils deeper than 50 cm. We identified a plethora of organisms exhibiting high levels of activity in deep (50-80 cm) soils (**Fig. 2,4**). These results are congruent with observations of deep soils having faster carbon mineralization rate, when normalized for microbial C biomass, compared to shallow soils [100] [101]. We show that archaeal species are not only present in deep soils [38, 39], but also have moderate AFE values reaching up to 0.1 (**Fig. 2,4**). Active archaea include Nitrososphaerota, Thermoproteota, and Euryarchaeota, which have roles in autotrophy, oxidation of ammonia and carbon turnover. Thus, we conclude that Archaea are important in biogeochemical cycling within deep soil. The metagenomics-informed SIP can detect active organisms that are often missed by amplicon sequencing, especially some CPR bacteria [52] and Archaea [102].

Eukaryotic viruses are overlooked in amplicon sequencing studies and their eukaryotic hosts are often missed due to primer choices. Studies have shown that viruses infecting fungi and oomycetes are abundant in soils, influencing fungal community dynamics [103–105]. Given soil complexity, it is notable that we reconstructed large fragments of a >381 kbp of the genome of a giant eukaryotic virus from deep grassland soil. Isotope labeling of the viral genome suggests that it was replicating in the genome of an oomycete that was seemingly active upon laboratory wet up for isotope labeling (**Supp. Fig. 21**). In situ, viral replication could impact turnover of carbon (probably root-derived) in deep soil by the saprophytic hosts.

### Seasonal dynamics of active cohorts in shallow soils

The active subsets of soil microbial communities in shallow soils experience significant seasonal shifts, likely tied to water availability (**Fig. 3**). During wet-up, either after the fall rain (Hopland) or in the laboratory wet up experiment of the July sample (Angelo), we observed large groups of organisms that were only active in these samples (**Fig. 2-4**). This pattern suggests the presence of specialist microbes, able to take advantage of the sudden moisture and nutrient availability. This phenomenon is consistent with the concept of "hot moments", where microbes spike in activity under favorable conditions [106, 107]. Very few organisms are active only in May (**Fig. 3, Supp. Fig. 16**), but these organisms show relatively low activity compared to organisms that are active in May and other time periods. This indicates that the most active organisms during peak plant productivity (May) are generalists. The abundance of diverse nutrients derived from growing plants may support a broad range of organisms [108], limiting niches for specialists.

### Highly represented functions in Angelo and Hopland cohorts

The most highly represented active microbial functions relate to degradation and processing of diverse and complex carbon compounds and their breakdown products that feature in central carbon metabolism. The data suggest continuous availability of organic matter year-round and underscore the fundamental role of soil microbes in decomposing plant and fungal biomass throughout the seasons [109].

The distinct functional profiles in the May cohorts, particularly the increased presence of nitrate and nitrite metabolism genes, indicate a larger role of nitrogen cycling processes at this time point relative to other time points. This could be linked to changes in soil moisture and temperature, enhancing nitrification and denitrification activities that reflect microbial response to elevated nitrogen availability. The presence of arsenate reductases might suggest microbial strategies to mitigate arsenic toxicity under fluctuating redox conditions typical of spring soils [110].

In June, the heightened capacity for thiosulfate oxidation among active organisms suggests a significant role of sulfur cycling during early summer [111]. The findings suggest microbial release of reduced sulfur from biomolecules such as polysaccharides, compatible solutes, amino acid degradation.

### CPR bacteria have complex relationships with the microbial community

The most active organisms in shallow Angelo soils are representatives of the CPR from the Saccharibacteria phylum [52, 112]. Previously, evidence was reported for the incorporation of ^13^C derived from plant exudates into the DNA of rhizosphere Saccharibacteria [54]. Notably, in that study the identified Saccharibacteria genomes carry genes only for conversion and modification of nucleotides, but do not contain complete nucleotide synthesis pathways. As these bacteria do not have the ability to synthesize nucleotides, it was inferred that isotope incorporation occurred though recycling of labeled nucleotides derived from other organisms or viruses [54]. It should be noted that other members of the CPR, such as Peregrinibacteria, have been observed to contain nucleotide synthesis pathways [113], however to our knowledge this is the first time Saccharibacteria have been identified to carry complete nucleotide synthesis pathways (**Fig.4, Supp. Fig. 18**). As our Saccharibacteria are predicted to be able to synthesize DNA *de novo*, isotope incorporation into newly produced DNA of these episymbionts does not imply recycling of DNA synthesized by other organisms, but incorporation during independent growth. The presence of ubiquinol oxidases and F-type ATPase opens the possibility for aerobic respiration, with greater energy generation than would be afforded by fermentation (their typically predicted metabolism). This may contribute to the high AFE values reported for Saccharibacteria in the current study.

Predatory behavior of Saccharibacteria associated with some Actinomycetota hosts was reported in some co-cultivation experiments conducted using organisms from the oral microbiome under resource limiting conditions [51]. This is in agreement with the presence of systems for concentrating CO_2_ as bicarbonate in Saccharibacteria active during the dry-down timepoints, since bicarbonate is a known controller of toxin expression [58, 59]. It is possible the Saccharibacteria is using those genes to control its epibiotic cycle, based on carbon availability.

### Actinomycetota have broad activities in time and depth

Our findings build upon previous research at the Angelo site [38] and contribute new insights into the phylogenetic and functional diversity of soil Actinomycetota by identifying two major new clades. The expansion of the Rubrobacteridae class is particularly notable, as signature protein analysis identifies it as the deepest branch within Actinomycetota [114]. The novel clades’ enrichment in thiosulfate oxidation capabilities (**Supp. Fig. 26-29**) points to a role in sulfur cycling. We found Actinomycetota change in activity levels over time points and vary in activity levels between the two grassland soil ecosystems, and with soil depths. Notably, the cohort active only in May and only at Angelo is composed exclusively of Actinobacteria, including a single representative from novel clade 1.

### Bacteroidota active in Hopland

Bacteroidota are the most active organisms detected in the current study, active only in Hopland soils, at all depths, and almost all time points. These bacteria are known to participate in remineralization of organic materials into micronutrients and our CAZy analysis supports this (**Supp. Data 3**). The lack of active representatives from Bacteroidota in the two driest timepoints (0-10 cm at June and September before rain), suggests that they require prolonged periods of >5% moisture to be active. Their activity throughout all timepoints in deep soil (**Fig. 2**) likely reflects persistent moderate levels of moisture throughout the entire year (**Supp. Fig. 1**).

### Predatory bacteria

The presence of highly active predators in the wet-up period but not during the dry-down (**Fig. 2,4**) supports the ecological theory that predatory control of lower trophic levels increases with higher ecosystem productivity [115–117]. Our results of highly active predatory bacteria are in agreement with previous surveys that show higher growth and carbon uptake in predatory bacteria compared to non predators [118].

### Haliangiales effectors

We identified a likely predator from the Haliangiales order as highly active in shallow Angelo soils (**Fig. 4**). The draft genome for that active Haliangiales contains genes for a contractile injection system which is evolutionarily related to phage tails and type VI secretion systems, known to support predatory lifestyle (**Fig. 7**). We identified a likely alginate lyase, syntenic with the contractile injection system genes (**Fig. 7**). Lyases of this form are capable of cleaving alginate, a component of bacterial cell walls [119] and biofilms, and this can be an effective way of lysing cells during predation. Other possible effectors we identified in the Haliangiales genome are rearrangement hotspot proteins, known to facilitate intercellular competition [120] and found to be essential in the function of type VI secretion systems for some bacteria [121]. Finally, the presence of von Willebrand domains and transcription activator-like effectors (**Supp. Fig. 38, Supp. Data 5**) could be an indication that Haliangiales can target eukaryotic cells, given those are effectors that work best against higher eukaryotes and plants.

## Conclusions

This study provides new insights into how the active fractions of two Mediterranean grassland soil microbiomes, shaped by contrasting rainfall inputs, shift over the course of a year in response to seasonal water availability. While overall microbial community composition remains relatively stable, we uncover significant temporal and spatial variability in the active organisms, with distinct patterns emerging across soil depths and locations. Deep soils (>20 cm), often overlooked in microbial studies, exhibit clear evidence of in situ microbial activity linked to specific organism groups.

We highlight the role of diverse ecological strategies, including predatory lifestyles, exemplified by active Haliangiales organisms in shallow Angelo soils, which we found to encode a contractile injection system. Other strategies observed to have high levels of activity across all timepoints involve episymbiotic Saccharibacteria who unexpectedly revealed nucleotide synthesizing capabilities, challenging assumptions about their relationship with the rest of the soil microbiome. This study underscores the importance of integrating activity data with depth and seasonal variability to better understand microbial dynamics in grassland soils.

## Acknowledgements

We thank Mengting (Maggie) Yuan for valuable discussions and help in the greenhouse, Jeff Kimbrel for support in analyzing qSIP data, Christina Ramon for help with total C/N processing, and Leila Wahab for sharing soil bulk density information. Furthermore, we would like to thank Alison Smith, Troy McWilliams, Greg Solberg, and the rest of the Hopland Research and Extension Center and Angelo Coast Range Reserve staff for logistical and sampling support. We thank all members of the Banfield and Firestone labs, and LLNL staff scientists who helped with soil sampling and soil processing. This research has been supported by the US Department of Energy, Office of Biological and Environmental Research, Genomic Science program LLNL “Microbes Persist” Scientific Focus Area (award no. SCW1632).

## Author contributions

The study was designed by J.F.B, S.J.B., J.PR., and P.I.P. Sampling design and organization was performed by K.EM. and P.I.P. Stable isotope incubations and DNA extractions were performed by K.EM. and P.I.P. Density fractionation of DNA was performed by G.M.A. Raw read quality control was performed by R.S. Assembly, binning, and genome analysis were performed by P.I.P. with the help of R.S. and S.L. R.S. and S.L. contributed to data handling on ggKbase. All bioinformatic data analysis and figure generation was performed by P.I.P. with input from J.F.B, S.J.B., and J.PR. P.I.P. and J.F.B. wrote the paper with input from all the authors.

## Data availability

Metagenomic data generated in this study have been submitted to the NCBI BioProject database (https://www.ncbi.nlm.nih.gov/bioproject/) under accession number XXXXX. The Haliangiales and NCLDV genomes can be accessed via ggKbase (https://ggkbase.berkeley.edu/xxxx).

## Methods

### Study sites, sampling, and edaphic measurements

Soil samples were collected from the Buck field in the Hopland Research and Extension Center and south meadow field site in the Angelo Coast Range Reserve. Triplicate soil cores were extracted with a Geoprobe 54LT when the weather permitted, and manual slide-hammers were used when the soil was too wet to support the Geoprobe. When using manual slide-hammers, different sterilized 3 by 6 inch cylindrical polycarbonate inserts were used to reach a depth of at least 80 cm. Each replicate was taken from a 30 m^2^ circular plot. In total 9 samplings were performed at each site. Inserts were sealed, transported to UC Berkeley, and processed on the same day. Five depth profiles were established: 0-10 cm, 10-20 cm, 20-30 cm, 30-50 cm, and 50-80 cm. Soil was separated for edaphic measurements, for stable isotope incubations, for viromic extractions (0-20cm), and for frozen and dry archive. Soils for the frozen archive were flash frozen in liquid N_2_ and stored at -80°C. Sampling started in April 2022 and ended in November 2022, following the major water dynamics of the year. Starting with the spring peak plant productivity in April-May, followed by the summer dry-down and finally sampling within 72 hours of the first wet-up in the fall.

For the purposes of performing isotope incubation, initial Gravimetric Water Content (GWC) was measured after an overnight oven-drying at 100°C. Final GWC was estimated by drying soils at 100°C until the weight of samples stabilized. Nitrogen and carbon concentration (%N dw and % C dw, respectively) were estimated by manually removing visible root material from oven-dried soils, grinding the soil with mortar and pistil and sieving out any rocks, through light tapping of the mortar. Ground soil was then wrapped in tin capsules and analyzed using a CHNOS Elemental Analyzer (vario ISOTOPE cube, Elementar, Hanau, Germany) at the Center for Stable Isotope Biogeochemistry at the University of California, Berkeley, CA, USA. The percent calibration and quality control assessment were based on NIST (National Institute of Standards and Technology, Gaithersburg, MD, USA) reference materials were certified % C and % N values.

We measured the acidity of each soil sample using 10 ml of 0.01M CaCl_2_ and 5 grams of soil. Measurement was done with a calibrated Oakton 110 series pH meter (Eutech Instruments, Malaysia). Each sample was measured twice within a 5 minute time period and the average of the two values was reported. If the two measurements were different by more than 0.5 units a third measurement was performed, removing the outlying value.

Moisture retention curves were generated using the UC Davis Analytical lab where soil was brought to near saturation and allowed to equilibrate under a set atmospheric pressure potential [122].

### Stable Isotope Incubations and CO2 measurement

Large portions of the soil microbial community can be inactive. Thus, making conclusions about microbial activity, only on the basis of read mapping inferred relative abundance can produce incorrect conclusions. To discern between active and inactive portions of the community we applied H_2_^18^O stable isotope probing. To ensure enough isotope incorporation and to minimize baseline ^16^O-water content, soils were gently dried down to around 10% GWC in a room temperature hood, when needed as determined by initial GWC values. Five grams of soil were measured in cryo-pods (Fisher Scientific 5005-0015) for shallow samples (0-20cm) and ten grams of soil were measured for deeper soils (20-80cm). After drying, t0 samples were frozen and kept at - 80°C until DNA extraction. Half of the samples received enough natural abundance H_2_^16^O to bring the sample up to 25% GWC, and the other half received 97-atom % or 98.5-atom % H_2_^18^O (Millipore Sigma 902187-50G) to bring the sample up to 25% GWC. To measure CO_2_ production during incubation, samples were placed in airtight glass jars fitted with septum for seven days. Five ml of headspace were sampled at zero, four, and seven days using a syringe in evacuated vials. The CO₂ concentration was measured by taking five ml with a syringe from the sample vial and using a Li-Cor 850 CO₂ analyzer. To produce CO_2_ concentrations, raw gas data was analyzed with the following script [123]. After the incubations were completed, soils were harvested and stored at -80°C until DNA extraction.

### DNA extraction

Due to time constraints and constraints on the amount of sequencing we can perform, we extracted DNA from five timepoints and three depths: 0-10 cm, 20-30 cm, 50-80 cm. In total 276 samples were randomized so that each batch of processing does not contain more than one sample from the same ecosystem, time, and depth. DNA extractions were performed using the QIAGEN DNeasy PowerMax Soil Kit (12988-10) using a modified version of the manufacturer’s protocol in batches of 12 samples. The modifications were used on samples from deep soil profiles (>50cm) and included additional steps of back-extractions [124], allowing us to obtain enough DNA for the subsequent fractionation. DNA quality was established with Nanodrop and quantified with Qubit. Four ug of DNA were aliquoted for fractionation from the incubated samples.

### DNA fractionation

High-throughput stable isotope probing (HT-SIP) was completed on DNA following a modified version of the previously published protocol [125]. Briefly, 4 µg of DNA was added to 1xTE buffer to 150 µl, 0.95 mL gradient buffer (10 mM Tris-HCl pH 7.5, 1 mM EDTA pH 8.0 and 100 mM KCl), 5 µl gradient buffer plus 0.1% Tween-20 and 4.5 ml of cesium chloride (CsCl) stock 1.885 g ml-1. Samples brought to the final density between 1.725-1.730 g ml-1. This mixture was added to a 5.1 ml ultracentrifuge tube (Beckman Coulter Cat. 342412). Density gradient was established by centrifuging samples for 108-136 h at 176,284 x g at 20 °C in a Beckman Coulter Optima XE-90 ultracentrifuge using a VTi65.2 rotor. CsCl gradient was fractionated into 22 fractions (∼250 µl each) using an Agilent Technologies 1260 Isocratic Pump and 1260 Fraction Collector. CsCl was displaced with sterile water pumped at 0.25 ml min-1. Density of each fraction was determined using a Reichart AR200 digital refractometer with modified lens cover to read 5 µl. Using a Hamilton Microlab STAR liquid handling robot, 500 µl PEG solution (30% PEG 6000, 1.6 M NaCl) and 35 µl of 1:5 diluted Glycoblue (Invitrogen, ThermoFisher Cat. AM9515) was added to each fraction and incubated overnight at room temperature. Samples were spun at 4198 RCF for 5 h at 10°C. PEG was removed and 950 µl 70% ethanol was added to each fraction. Samples were spun at 4198 RCF for 1.5 h at 10 °C. Ethanol was removed and DNA was resuspended in 40 µl of 1x Tris-EDTA (pH 7.5). DNA concentration was quantified with a PicoGreen fluorescence assay (Invitrogen, ThermoFisher Cat. P7581).

Samples were processed in batches of 16 including ^16^O and ^18^O samples. Due to technical reasons two batches had a shift in the densities that affected all samples within a batch (**Supp. Fig. 39**). This resulted in some samples showing ^18^O densities which are lower than the same for a corresponding ^16^O sample. Such batches were adjusted by taking advantage of the fact that ^16^O samples should have the same mean density distribution. A linear regression was calculated from the densities of all ^16^O samples that showed expected densities and reasonable shifts between paired ^16^O and ^18^O samples (**Supp. Fig. 40**). Likewise, a linear regression was used to represent each of the samples from problematic batches, then the difference between the two lines was calculated and the densities of the problematic samples were adjusted by that difference (**Supp. Fig. 41**).

### Fraction binning and sequencing

To lower the number of sequencing runs, fractions were binned based on DNA amounts and densities down to five bins, totaling 890 fractions. The binning cutoff values were calculated based on ^18^O samples and involved iterative calculation which ensured that no bins have overlapping densities, and each binned sample will have enough total DNA for sequencing (**Supp. Fig. 42**). The density thresholds used are 1.72904, 1.72240, 1.71584, and 1.70770. The code used for determining binning thresholds and performing linear regression can be found at https://github.com/petaripenev/SIP/blob/master/scripts/parseSIPoutput.py.

Metagenomic library preparation and DNA sequencing were performed by Novogene. Metagenomic libraries were prepared for sequencing on an Illumina Novaseq 6000 platform, producing 150 bp paired-end reads with a target inter-read spacing of 350 bp.

### Metagenomic assembly and annotation

Raw reads were initially assessed with FastQC (http://www.bioinformatics.babraham.ac.uk/projects/fastqc/). Illumina adapters were trimmed with bbduk v37.50 using parameters: k=23, mink=11, hdist=1, tbo, tpe, ktrim=r ftm=5 and screened for phiX and Illumina artifacts with k=31, hdist=1. Next, reads were quality trimmed using sickle v1.33 and default parameters. All 89 samples were individually de novo assembled using megahit v 1.2.9 [126] with the following parameters: -k-min 29, -k-max 255, –low-local-ratio 0.333. Sequencing coverage of each contig was calculated by mapping raw reads back to assemblies using bbmap v39.01 (https://sourceforge.net/projects/bbmap/) with parameters: -minid 0.97, - ambiguous random. Annotation was performed on contigs larger than 1000 nucleotides and included prediction of 16S and tRNA with cmsearch v 1.1.4 [127] and tRNAscan-SE v1.3.1 [128]. Open reading frames (ORFs) were predicted with prodigal v 2.6.3. Genes were annotated against kegg [129], uniprot [130], and uniref90 with diamond v 2.1.8 [131], using parameters: -e 0.00001, –sensitive, –iterate, –max-target-seqs 1, -c 1, -b 20.

### Abundance analysis

To estimate the microbial community composition we applied rpS3 abundance analysis, which has previously proven to give a good species-level representation of metagenomic samples [38]. In brief, rpS3 genes were identified from all assembled contigs larger than 1000 nucleotides with a custom HMM, using hmmsearch v3.3 [132]. The nucleotide sequences of those genes were clustered with vsearch v2.13.3 at 99% identity. For each cluster the longest contig was selected for read mapping, which was done with bbmap with parameters -minid 95 and -ambiguous random. Counts were calculated with coverm using the tpm method and taxonomy was established by blast against the a rpS3 subset of the NR database.

### Atom Fraction Excess calculation

To estimate activity of species, we calculated atom fraction excess for each SG defined from the rpS3 abundance analysis (“features” in qSIP pipeline). Reads from the fractionated samples were mapped to the contigs representing each rpS3 cluster with bbmap using perfectmode. Coverage of each contig was calculated with coverm v 0.7.0 in contig mode with the reads_per_base method. The resulting count table was used with the qSIP2 R package (https://jeffkimbrel.github.io/qSIP2/index.html) to calculate AFE values following the standard qSIP workflow. Features were filtered to be present in minimum three sources (biological samples) and four fractions, except for 50-80 cm Hopland samples from the May sampling where we used a minimum of two sources. This was done because replicate A failed to produce enough DNA for fractionation and sequencing (**Supp. Data 2**) and was removed from the experiment. Confidence intervals for each feature (**Fig. 2,4**) were calculated with 1000 resamples at 0.9 confidence interval.

### Genome binning, dereplication, and metabolic pathway analysis

To explore the functional makeup of our dataset we performed genomic binning, using multiple binning softwares. To generate sufficient mapping data for the depth-based binners we performed all vs. all mapping using bbmap in perfectmode between the samples and the reads. However, due to the size of the project we limited this to samples from the same depth and the same ecosystem. For binning we used metabat2 v2.15 [133], concoct v1.1.0 [134], maxbin2 v2.2.7 [135], and vamb v3.0.2 [136]. Results of all four binning softwares were combined with DASTool [137], their quality was determined with checkm2 [138], and were de-replicated with dRep [49]. In total we generated 4493 de-replicated bins. By connecting the presence of rpS3 clusters to bins we identified that 6560 SGs have detected activity and are binned, while 1044 are active and are unbinned. Metabolic pathway analysis was performed with DRAM [139] on all de-replicated metagenomes.

### Seasonal cohort definition

To understand the makeup of microbial organisms during different timepoints we grouped them by activity, using distance covariance clustering. Clusters are determined by an all vs all pairwise distance correlation from AFE values across the five timepoints. Clustering was done after calculating significant correlations using a T-statistic and controlling for multiple tests with the Benjamini/Hochberg method (**Supp. Fig. 14,15**). Final clusters were determined using the ward method, followed by a threshold determination that used 15 clusters as the maximum number. Cohorts can include single or multiple clusters, merging clusters in a cohort was determined manually.

### Phylogenetic analysis

To explore the phylogenetic make up of Euryarchaeota, Saccharibacteria, and Actinomycetota within our project we build phylogenetic trees based on rpS3. Reference protein sequences from each lineage were collected from NCBI, using the phylogeny browser. Those sequences were clustered with cd-hit [140] at 90% identity. Alignments were constructed based on a reference structure-guided alignment from the SEREB database [141, 142] by using mafft [143] with the –addfull parameter. Heavily gapped regions from the resulting alignments were removed with trimal [144]. Finally, phylogenetic trees were built with IQ-TREE v1.6.12 [145] with 1000 ultrafast bootstrapping [146] and the automatic model finder. The model finder identified LG+R5 as the best model for the Euryarchaeota tree, TIM2e+R5 for the Saccharibacteria tree (we used nucleotide sequences), and LG+R9 for Actinomycetota tree.

### Structural modeling

To identify proteins with functions related to predatory behavior we modeled hypothetical proteins with size up to 800 amino acids from the highly active Haliangiales MAG. We used the ColabFold [147] (v. 1.5.5) implementation of AlphaFold2 [148] with amber relaxation, three prediction recycles, a template search against the PDB database, and MSA construction against mmseqs2_uniref_env [149]. Predicted protein structures were queried with FoldSeek [150] against the PDB database to obtain functional annotations for each predicted structural domain.

The Haliangiales CIS was modeled using reference structures of CIS (7B5H, 7B5I) [76] and phage baseplate (5IV5, 5IV7) [77]. This was achieved by superimposing proteins from the CIS genomic region (**Fig. 6C**) to parts of the reference structures until a coherent and complete structure was achieved. To achieve that, proteins from the baseplate wedge and the cap were remodeled as complexes with ColabFold and AlphaFold3 [151].

