## Supplementary Information for "The active subset of grassland soil microbiomes changes with soil depth, water availability and prominently features predatory bacteria and episymbionts"

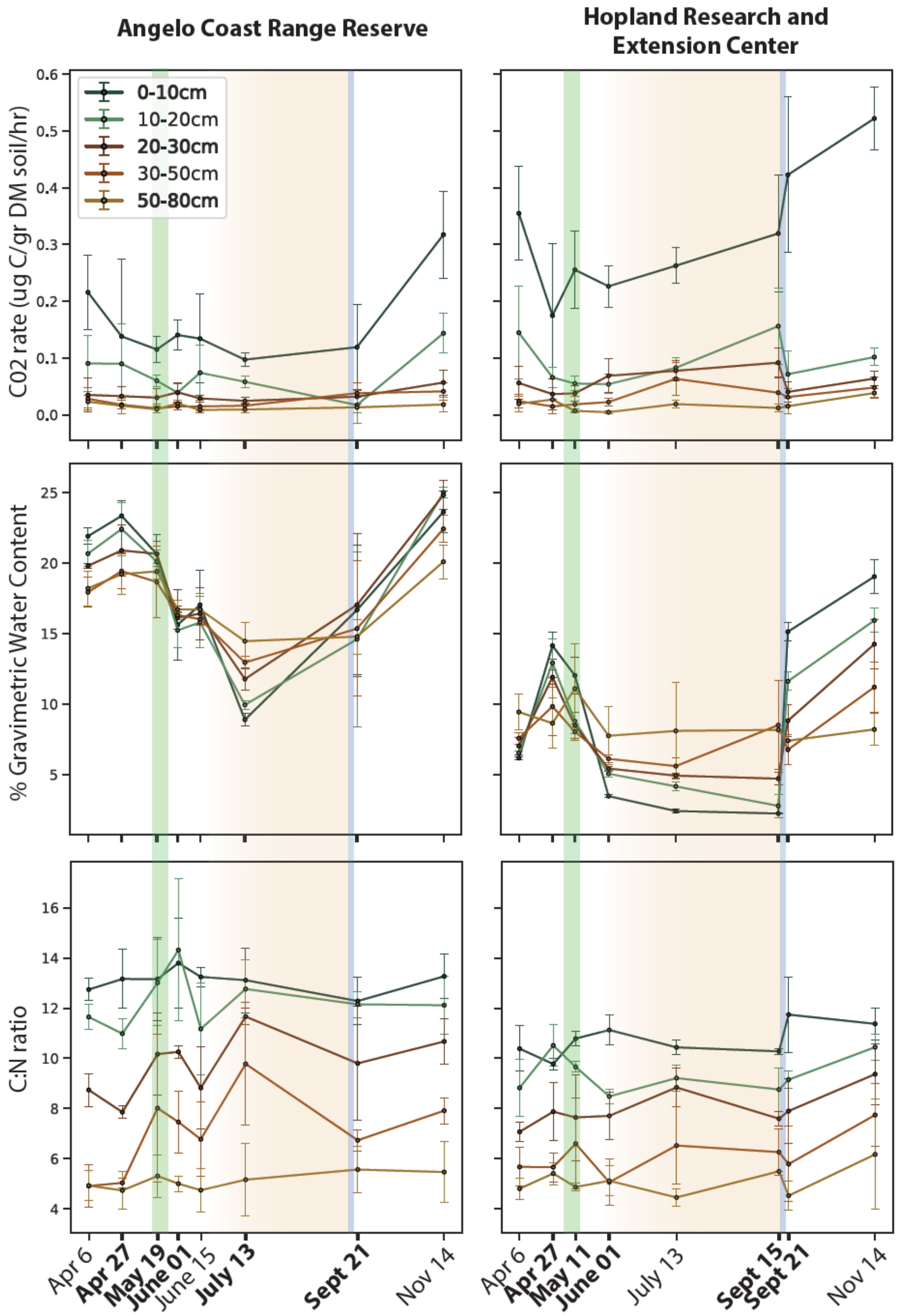


*Figure 1. Metadata measurements for Angelo (left column) and Hopland (right column). First row: CO2 rate measured during seven-day isotope incubations. Second row: gravimetric water content of the sample field moisture. Third row: ratio of total carbon against total nitrogen from the field samples. Error bars are calculated based on three biological core replicates. Important periods of the Water Year are shown with background colors: green - peak plant productivity; brown - drying down; blue - first fall rain event. Timepoints and depths of samples used for are indicated in bold.*

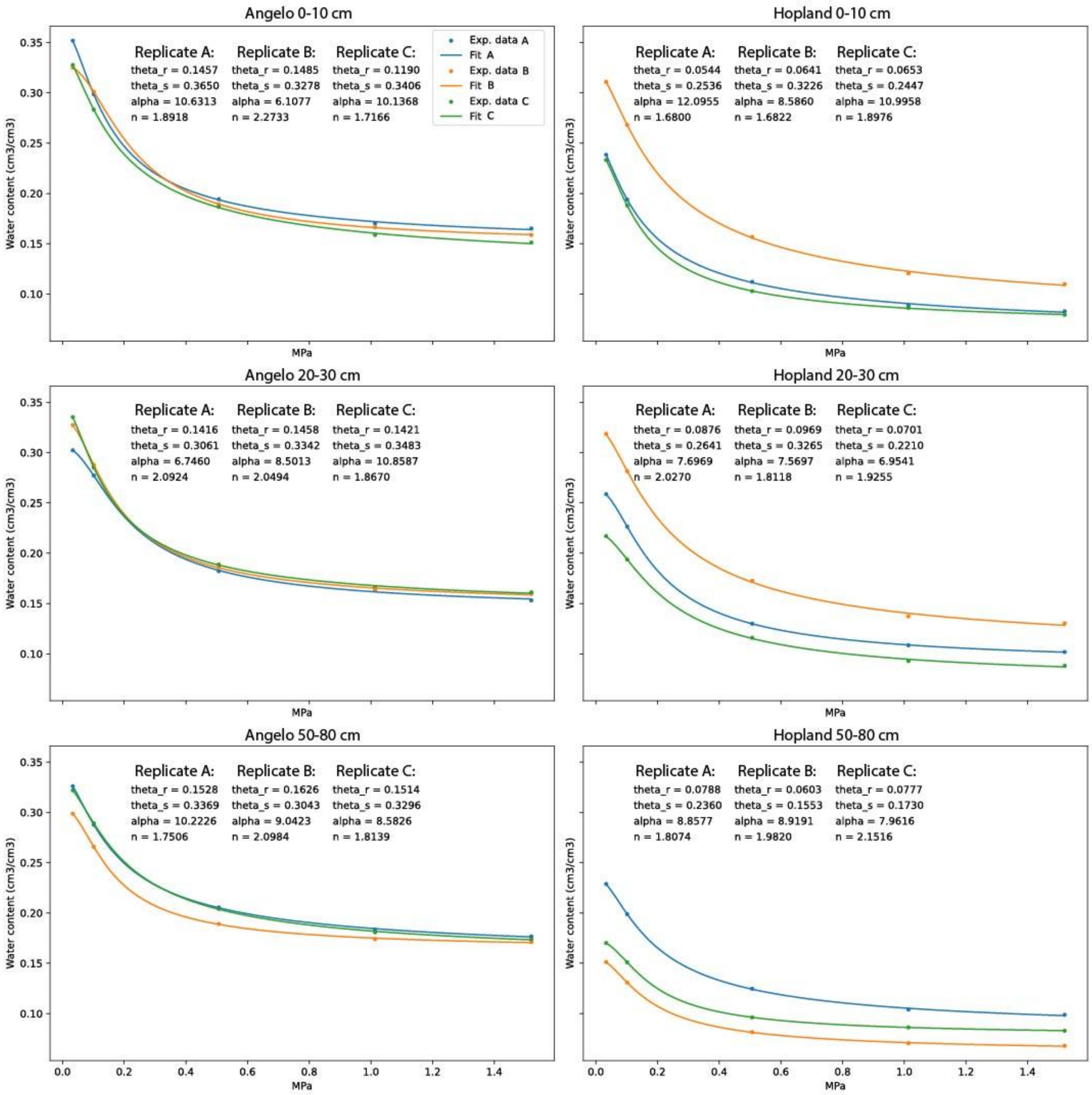


*Figure 2. Moisture retention curves for three replicates from each depth (rows) and ecosystem (columns). Each plot represents results of the calculated volumetric water content and head pressure for three replicates. Curves were fitted to the experimental data using the curve_fit function from the scipy package (v. 1.10.1) and the parameters were calculated using the van Genuchten equation. Bulk density used to calculate volumetric water content for Angelo was collected during a previous sampling trip (2020) and was cross checked with volumetric water content measurements from the weather station at Angelo South Meadow (https://dendra.science/orgs/sandbox/status/sandbox-south-meadow-ws). Bulk density for Hopland was sampled April 2023.*

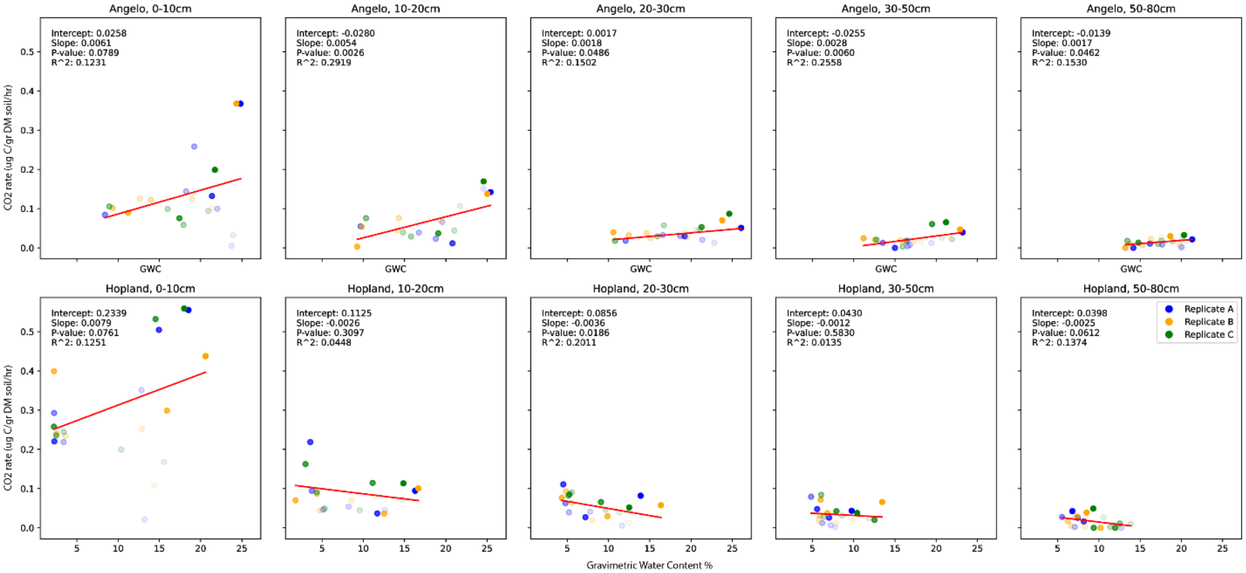


*Figure 3. Effect of gravimetric water content on CO2 rate for different ecosystems (rows) and depths (columns). Replicate A is colored blue, B is yellow, and C is green. Different timepoints are displayed with transparency (earlier timepoints are more transparent). The model line was fitted with a general linear model implementation of statsmodels python package (v. 0.14.1).*

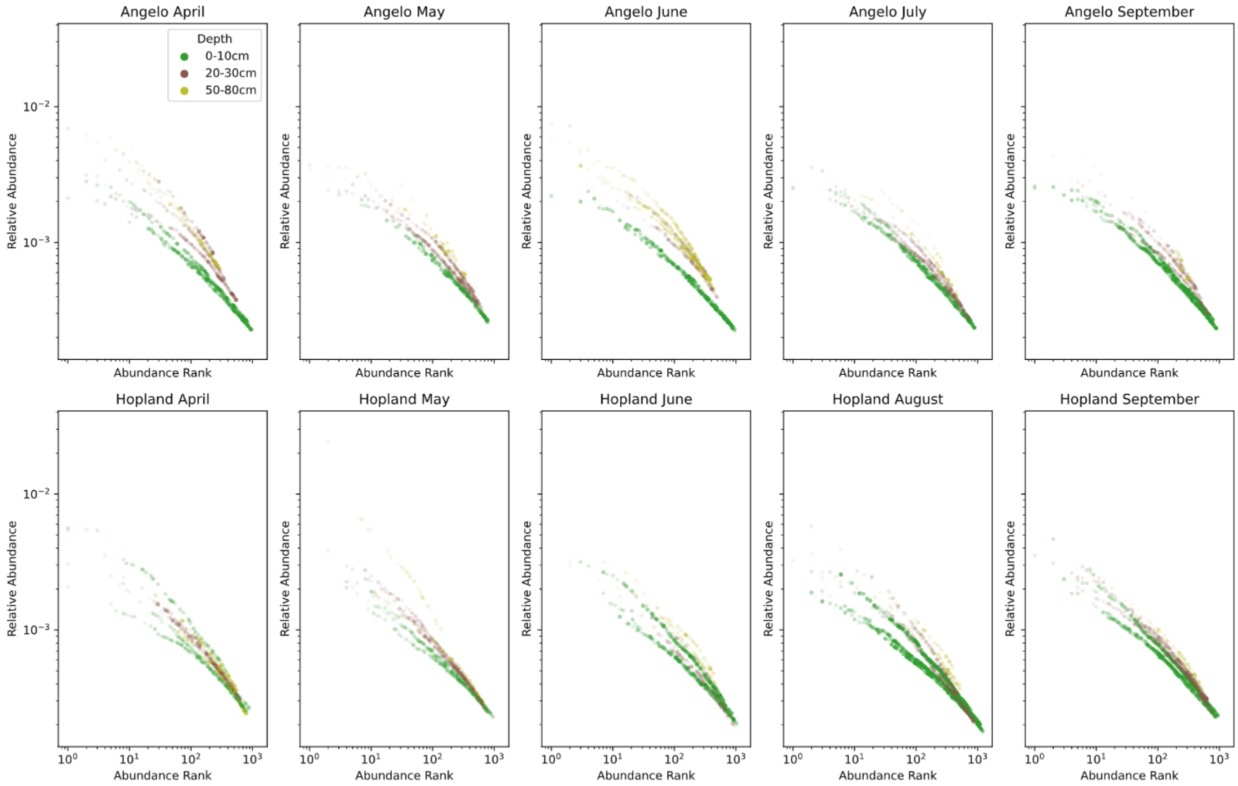


*Figure 4. Abundance ranks as a function of the relative abundances for active species accounting for the top 40% abundance for each ecosystem (rows) and timepoint (columns). Depths are colored with green for 0-10cm, brown for 20-30cm, and yellow for 50-80cm. Activity of each species as AFE values are indicated with transparency ranging from 0 – completely transparent to 1 – no transparency, normalized from 0 to the maximal value for a given combination of ecosystem and depth. Both axes are logarithmic.*

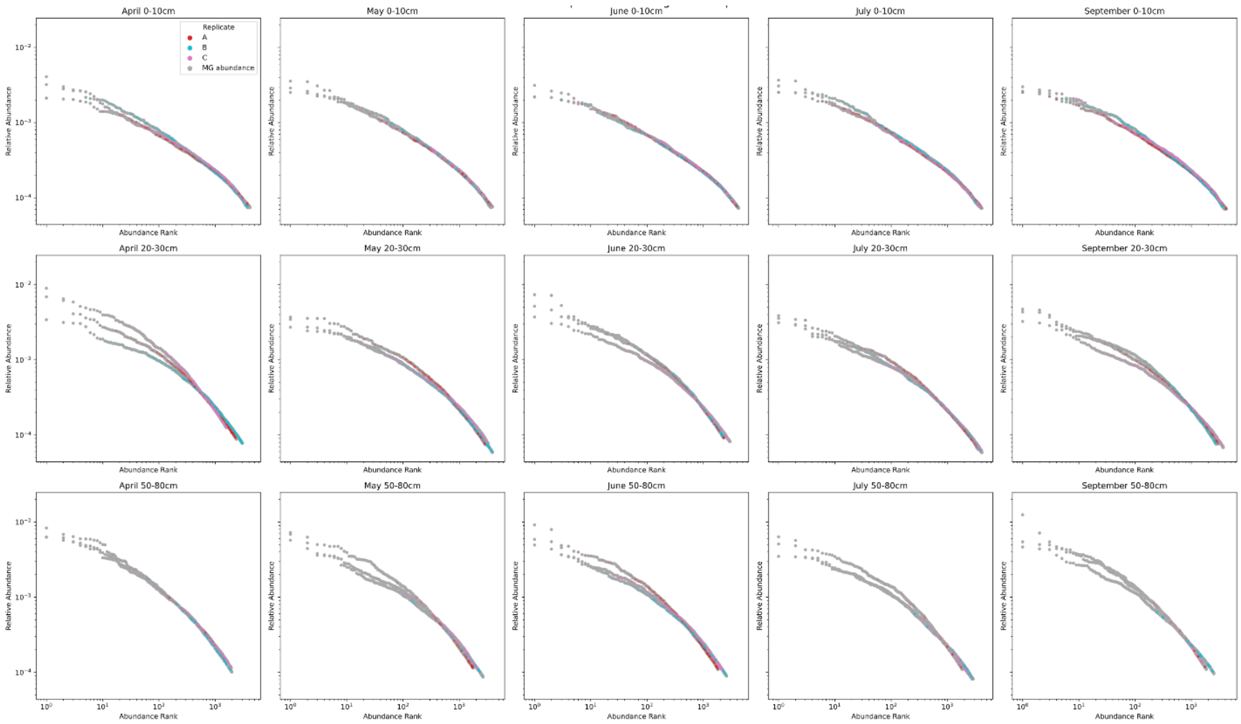


*Figure 5. Abundance ranks as a function of the relative abundances for all species accounting for the top 80% abundance for each depth (rows) and timepoint (columns) for Angelo. Species with detected activity (AFE > 0) are colored by replicate (red for A, cyan for B, and magenta for C) and AFE values are indicated with transparency ranging from 0 – completely transparent to 1 – no transparency, normalized from 0 to the maximal value for a given combination of time and depth. Both axes are logarithmic.*

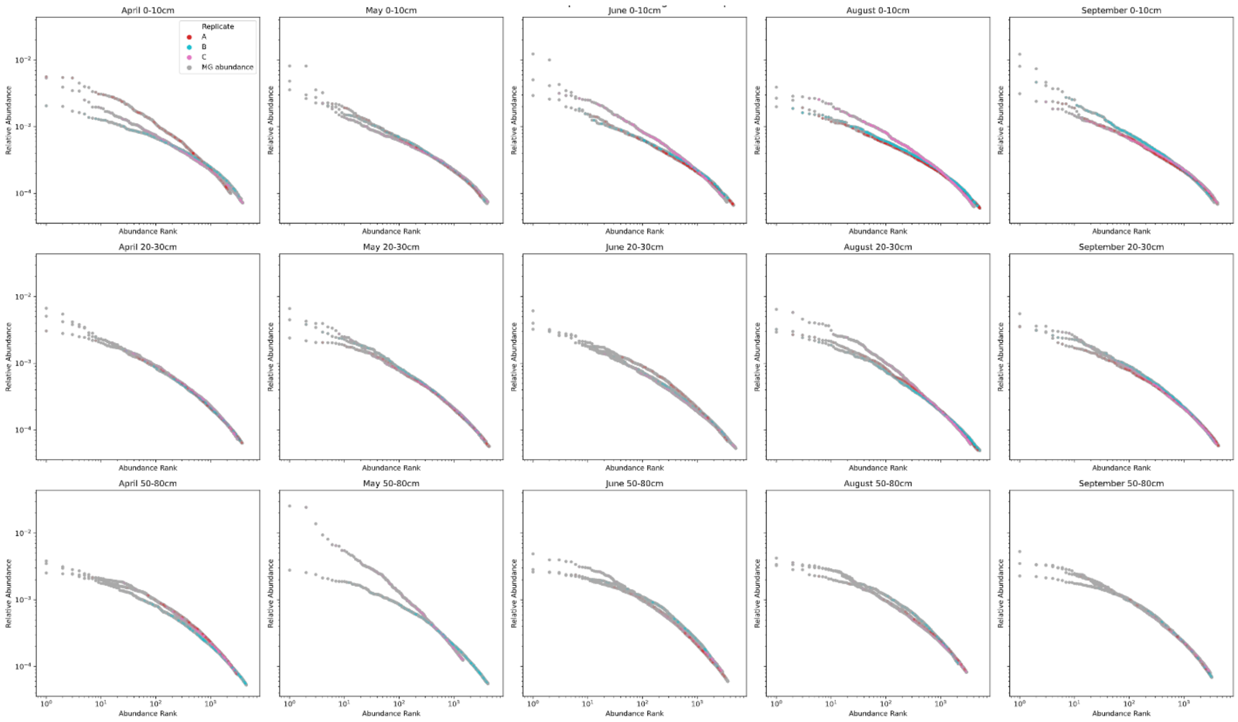


*Figure 6. Abundance ranks as a function of the relative abundances for all species accounting for the top 80% abundance for each depth (rows) and timepoint (columns) for Hopland. Species with detected activity (AFE > 0) are colored by replicate (red for A, cyan for B, and magenta for C) and AFE values are indicated with transparency ranging from 0 – completely transparent to 1 – no transparency, normalized from 0 to the maximal value for a given combination of time and depth. Both axes are logarithmic.*

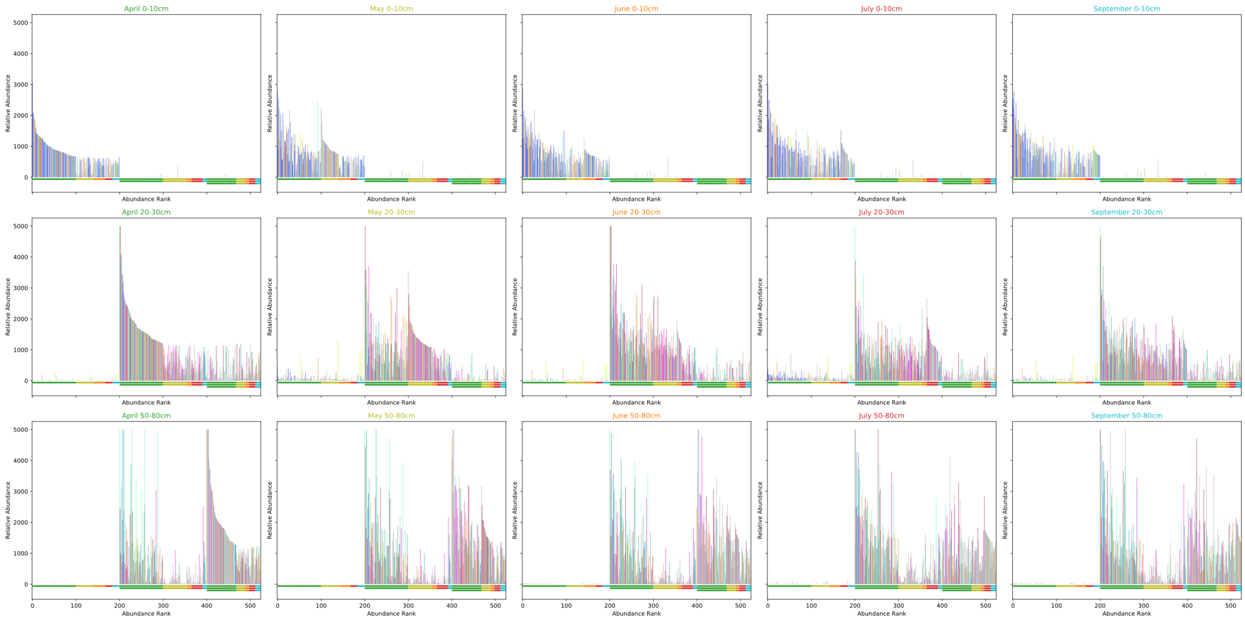


*Figure 7. Rank abundances of the top 100 species in the T0 reads for Angelo replicate A for each depth (rows) and timepoint (column). Relative abundance (tpm) is displayed on the y-axis as a column for each species. The position of each species is determined by descending order of their abundance rank starting from the upper left subplot. This means that if a species appears as rank 1 in the upper left subplot (April 0-10cm) it will continue to show up as rank 1 in the other subplots, even if its relative abundance is lower in the specific subplot. Therefore, all top 100 species for shallow soils also show up on plots of the deep soils and vice versa. Below the x-axis ranks corresponding to different timepoints are indicated with a colored line – green for April, yellow for May, orange for June, red for July, and cyan for September. Different depths are indicated with additional lines (one line for 0-10cm, two for 20-30cm, and three for 50-80cm).*

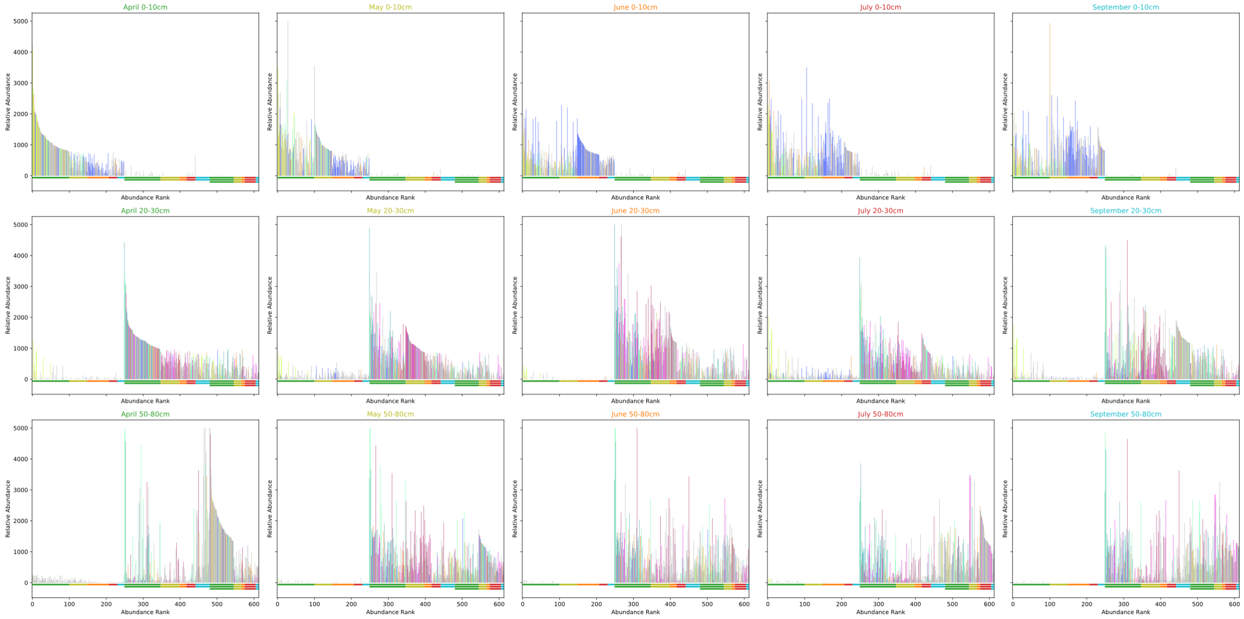


*Figure 8. Rank abundances of the top 100 species in the T0 reads for Angelo replicate B. Figure caption same as Figure 7.*

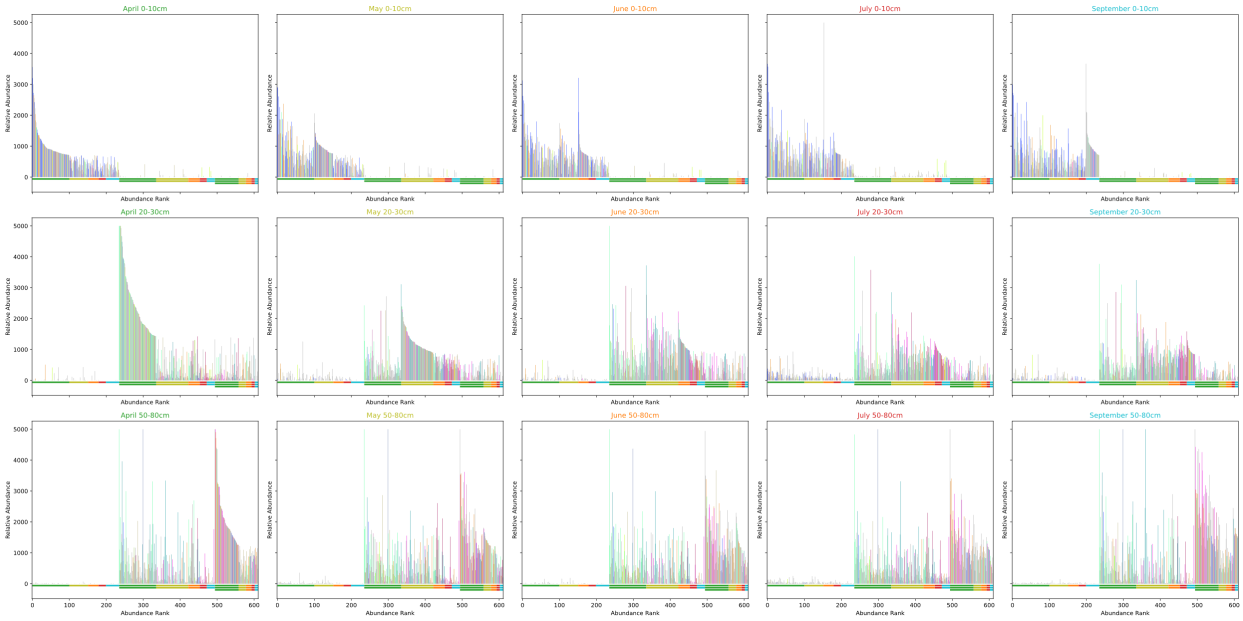


*Figure 9. Rank abundances of the top 100 species in the T0 reads for Angelo replicate C. Figure caption same as Figure 7.*

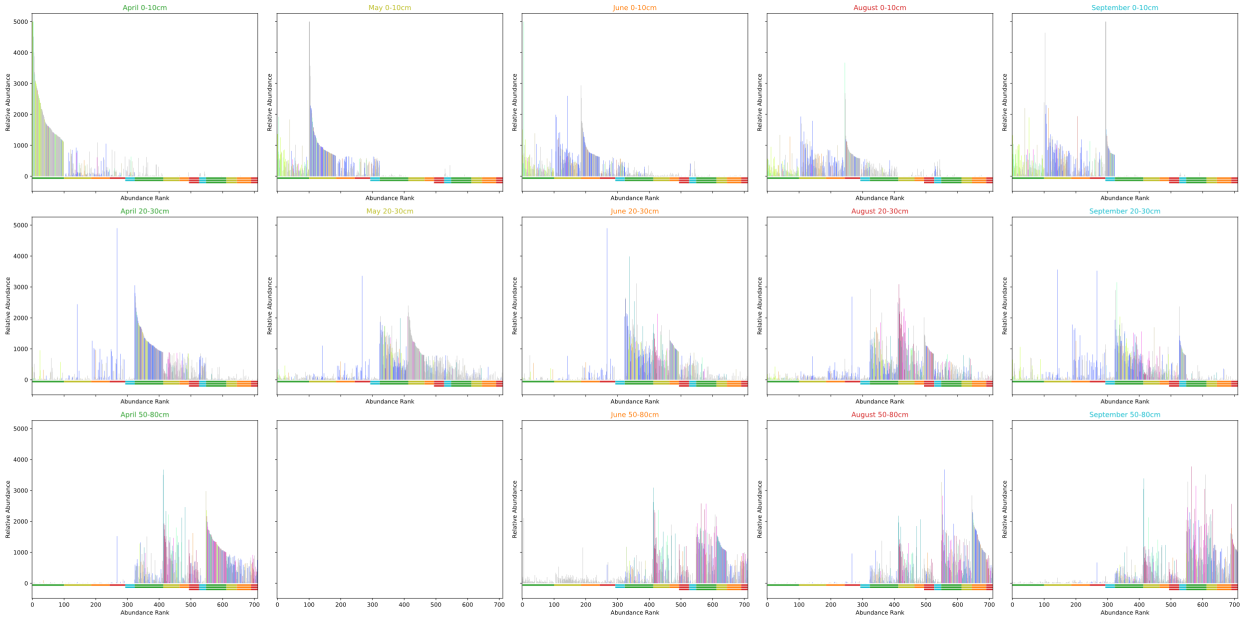


*Figure 10. Rank abundances of the top 100 species in the T0 reads for Hopland replicate A. Figure caption same as Figure 7.*

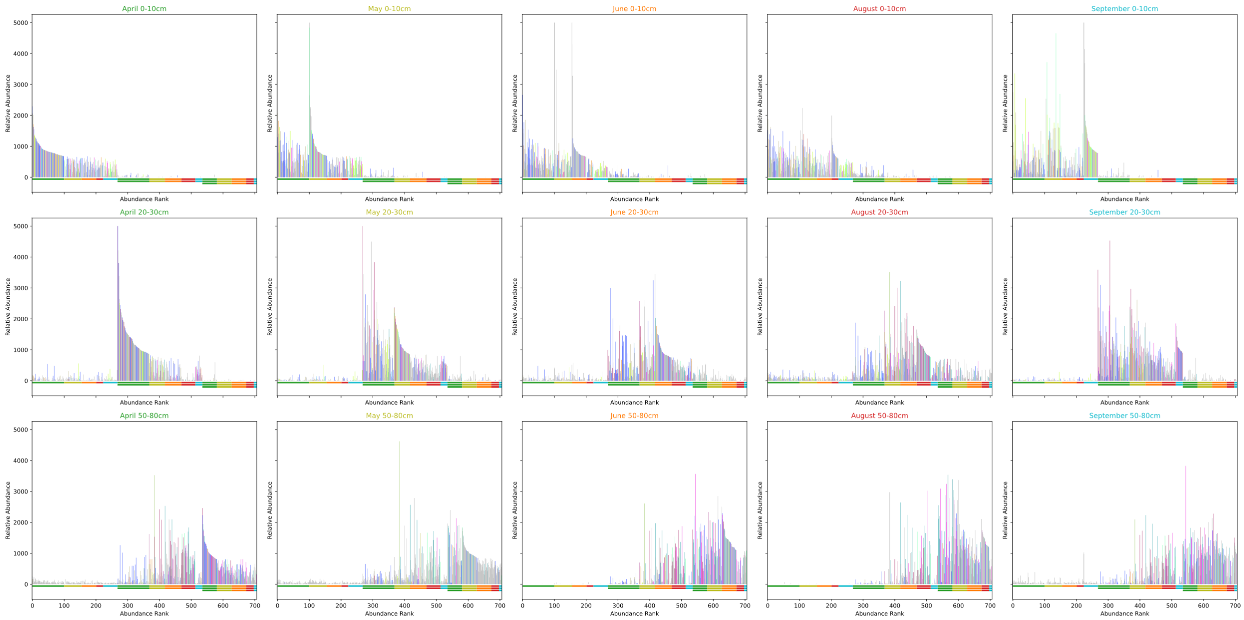


*Figure 11. Rank abundances of the top 100 species in the T0 reads for Hopland replicate B. Figure caption same as Figure 7.*

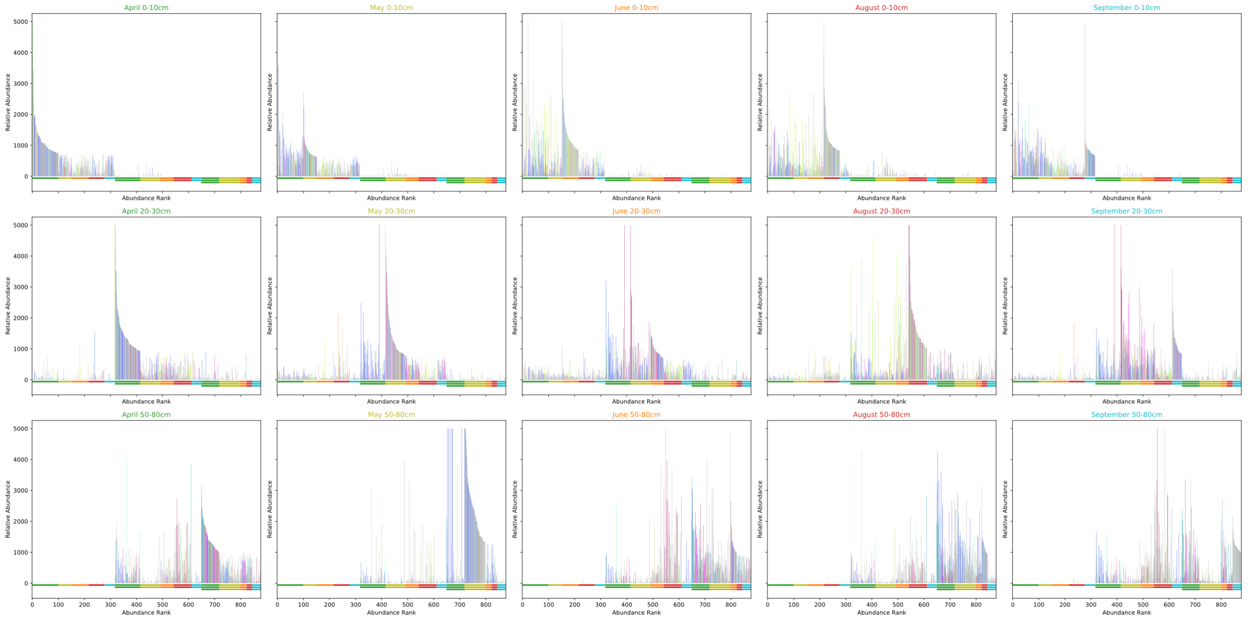


*Figure 12 Rank abundances of the top 100 species in the T0 reads for Hopland replicate C. Figure caption same as Figure 7.*

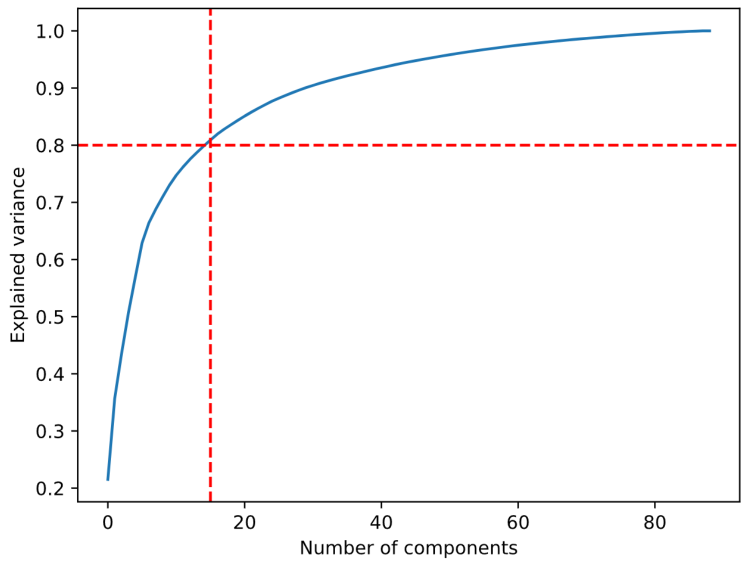


*Figure 13. Determination of number of components for PCA, used to transform species data for the CCA from main text Fig. 1. Transformations were performed with the sklearn python library (v. 1.3.2). The number of components determined by an 80% threshold is 15.*

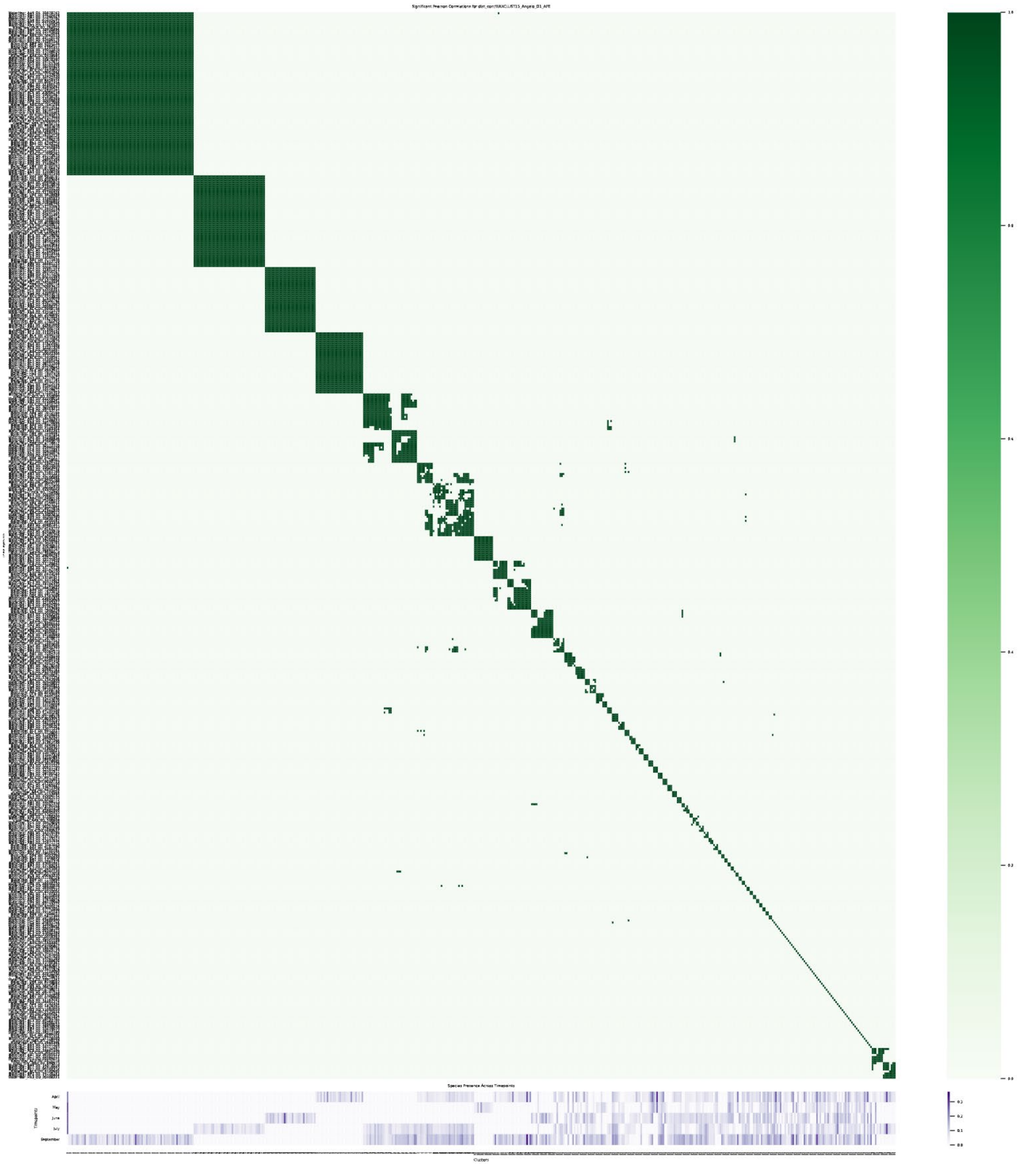


*Figure 14. All vs. all comparison of AFE changes over time for Angelo species at depth 0-10cm. The top heatmap shows the significant correlations between each pair of species calculated as the covariance over the standard deviation and controlled for multiple testing with Benjamini/Hochberg method. Bottom heatmap shows the activity of each species for each timepoint with April being on top and September on the bottom. Species are ordered by hierarchical clustering with the ward method and maximum number of clusters set to 15 with the maxclust from the scipy python package (v. 1.10.1). Clusters defined here are labeled 'cohorts' or 'seasonal cohorts'.*

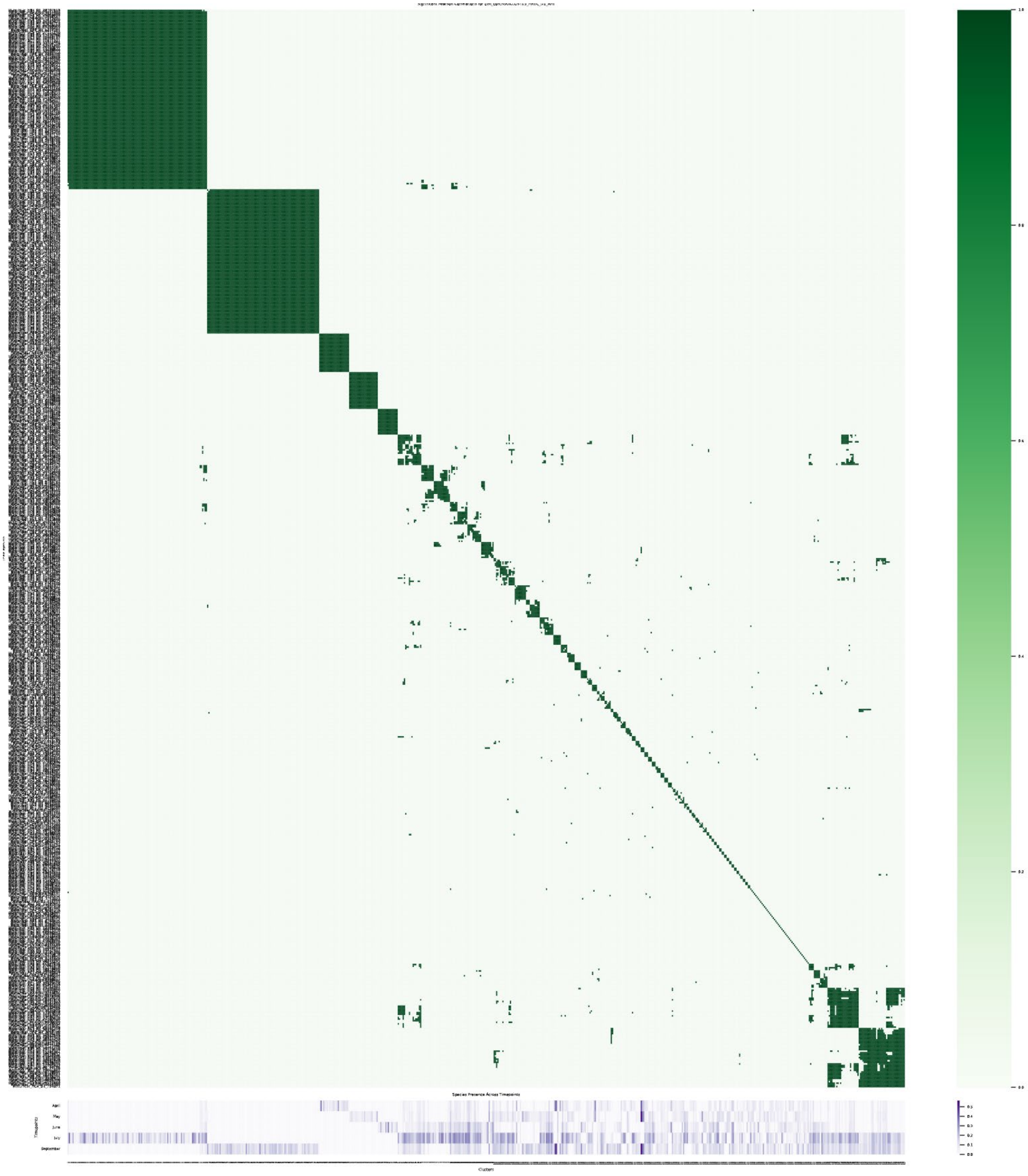


*Figure 15. All vs. all comparison of AFE changes over time for Hopland species at depth 0-10cm. The top heatmap shows the significant correlations between each pair of species calculated as the covariance over the standard deviation and controlled for multiple testing with Benjamini/Hochberg method. Bottom heatmap shows the activity of each species for each timepoint with April being on top and September on the bottom. Species are ordered by hierarchical clustering with the ward method and maximum number of clusters set to 15 with the maxclust from the scipy python package (v. 1.10.1). Clusters defined here are labeled 'cohorts' or 'seasonal cohorts'.*

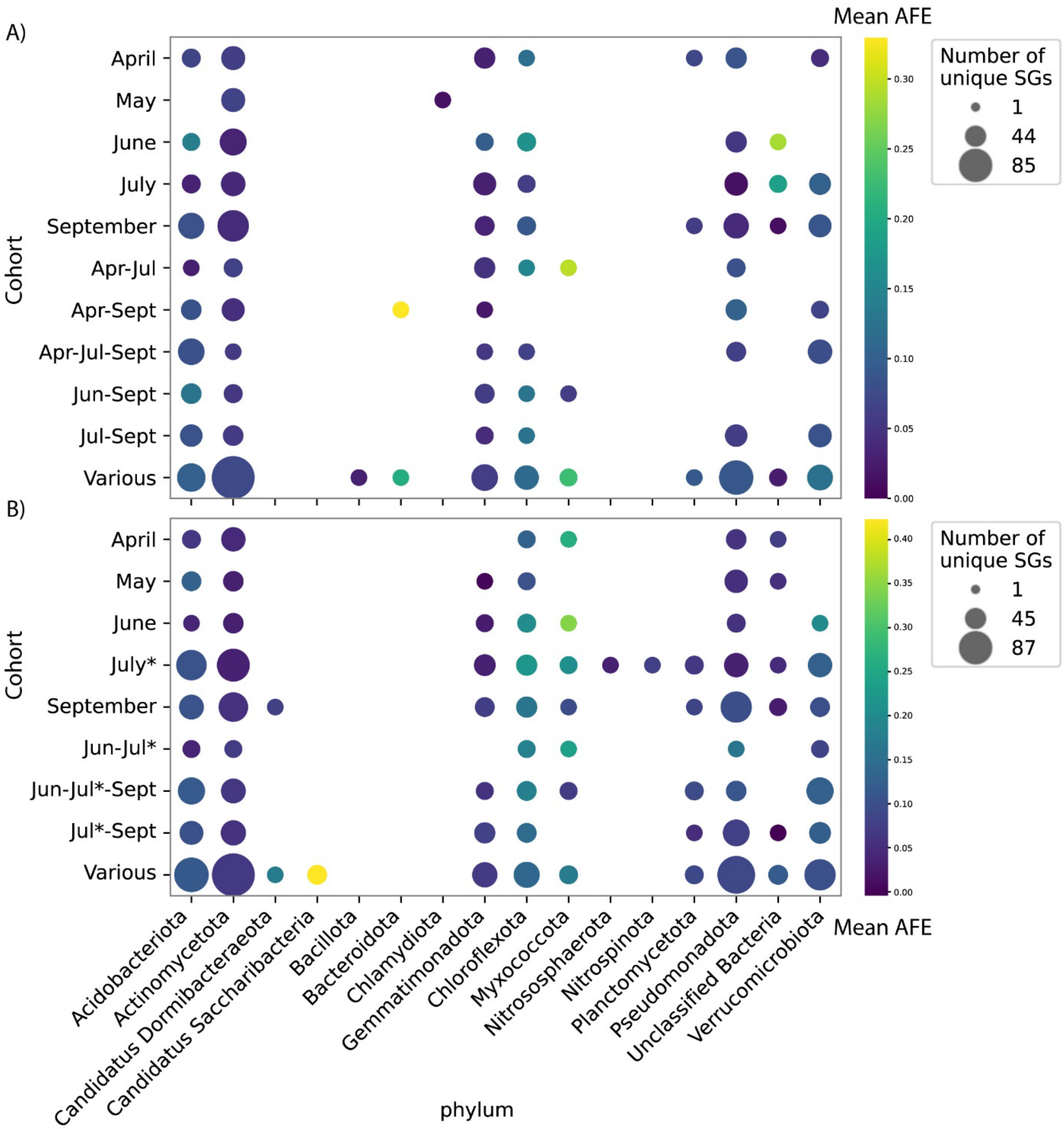


*Figure 16. Phylogenetic composition for each cluster between A) Angelo and B) Hopland. Average AFE values for each cohort are shown with colors and the number of unique species are shown with the size of the datapoints. * For clarity when comparing A) and B) the Hopland `September before rain` timepoint is labeled as July.*

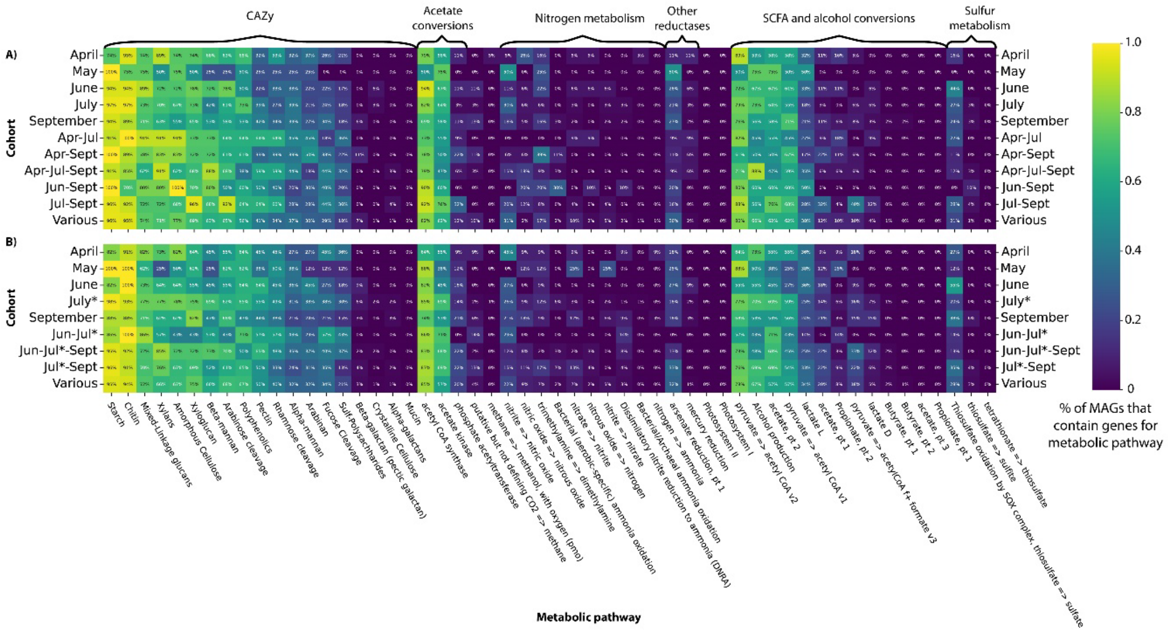


*Figure 17. Functional category composition for each cluster between A) Angelo and B) Hopland. Categories are determined based on DRAM results for genomes present in each cohort. The % of genomes containing each function is indicated with color. * For clarity when comparing A) and B) the Hopland `September before rain` timepoint is labeled as July.*

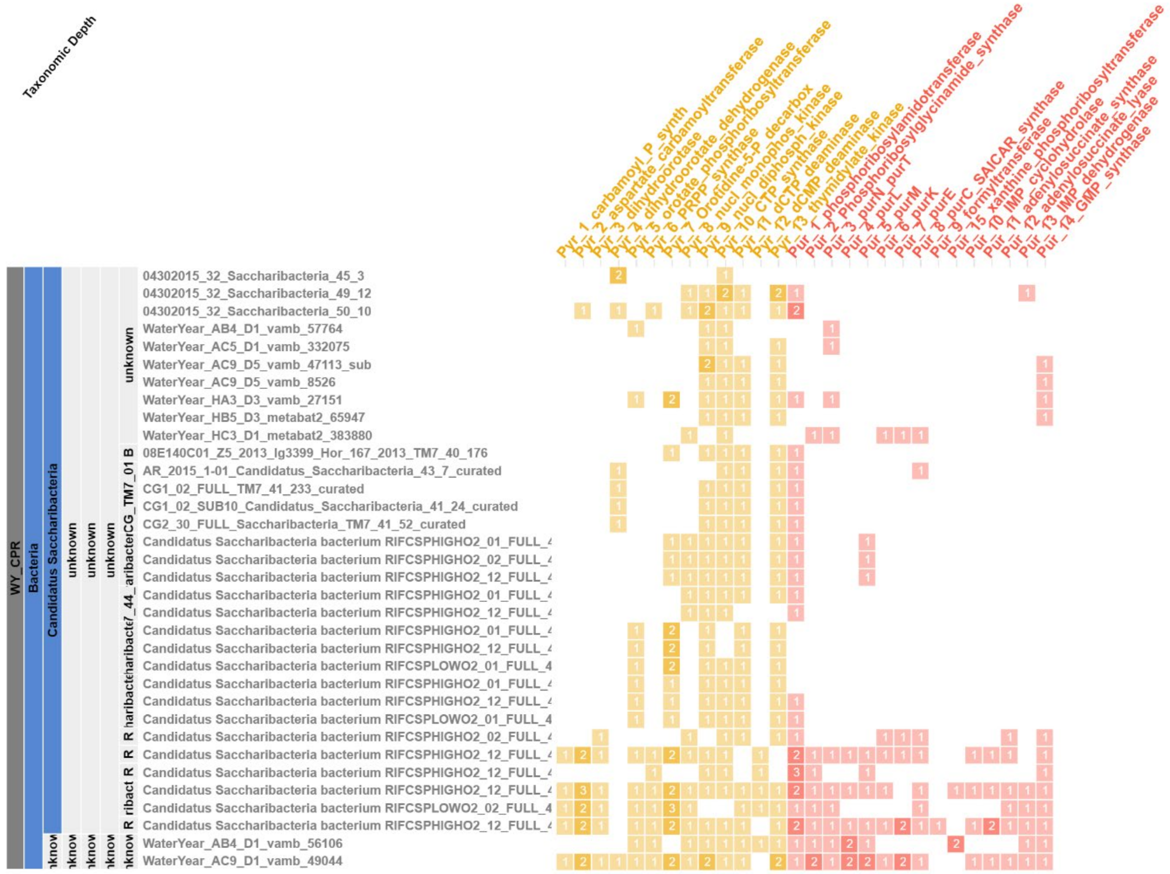


*Figure 18. Nucleotide synthesis capabilities of Saccharibacteria from the Water year project and other projects in ggKbase. The two pathways for pyrimidine and purine synthesis are indicated with yellow and red color respectively. The number of genes for each step are indicated with number within each cell.*

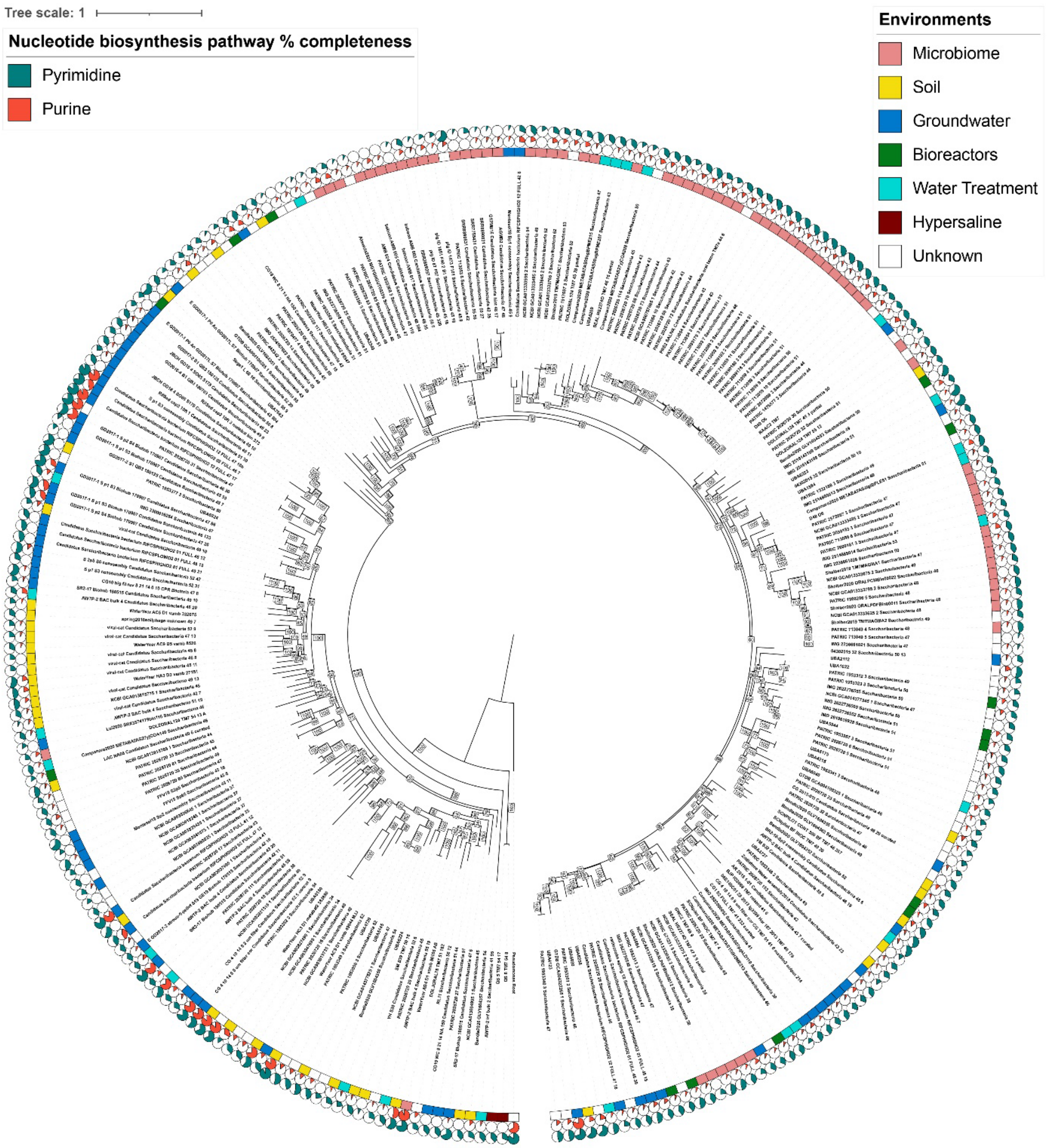


*Figure 19. Detailed phylogenetic tree of Saccharibacteria sequences used in main text Fig. 5. Completeness of the purine and pyrimidine synthesis pathways are indicated with red and green pie charts for each leaf. The environmental source when available is indicated with colored rectangle at each leaf. Bootstrap values are indicated at each branching point.*

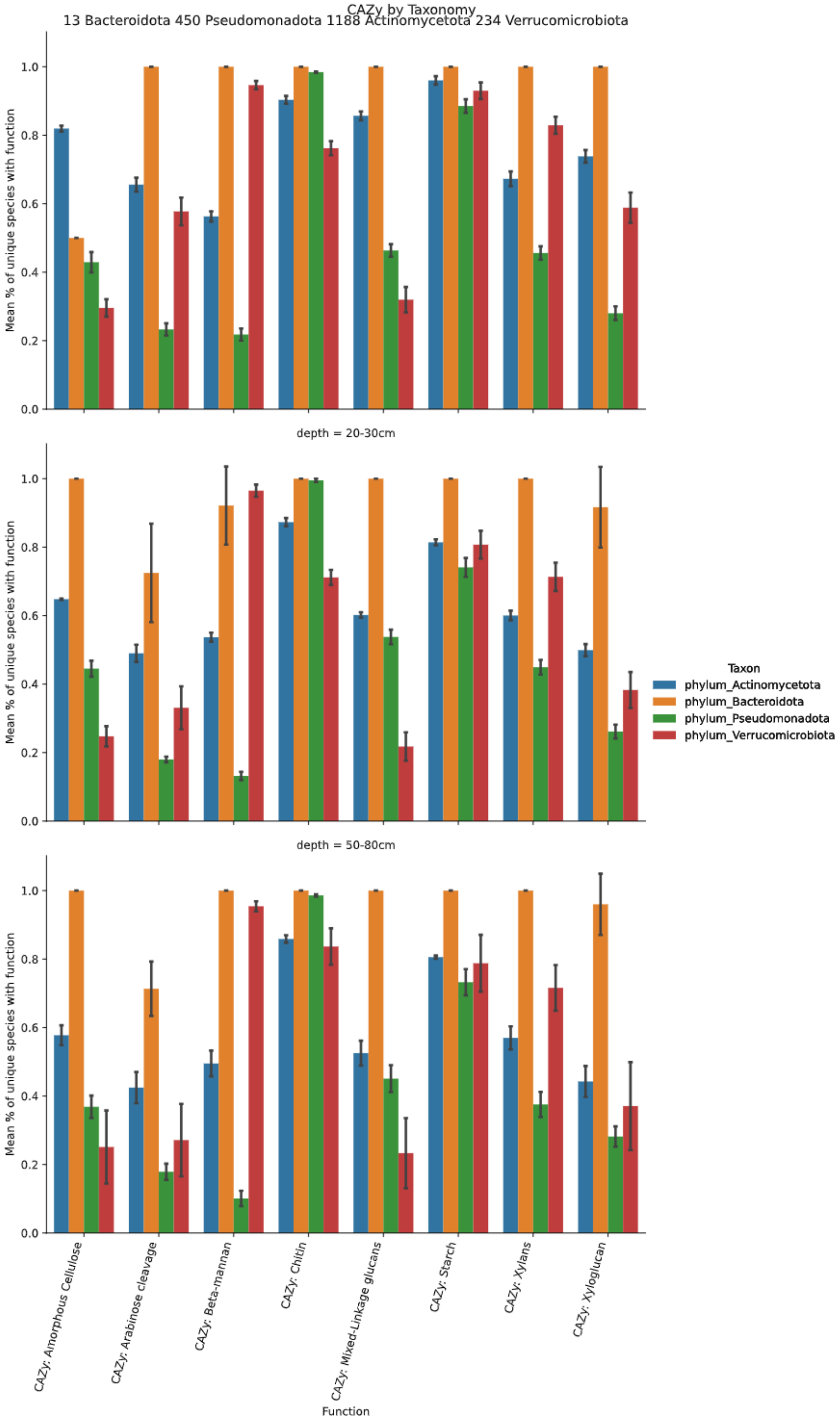


*Figure 20. Percent of species within four bacterial phyla with a specific CAZy function present at all depths (rows). The total species for Bacteroidota are 13 (orange), for Pseudomonadota are 450 (green), for Actinomycetota are 1188 (blue), and for Verrucomicrobiota are 234 (red).*

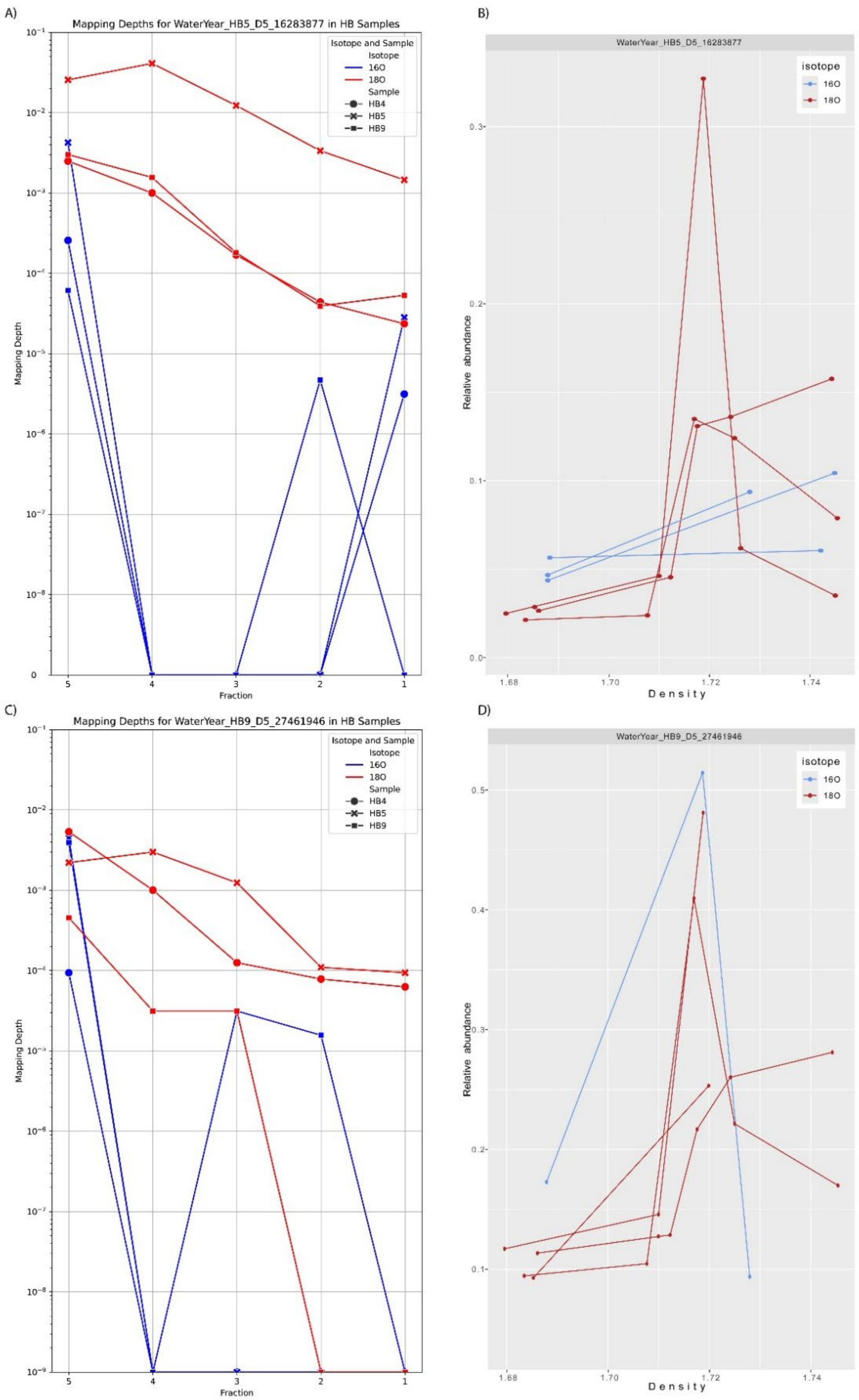


*Figure 21. Atom enrichment for the NCLDV (A) and B)) and its probable eukaryotic host (C) and D)). A) and C) show the raw read numbers mapping to the contigs for isotope and sample and different timepoints are shown with shapes. B) and D) show the relative abundance, which is calculated by the number of reads normalized by DNA amount within each fraction. Sequences were present only in replicate B.*

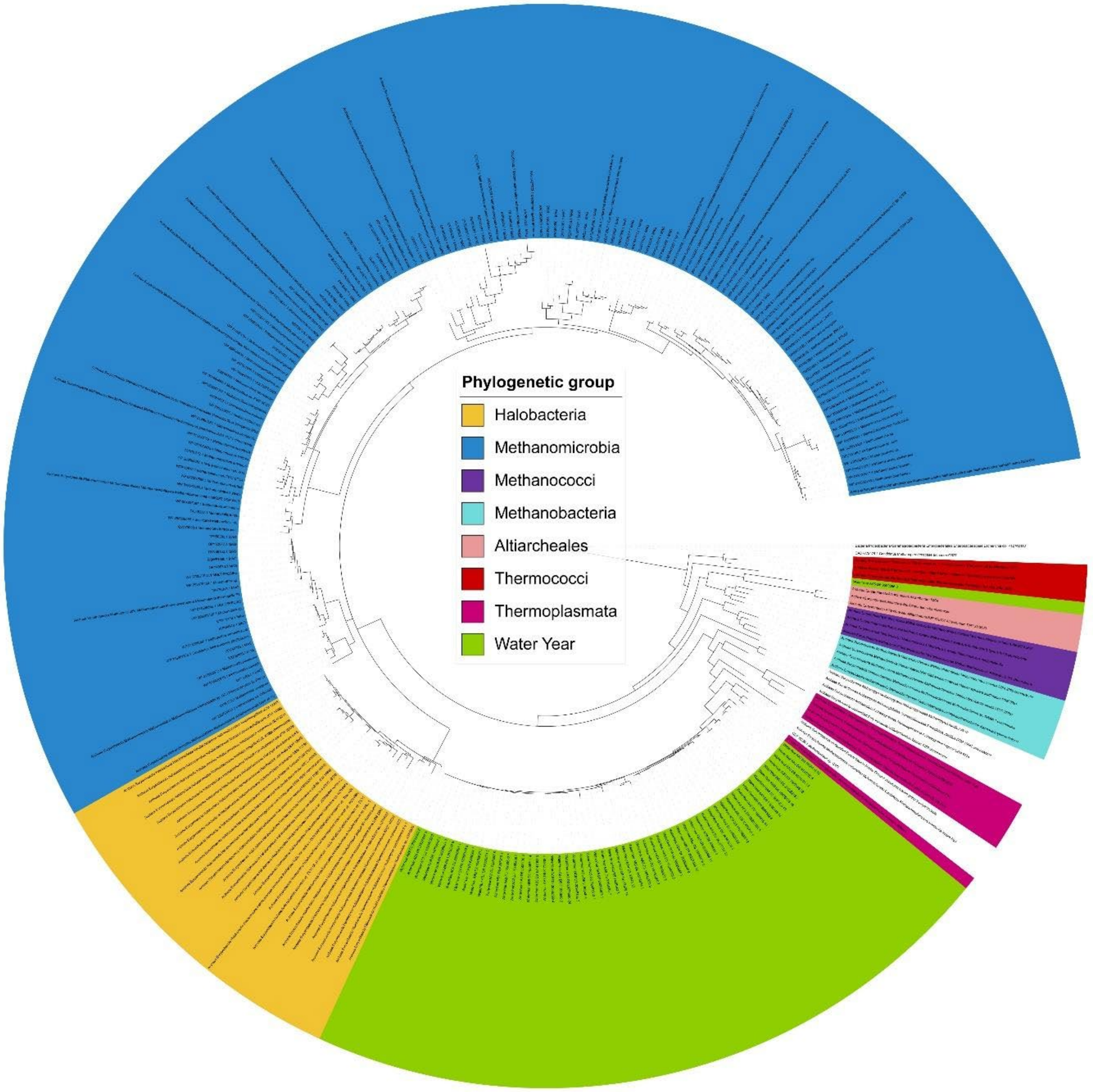


*Figure 22. Phylogenetic tree of rpS3 sequences from Euryarchaeota species present in the Water Year samples and reference sequences from NCBI. Different clades from the reference sequences are colored and samples from our project are colored green.*

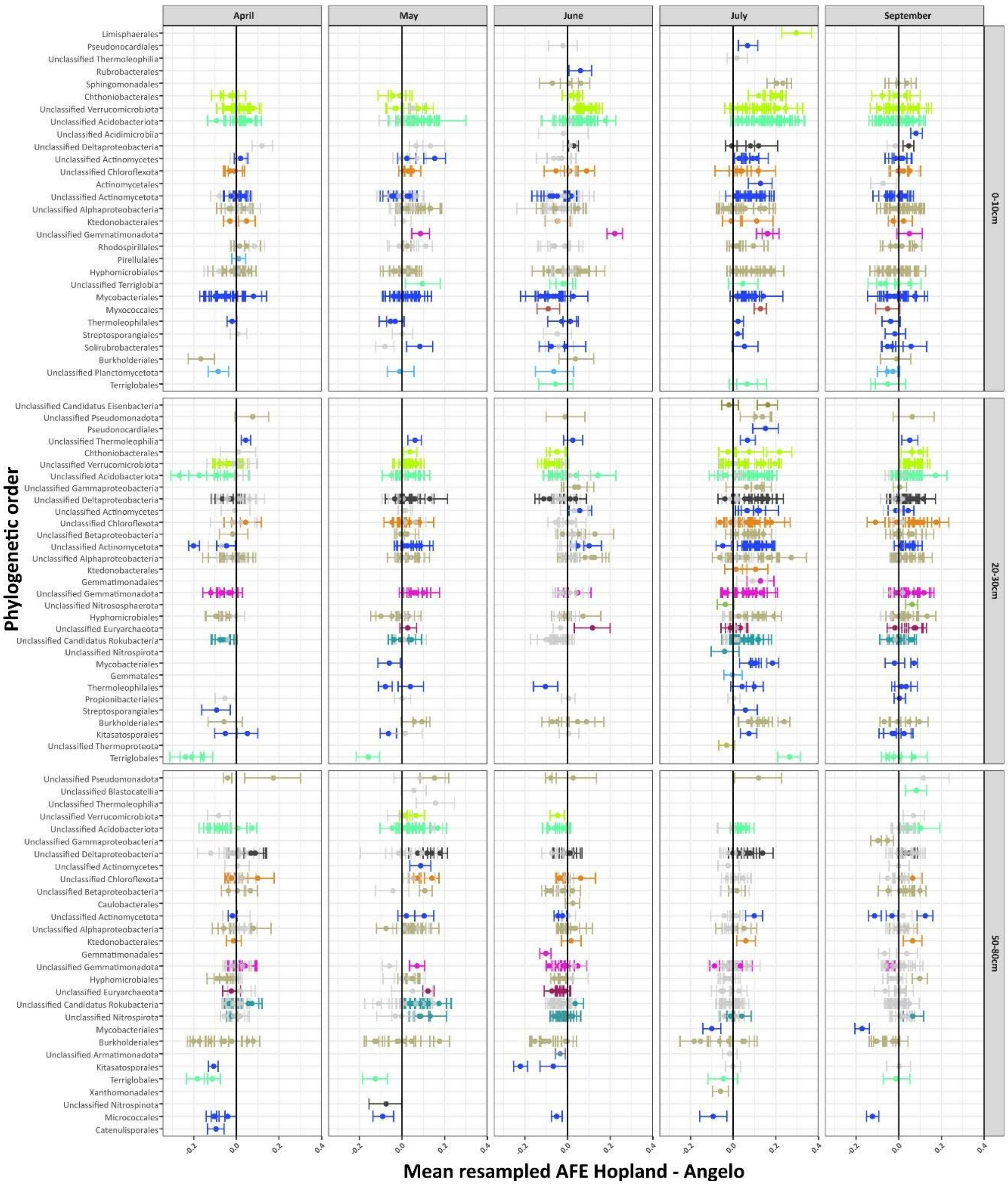


*Figure 23. Difference of AFE between organisms present in both Angelo and Hopland. Negative values indicate higher activity in Angelo, positive – higher activity in Hopland, and points close to the 0 – similar activity between Angelo and Hopland. Data are colored by phylum when the confidence interval of the original activity value is above 0. The confidence intervals depicted here are the ones for the ecosystem where greater activity is detected.*

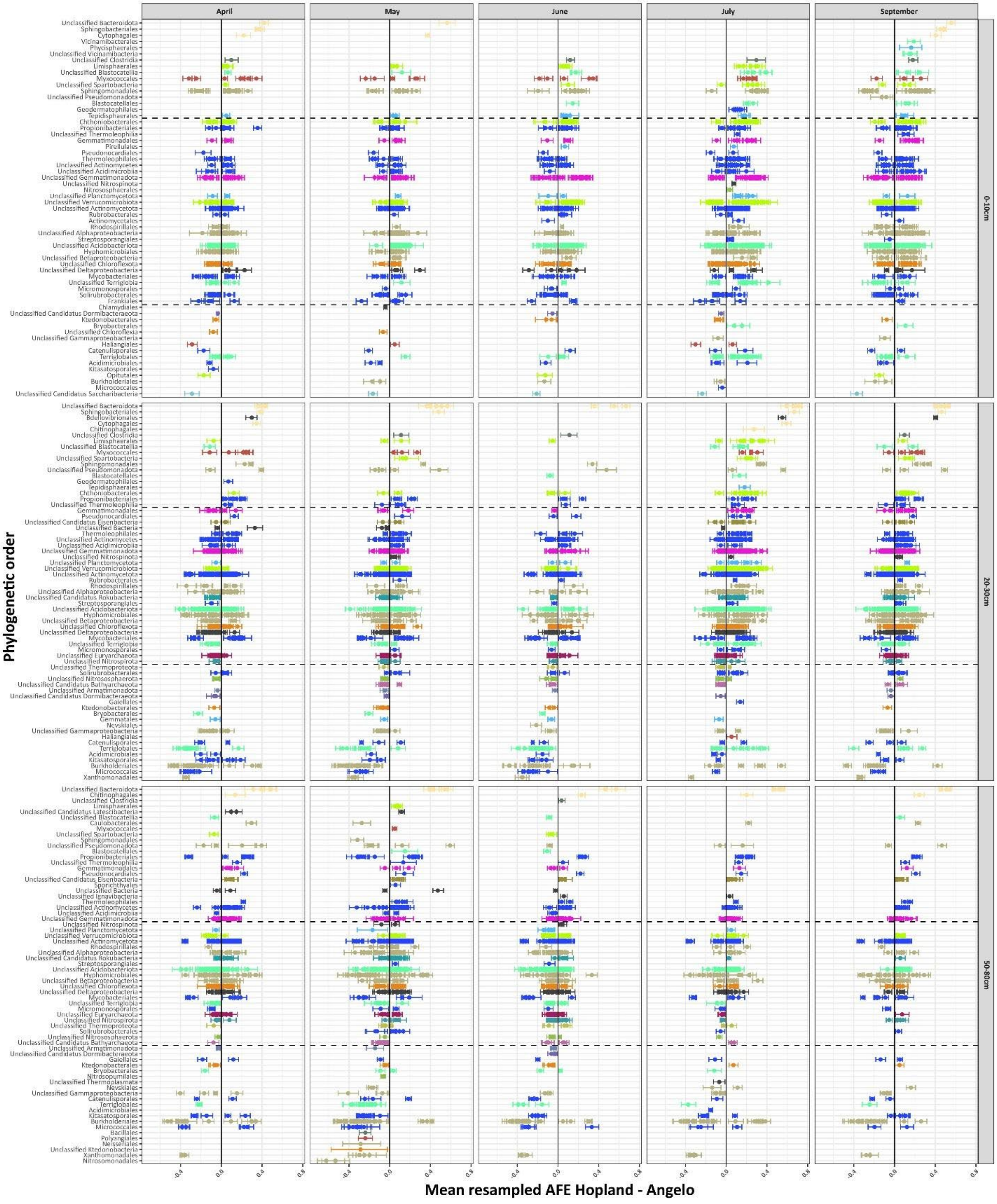


*Figure 24. AFE results for organisms only present in either Angelo or Hopland. Negative values indicate activity in Angelo. Dashed lines broadly categorize each plot in top (Hopland more active), middle (both ecosystems have equally active representatives from that genus), and bottom (Angelo more active) groups.*

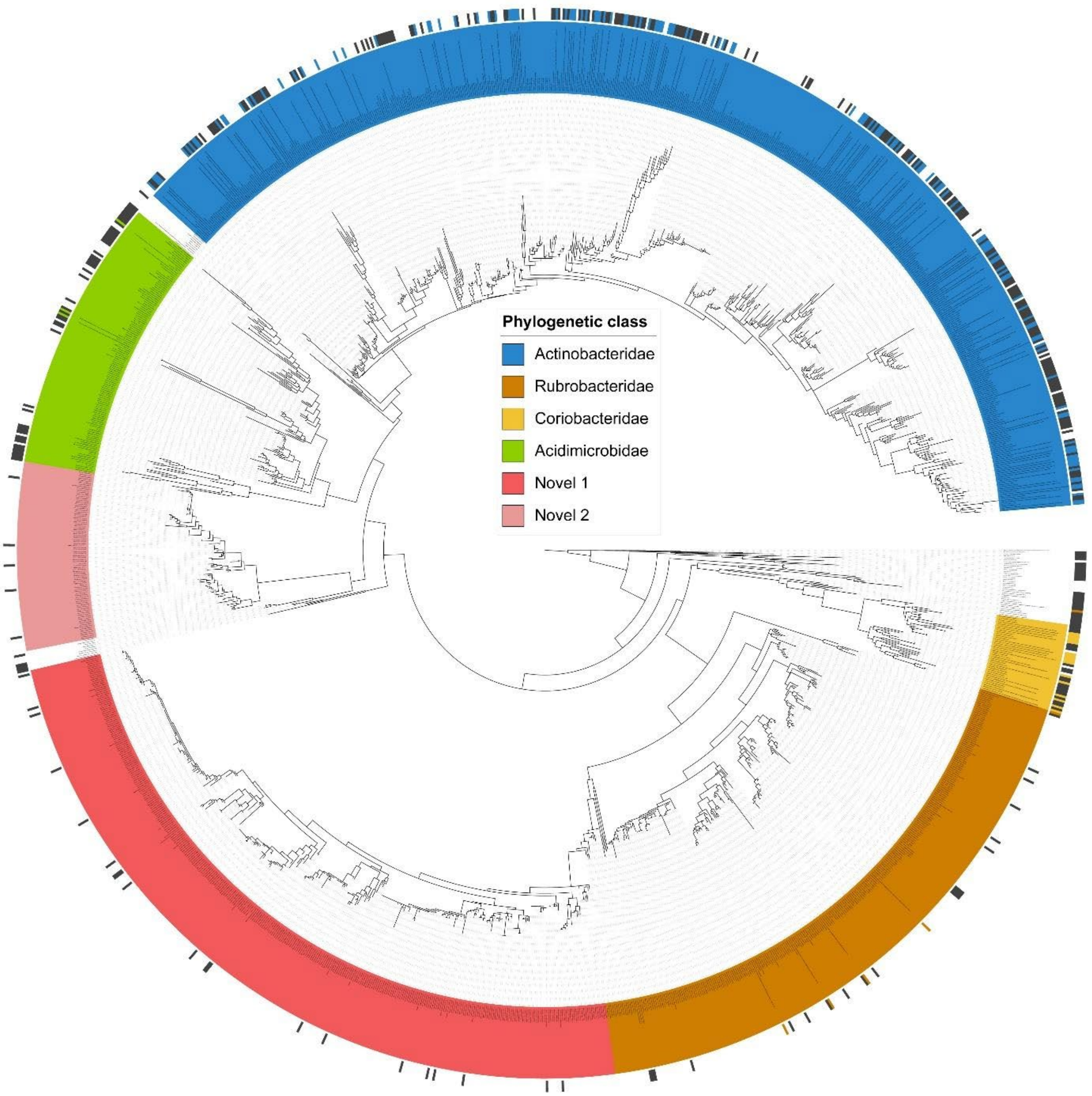


*Figure 25. Phylogenetic tree of rpS3 sequences from Actinomycetota species present in the Water Year samples and reference sequences from NCBI. Different clades were manually assigned based on reference presence and tree topology. Colored rectangles at the outer edge of the tree indicate references with known phylogenetic classification and gray rectangles indicate references with unknown phylogenetic classification.*

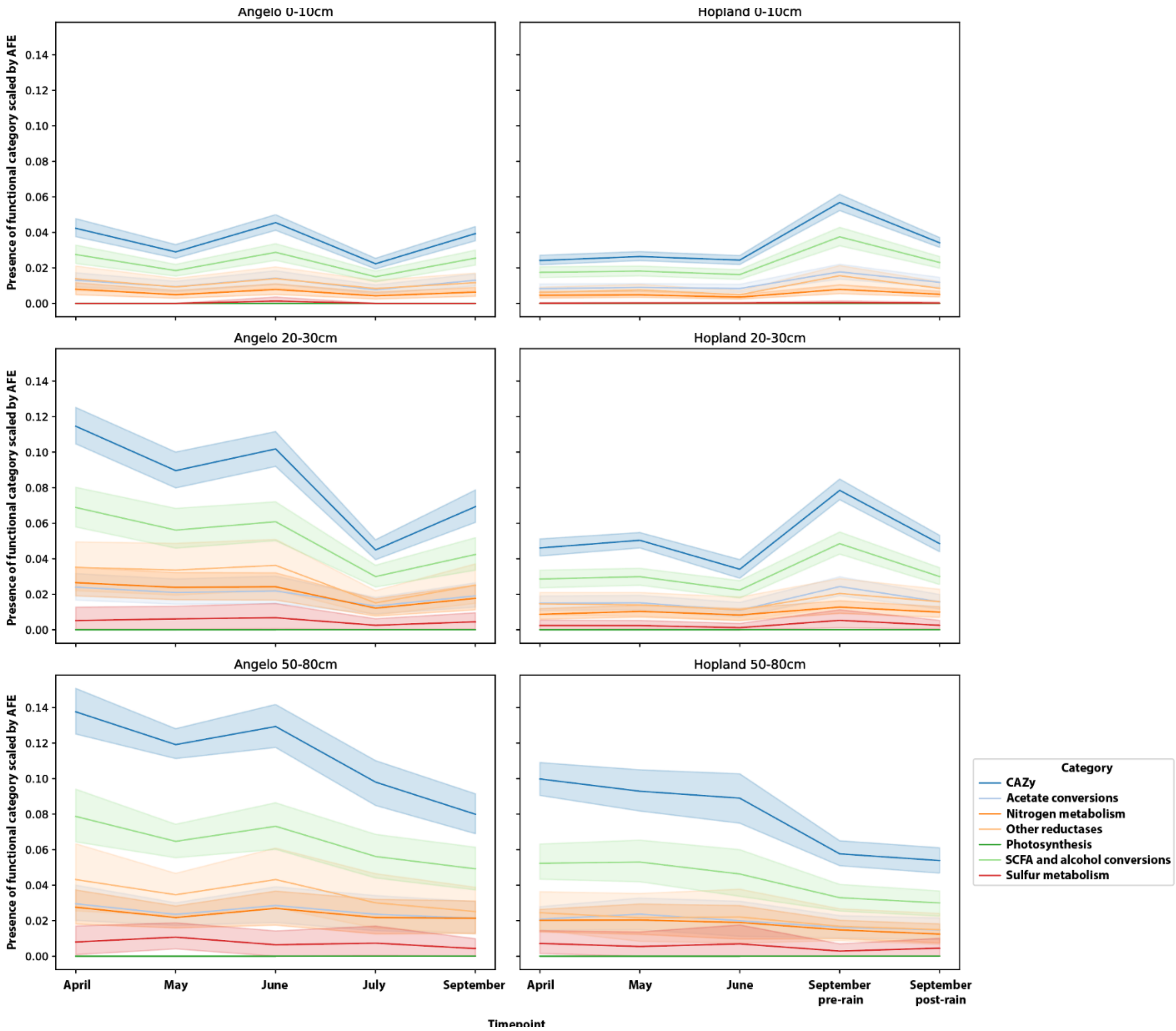


*Figure 26. Presence of metabolism categories within active Actinobacteridae for each ecosystem (column) and depth (row). The y-axis represents a scaled presence of functional category by the AFE for each species. Confidence intervals are based on the variability within the AFE values between species at 95%.*

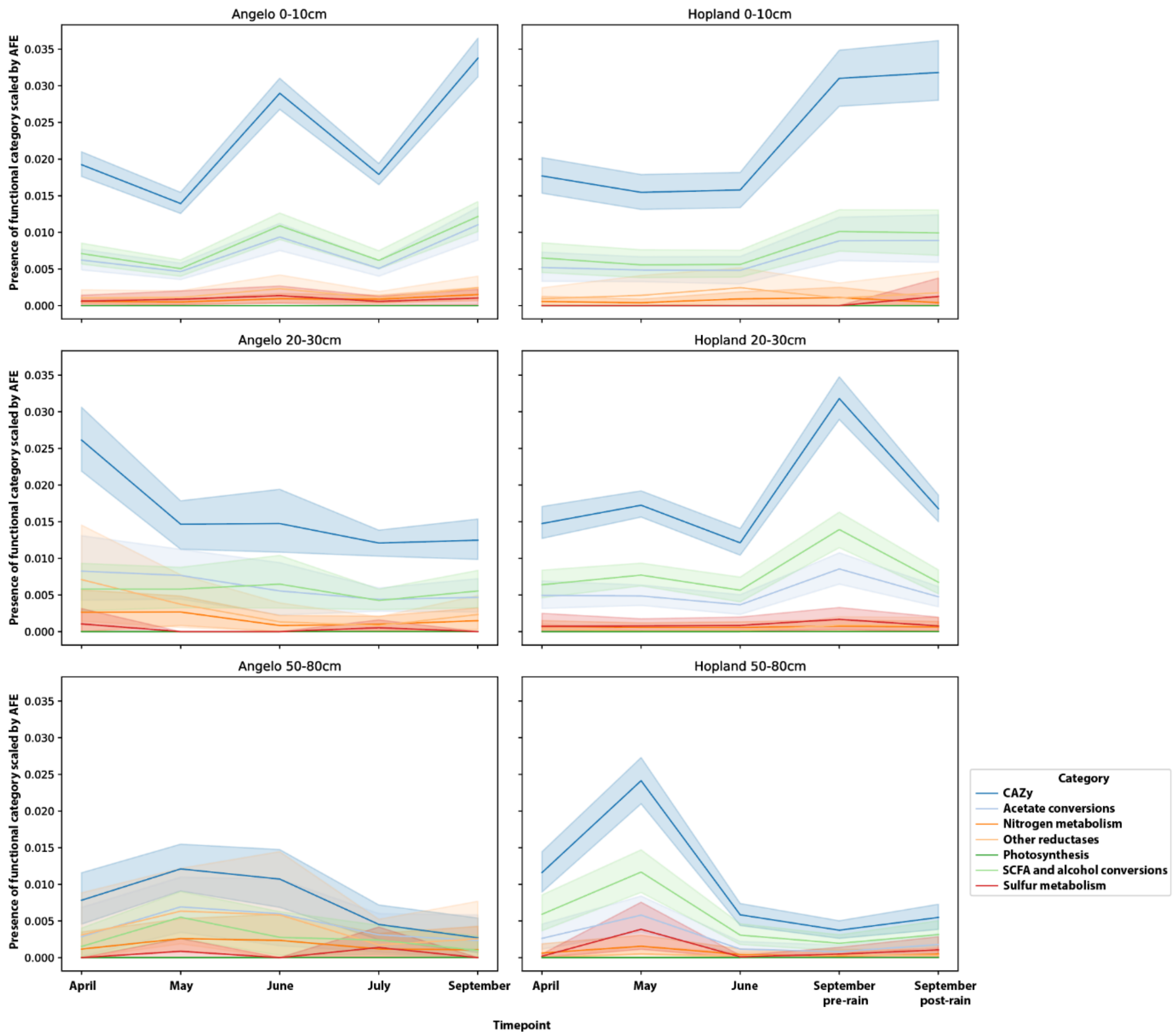


*Figure 27. Presence of metabolism categories within active Rubrobacteridae for each ecosystem (column) and depth (row). The y-axis represents a scaled presence of functional category by the AFE for each species. Confidence intervals are based on the variability within the AFE values between species at 95%.*

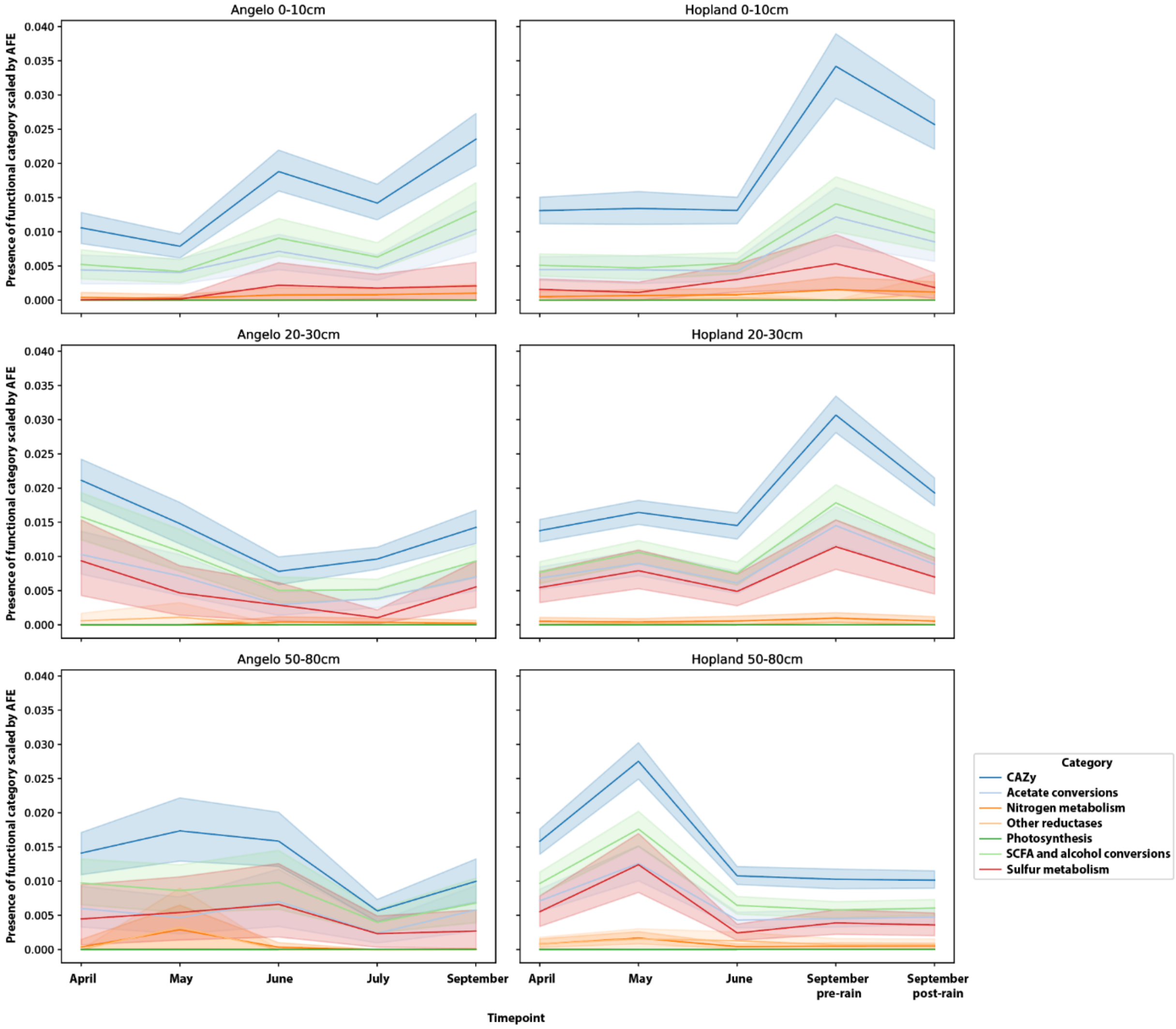


*Figure 28. Presence of metabolism categories within active Novel clade 1 species for each ecosystem (column) and depth (row). The y-axis represents a scaled presence of functional category by the AFE for each species. Confidence intervals are based on the variability within the AFE values between species at 95%.*

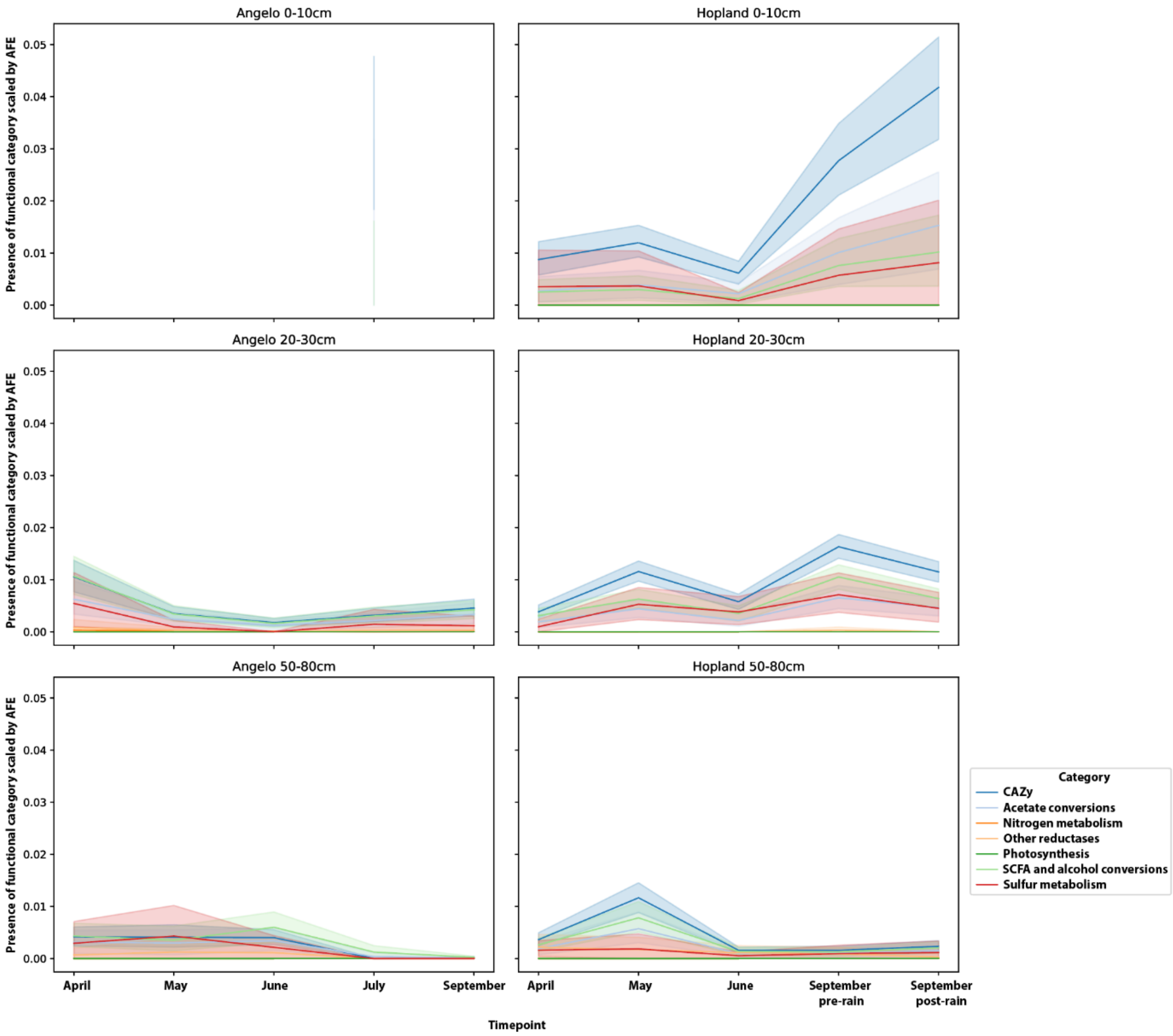


*Figure 29. Presence of metabolism categories within active Novel clade 2 species for each ecosystem (column) and depth (row). The y-axis represents a scaled presence of functional category by the AFE for each species. Confidence intervals are based on the variability within the AFE values between species at 95%.*

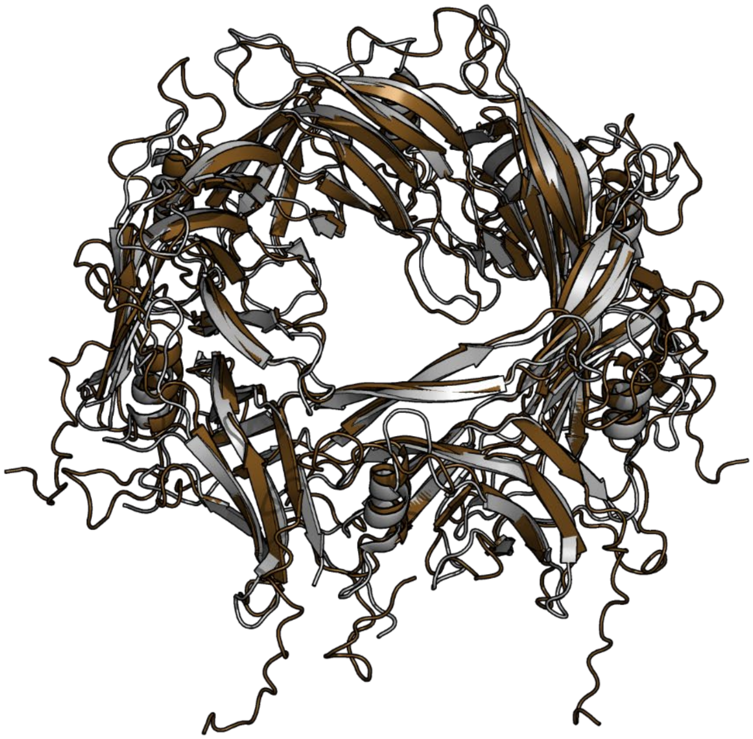

*Figure 30. Superimposition of Water Year Haliangiales six tube monomers (brown) and reference structures (gray). The PDB reference is 7B5H chains AD, BD, CD, DD, ED, and FD. Superimposition was performed with cealign from PyMOL, resulting in 2.5 RMSD for each monomer.*

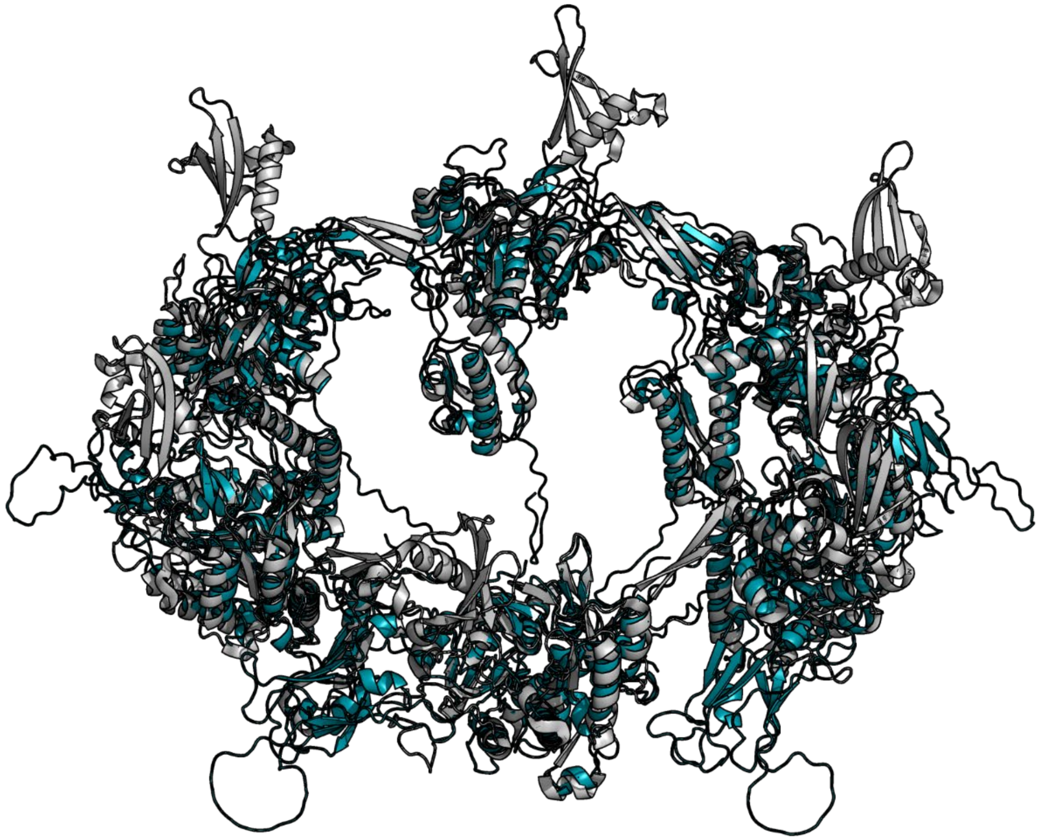

*Figure 31. Superimposition of Water Year Haliangiales six tube sheath monomers (cyan) and reference structures (gray). The PDB reference is 7B5H chains AC, BC, CC, DC, EC, and FC. Superimposition was performed with cealign from PyMOL, resulting in 4 RMSD for each monomer.*

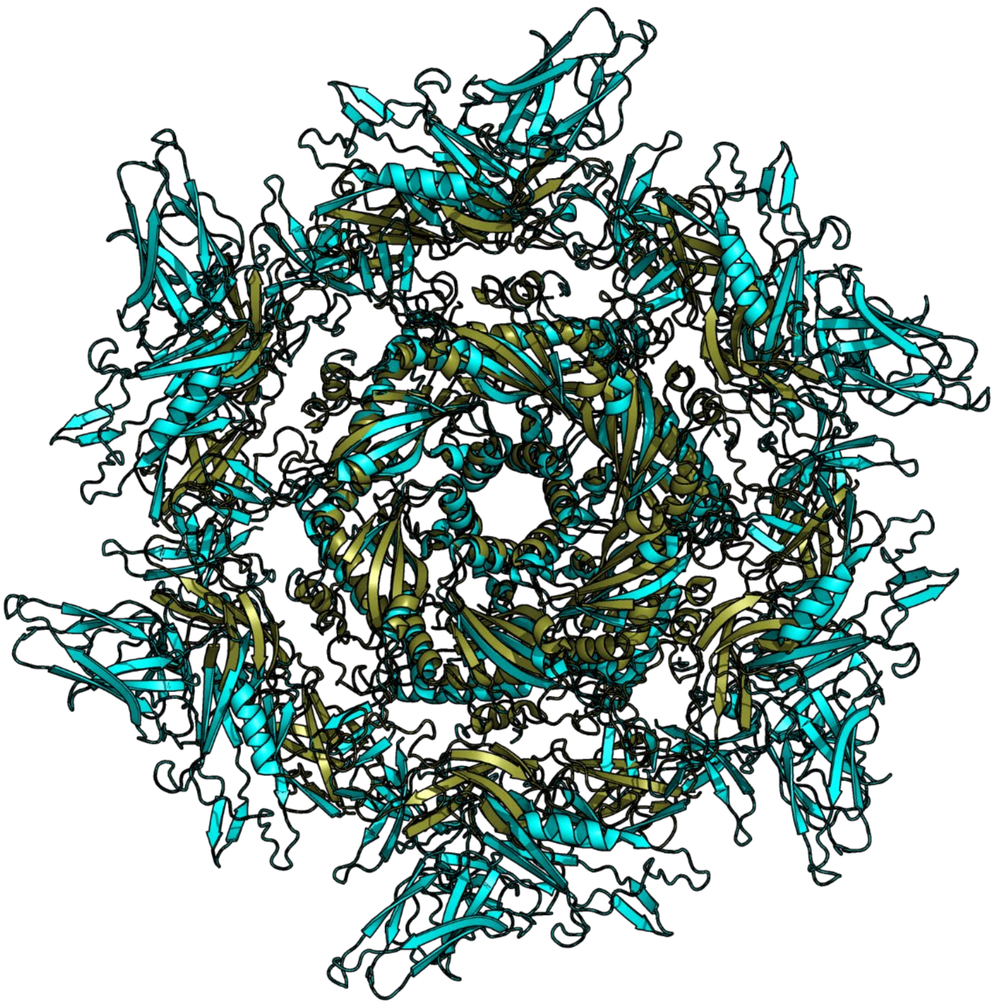

*Figure 32. Superimposition of Water Year Haliangiales six cap monomers (green) and reference structures (cyan). The PDB reference is 7B5I chains AA, BA, CA, DA, EA, and FA. Superimposition was performed with cealign from PyMOL, resulting in 3.7 RMSD for each monomer.*

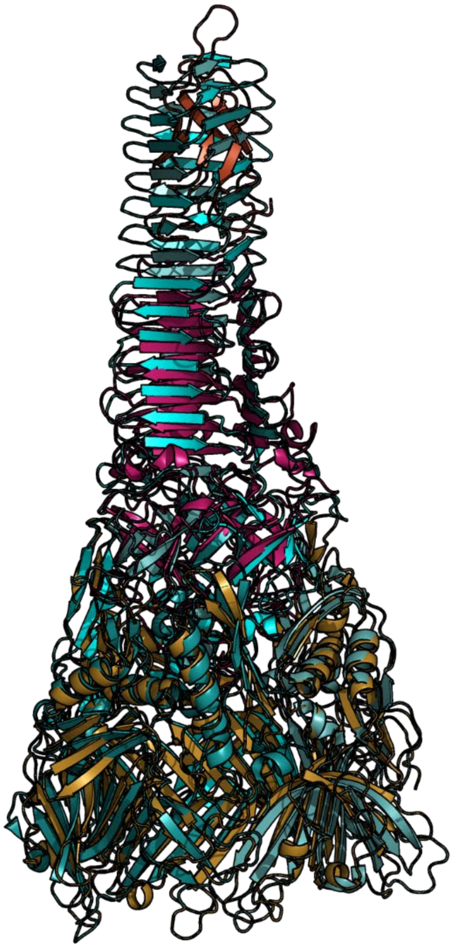

*Figure 33. Superimposition of Water Year Haliangiales spike (brown, violet, red) and reference structures (cyan). The PDB reference includes three monomers from 7B5H – chains AJ, CJ, and EJ. Superimposition was performed with cealign from PyMOL, resulting in 3.5 RMSD averaged over each superimposed monomer. The reference lacks a PAR tip polymer, which is present in our structure (red).*

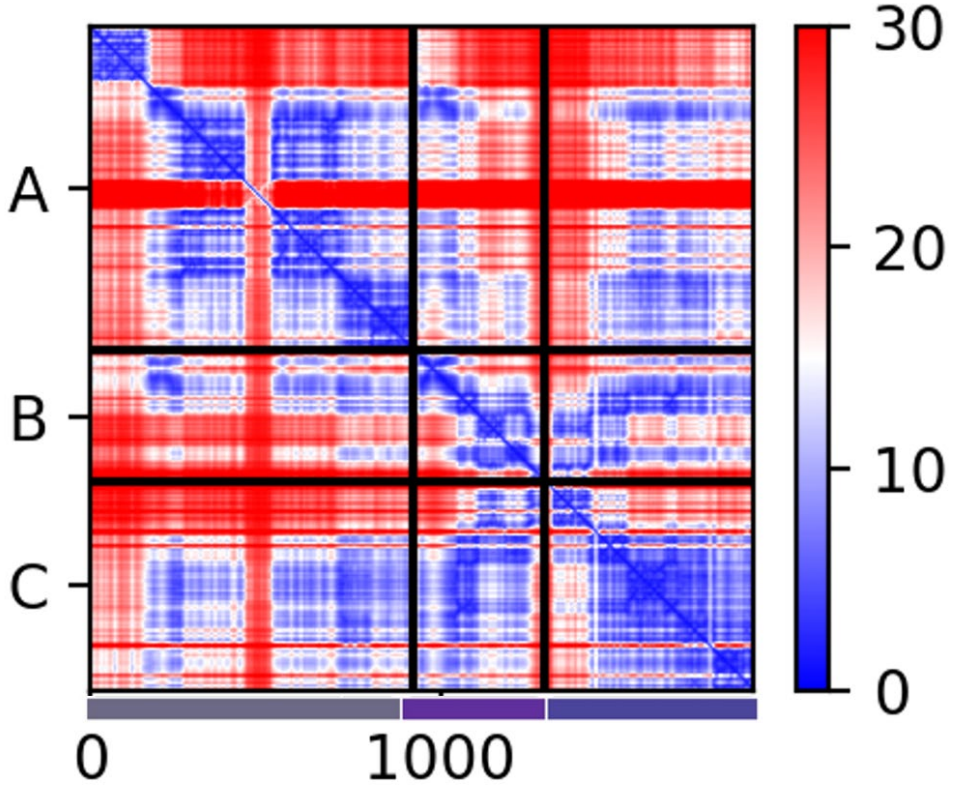


*Figure 34. Predicted alignment errors (PAE) generated for the heteromeric baseplate complex of the injection system. The three polymers are indicated with colors matching the main text Fig. 6 with a bar below the x-axis. Lower PAE is indicated with blue and means higher confidence in the interaction between residues. If there are blue regions outside the diagonal, between the polymer divisions this can mean the three proteins fold together.*

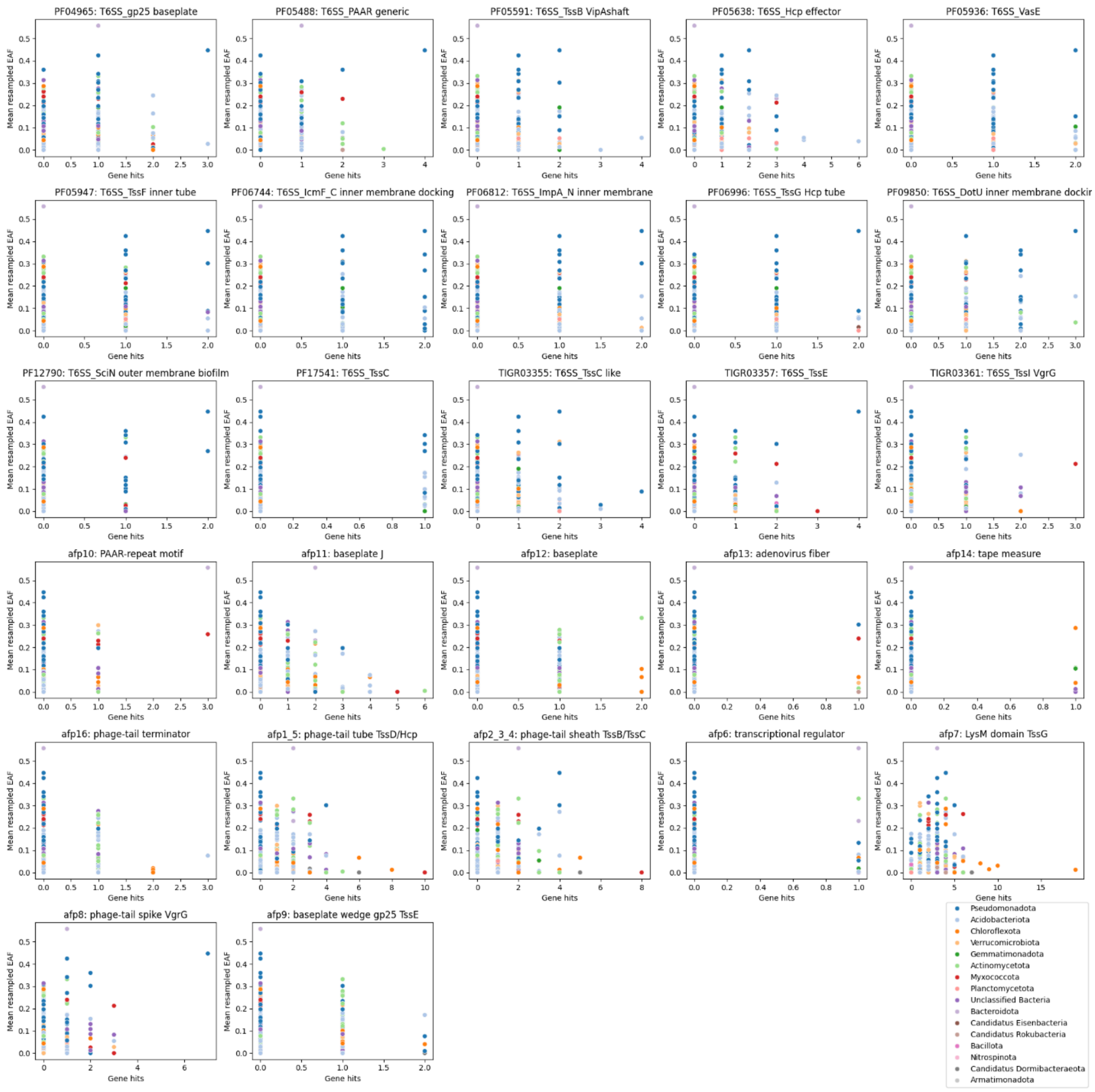


*Figure 35. The presence of T6SS and CIS genes as function of AFE in all genomes. The presence of the genes is detected with HMM. Shown results for genomes with at least 4 gene hits in either CIS (indicated with afp) or T6SS (hits from pfam or tigrfam indicated with T6SS) groups.*

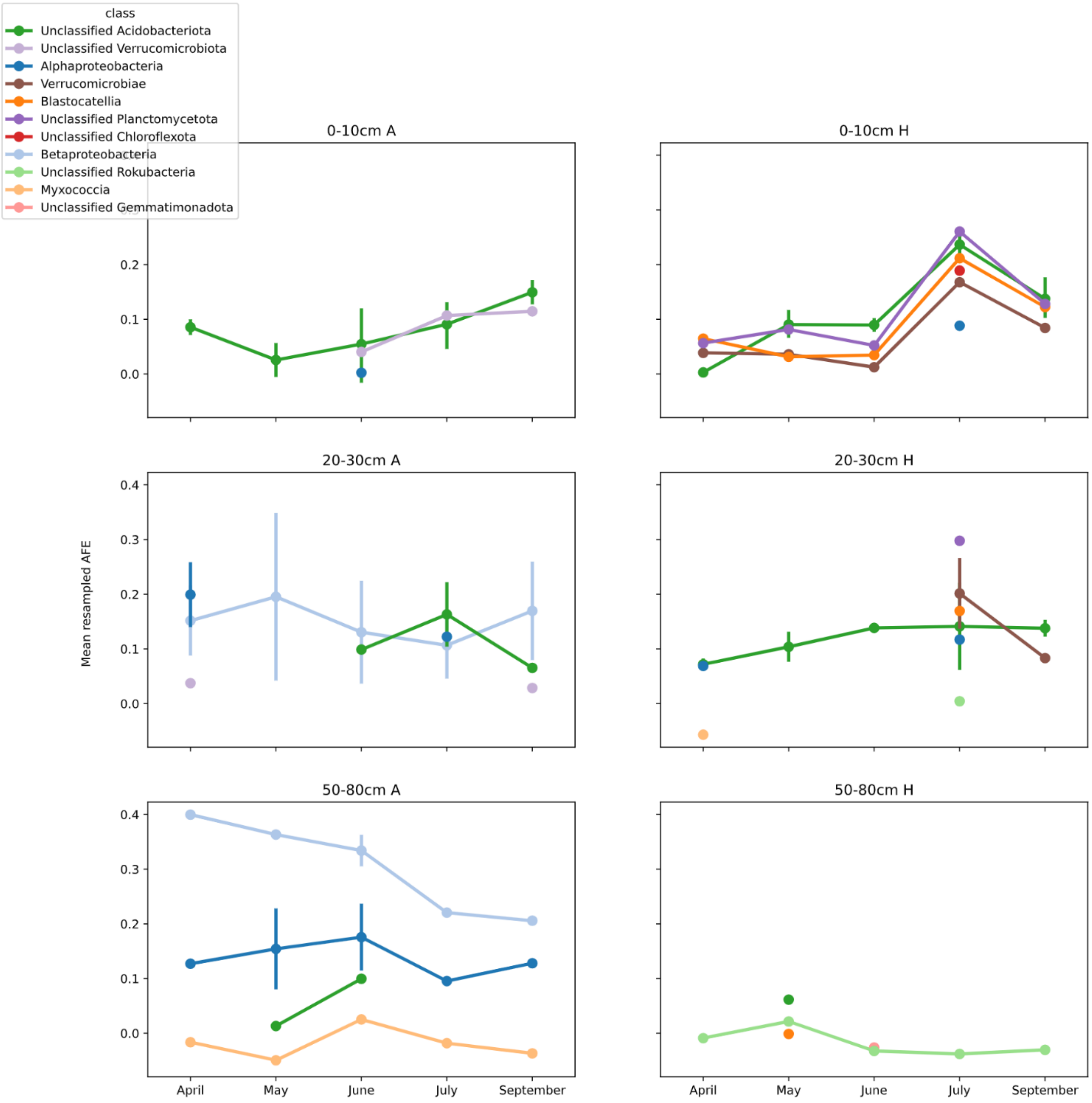


*Figure 36. The presence of genomes containing genes for the T6SS spike, tube, and sheath as function of AFE across time for each ecosystem (columns) and depth (rows). Different phylogenetic classes are indicated with colored lines.*

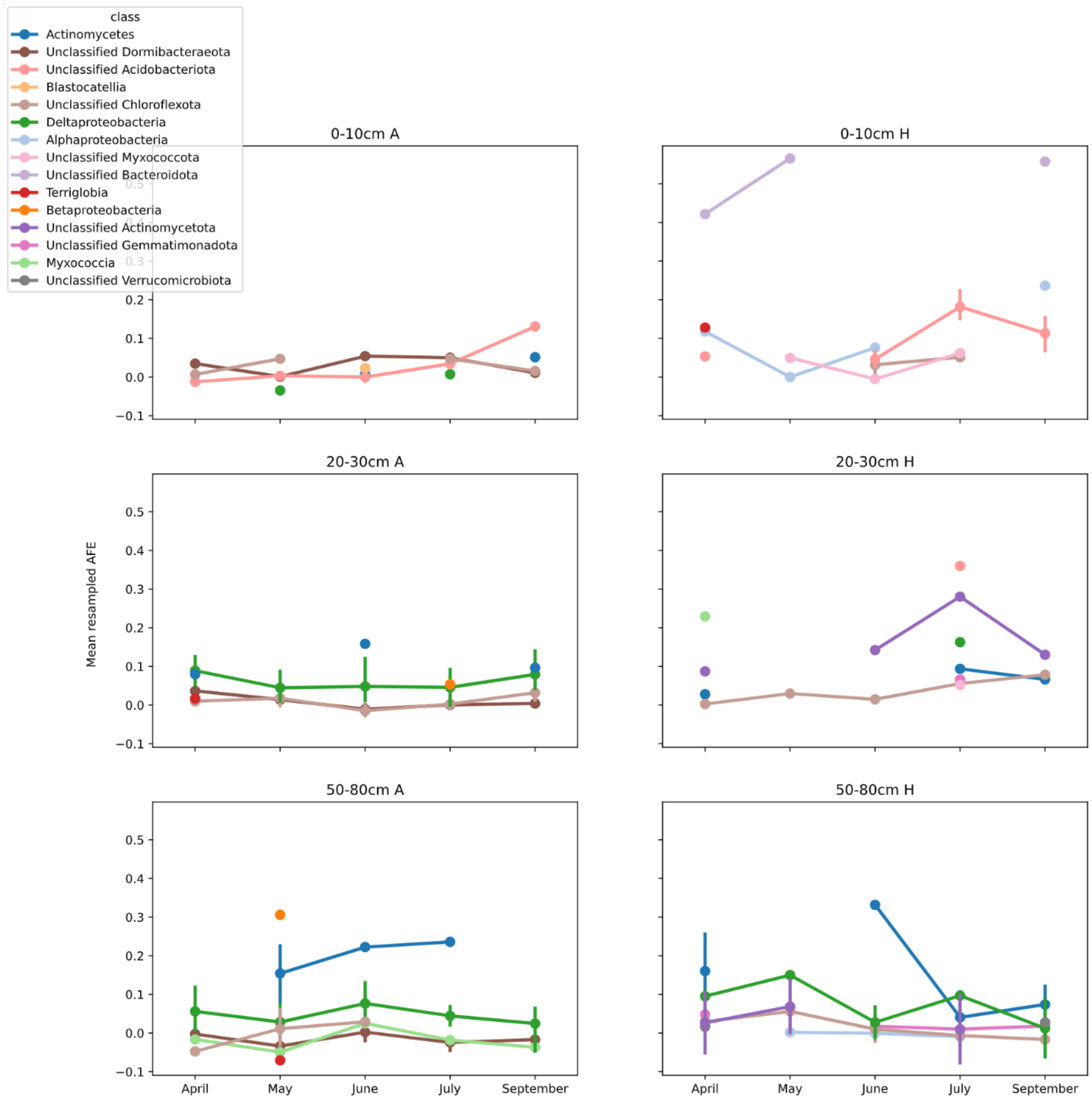


*Figure 37. The presence of genomes containing genes for the CIS spike, tube, and sheath as function of AFE across time for each ecosystem (columns) and depth (rows). Different phylogenetic classes are indicated with colored lines.*

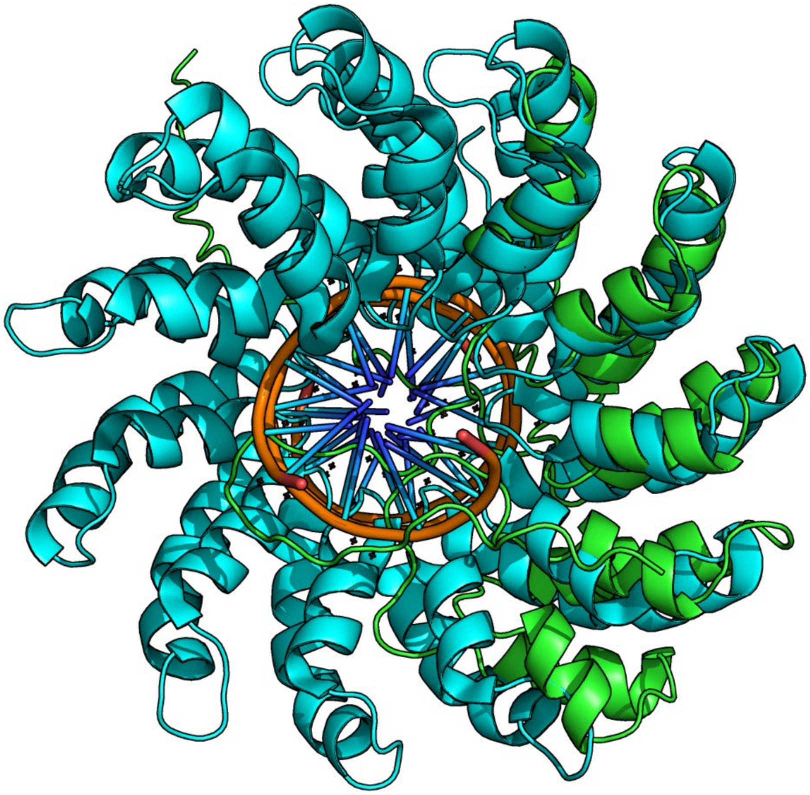

*Figure 38. Structural similarity between a Haliangiales possible effector (green) and a TAL effector repeat protein (cyan). The reference for the TAL protein is from pdb 4OSQ chains B (protein), G, and H (DNA). RMSD calculated with the cealign algorithm of PyMOL is 3.8 and the TM-Score generated by foldseek is 0.38. Foldseek ranked the hit against the reference with an E-value of 3.90e-2 and a probability score of 0.92*

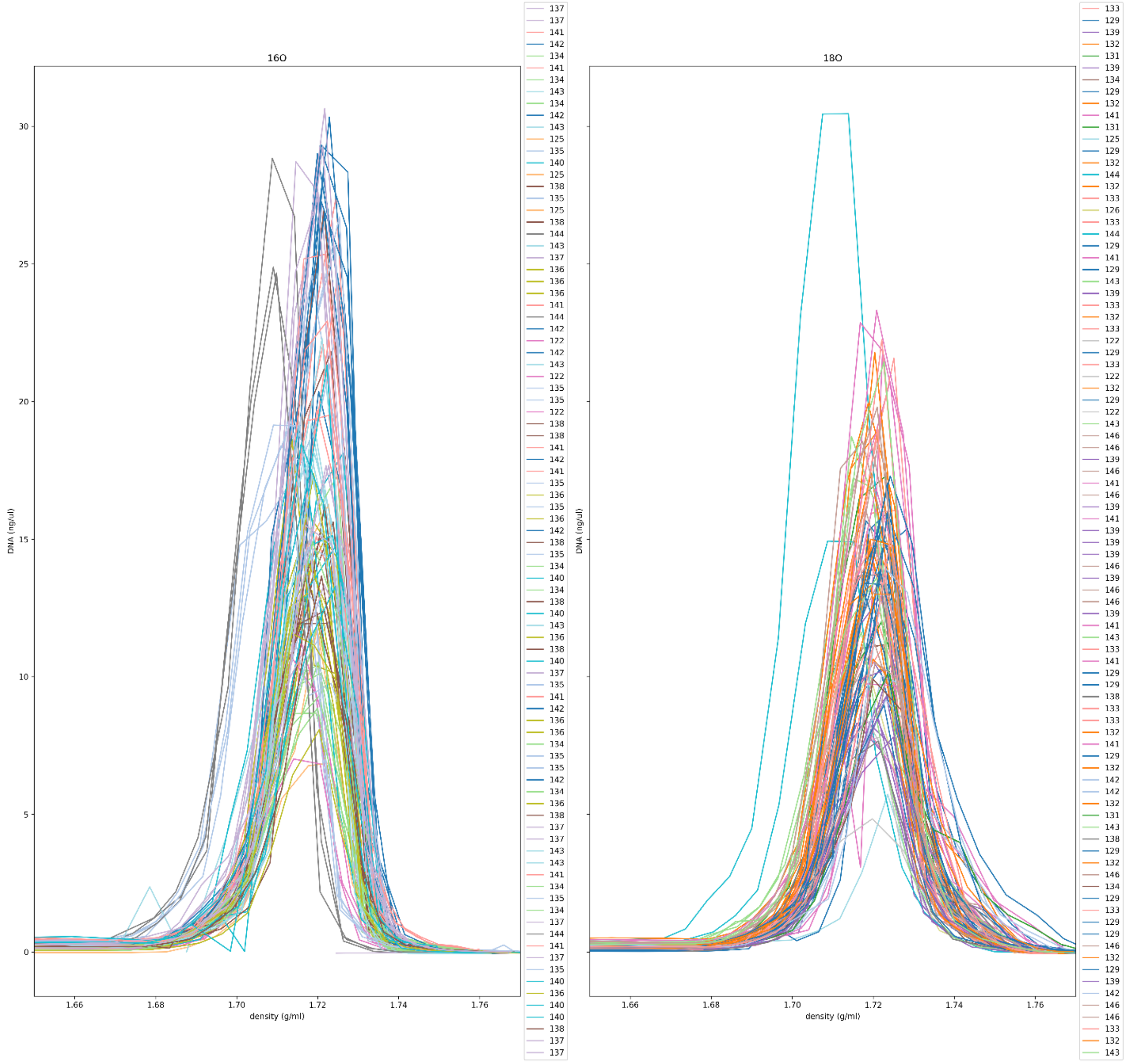


*Figure 39. Sample densities as a function of detected DNA amounts for 16O (left) and 18O (right). Each sample is represented by a colored line, lines sharing a color come from the same batch. Batches are indicated in the legend. Abnormal shifts are observed for batches 135 and 144 in the 16O samples. The 18O samples from batch 144 also show abnormally low densities.*

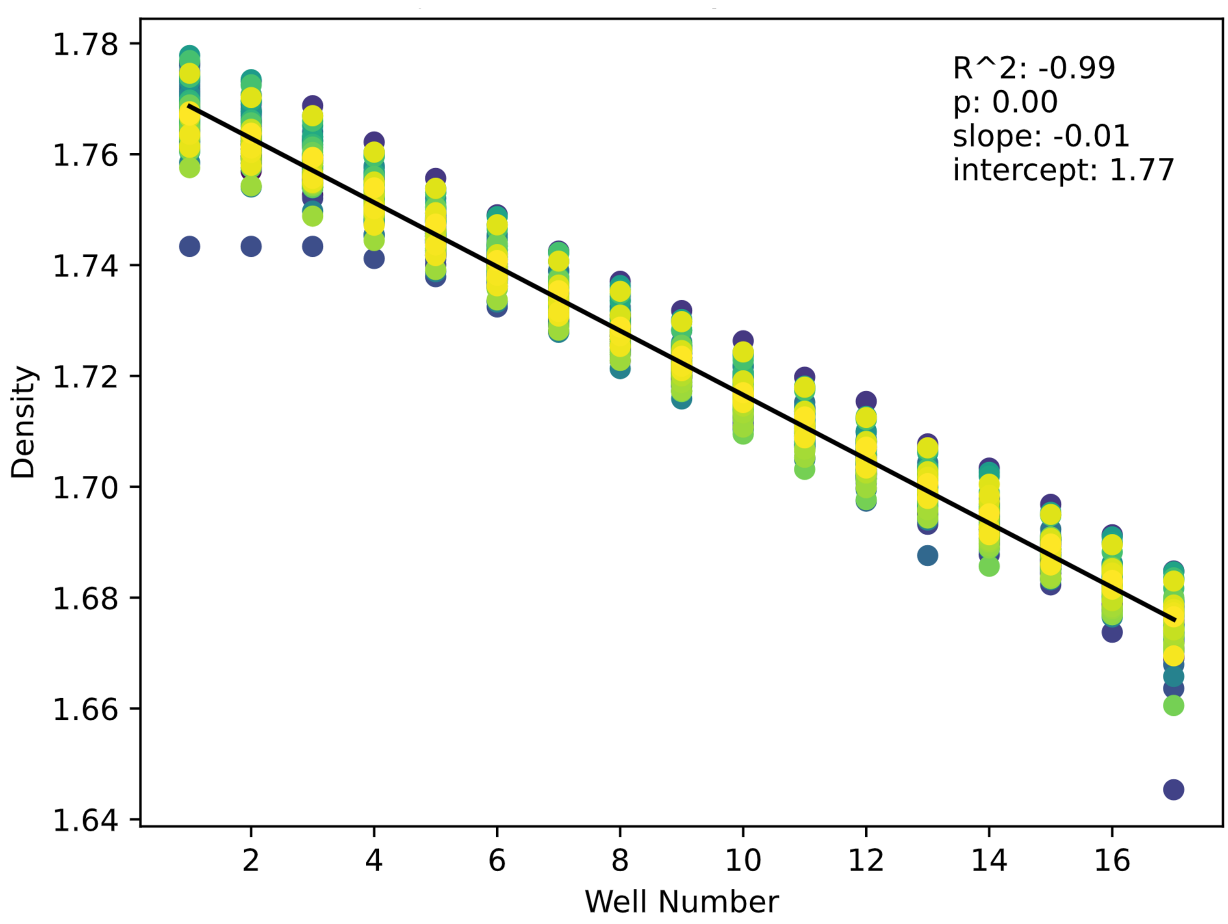


*Figure 40. Linear regression based on all 16O samples, except samples from batches which ran with abnormal densities (Fig. 39).*

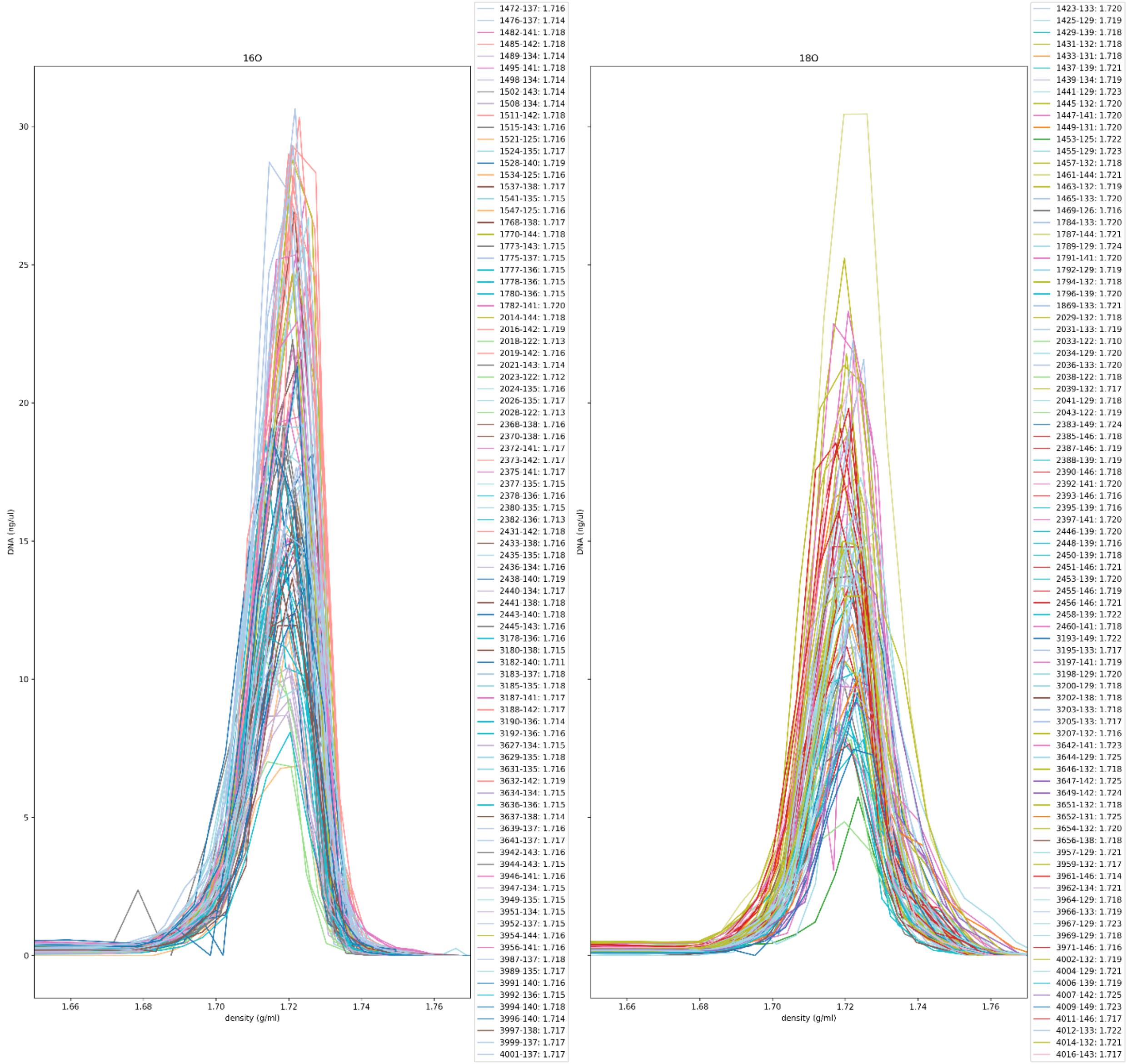


*Figure 41. Final sample densities where samples from problematic batches were adjusted based on the linear regression from figure 40. The legend shows a unique sample identifier, the batch, and the average density.*

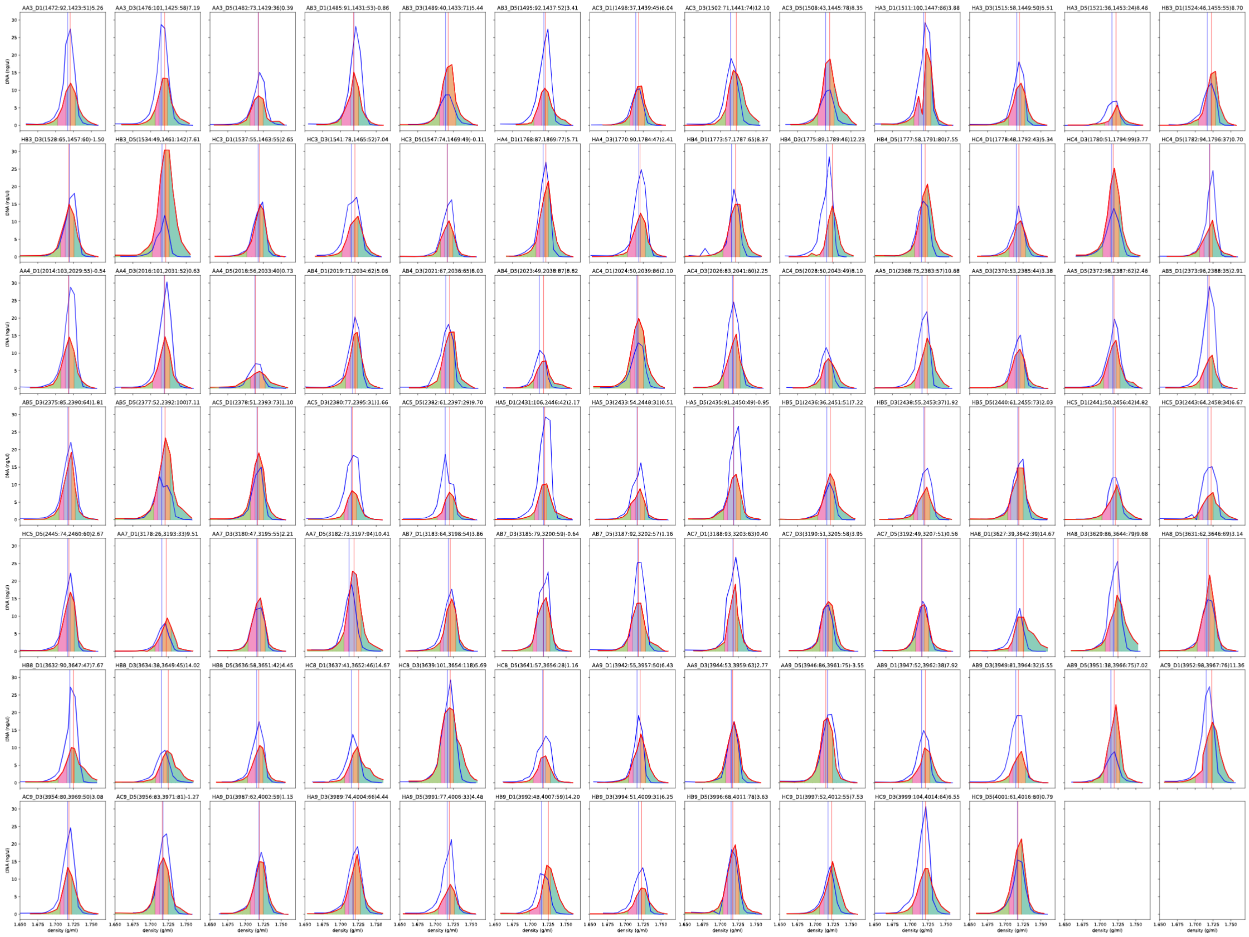

*Figure 42. Sample densities as a function of detected DNA amounts for 16O (blue) and 18O (red). Each sample is represented with its own subplot. The calculated mean average densities for each sample are represented with red (18O) or blue (16O) vertical lines. The determination of fraction bin groups for sequencing are shown with different background colors under the line for 18O.*

*Table 1. References for the used HMMs in the type six secretion system (T6SS) and extracellular contractile injection system (eCIS) analysis.*

| Name | description | System |
|---|---|---|
| **PF04965** | gp25 baseplate | T6SS |
| **PF17541** | TssC | T6SS |
| **TIGR03361** | TssI VgrG | T6SS |
| **TIGR03357** | TssE | T6SS |
| **TIGR03345** | TssH | T6SS |
| **TIGR03355** | TssC like | T6SS |
| **PF05488** | PAAR generic | T6SS |
| **PF05591** | TssB VipAshaft | T6SS |
| **PF05638** | Hcp effector | T6SS |
| **PF05936** | VasE | T6SS |
| **PF05947** | TssF inner tube | T6SS |
| **PF06744** | IcmF_C inner membrane docking | T6SS |
| **PF06812** | ImpA_N inner membrane | T6SS |
| **PF06996** | TssG Hcp tube | T6SS |
| **PF09850** | DotU inner membrane docking | T6SS |
| **PF12790** | SciN outer membrane biofilm | T6SS |
| **afp10** | PAAR-repeat motif | eCIS |
| **afp11** | baseplate J | eCIS |
| **afp12** | baseplate | eCIS |
| **afp13** | adenovirus fiber | eCIS |
| **afp14** | tape measure | eCIS |
| **afp15** | AAA+ ATPase TssH | eCIS |
| **afp16** | phage-tail terminator | eCIS |
| **afp1_5** | phage-tail tube TssD/Hcp | eCIS |
| **afp2_3_4** | phage-tail sheath TssB/TssC | eCIS |
| **afp6** | transcriptional regulator | eCIS |
| **afp7** | LysM domain TssG | eCIS |
| **afp8** | phage-tail spike VgrG | eCIS |
| **afp9** | baseplate wedge gp25 TssE | eCIS |
